# Pathogen community composition and co-infection patterns in a wild community of rodents

**DOI:** 10.1101/2020.02.09.940494

**Authors:** Jessica L. Abbate, Maxime Galan, Maria Razzauti, Tarja Sironen, Liina Voutilainen, Heikki Henttonen, Patrick Gasqui, Jean-François Cosson, Nathalie Charbonnel

**Author notes:** Corresponding author: E-mail address (N. Charbonnel).

## Abstract

Rodents are major reservoirs of pathogens that can cause disease in humans and livestock. It is therefore important to know what pathogens naturally circulate in rodent populations, and to understand the factors that may influence their distribution in the wild. Here, we describe the occurrence and distribution patterns of a range of endemic and zoonotic pathogens circulating among rodent communities in northern France. The community sample consisted of 713 rodents, including 11 host species from diverse habitats. Rodents were screened for virus exposure (hantaviruses, cowpox virus, Lymphocytic choriomeningitis virus, Tick-borne encephalitis virus) using antibody assays. Bacterial communities were characterized using 16S rRNA amplicon sequencing of splenic samples. Multiple correspondence (MCA), multiple regression and association screening (SCN) analyses were used to determine the degree to which extrinsic factors (study year and site; host habitat, species, sex and age class) contributed to pathogen community structure, and to identify patterns of associations between pathogens within hosts. We found a rich diversity of bacterial genera, with 36 known or suspected to be pathogenic. We revealed that host species is the most important determinant of pathogen community composition, and that hosts that share habitats can have very different pathogen communities. Pathogen diversity and co-infection rates also vary among host species. Aggregation of pathogens responsible for zoonotic diseases suggests that some rodent species may be more important for transmission risk than others. Moreover, we detected positive associations between several pathogens, including *Bartonella*, *Mycoplasma* species, Cowpox virus (CPXV) and hantaviruses, and these patterns were generally specific to particular host species. Altogether, our results suggest that host and pathogen specificity is the most important driver of pathogen community structure, and that interspecific pathogen-pathogen associations also depend on host species.

## 1. Introduction

Infectious diseases are among the most important global threats to biodiversity, wildlife and human health, and are associated with potential severe socioeconomic consequences (Daszak *et al*., 2000; Smith *et al*., 2006; Jones *et al*., 2008). Although combatting these risks is a main worldwide priority, our understanding of the processes underlying disease emergence still remains too limited for developing efficient prediction, prevention and management strategies. In humans, the majority of emerging pathogens originate as zoonoses from animal host populations in which they naturally circulate (Taylor *et al*., 2001; Jones *et al*., 2008). Thus, identifying the epidemiological features (e.g., prevalence, diversity, host specificity, geographic distribution) of zoonotic pathogen communities in their wild hosts, and the factors that influence pathogen occurrence in those communities, is as important to human health as it is to understanding the fundamentals of disease ecology (Garchitorena *et al*., 2017).

Both extrinsic and intrinsic factors can contribute to the composition of natural pathogen communities within and between wild animal species, populations and individuals. Factors extrinsic to the hosts include geographic location, climate, periodicity of epidemic cycles and abiotic features influencing inter-specific transmission opportunities (e.g., (Harvell *et al*., 2002; Burthe *et al*., 2006; Poulin *et al*., 2012). Factors extrinsic to the pathogens such as host species identity, sex, age, and body condition as well as genetic and immunogenetic features have also been intensively studied (e.g., (Beldomenico *et al*., 2008; Streicker *et al*., 2010, 2013; Salvador *et al*., 2011; Charbonnel *et al*., 2014; Bordes *et al*., 2017)). Although less investigated, inter-specific ecological interactions (e.g., competition, facilitation) among pathogens within animal hosts are also likely to be an important intrinsic force in determining the composition of pathogen communities. Ecological interactions between free-living species are well-known to play a part in the distribution, abundance, and many other qualitative and quantitative features of populations; the application of this basic tenant of community ecology to pathogen incidence and expression of disease has become recognized as imperative for assessing both risks and potential benefits posed to human health, agriculture, wildlife, and conservation (Pedersen & Fenton, 2007). Simultaneous infection by multiple parasite species is ubiquitous in nature (Petney & Andrews, 1998; Cox, 2001; Moutailler *et al*., 2016), and prior infections can have lasting effects on future susceptibility via e.g., changes to host condition and behavior or through immune-mediated processes (Singer, 2010; Quiñones-Parra *et al*., 2016; Kumar *et al*., 2018; Karvonen *et al*., 2019). Interactions among co-circulating parasites may have important consequences for disease severity, pathogen transmission, host and pathogen evolution or co-evolution, and community-level responses to perturbations (Jolles *et al*., 2008; Telfer *et al*., 2010; Alizon *et al*., 2013; Seppälä & Jokela, 2016; Abbate *et al*., 2018). Consequences of interaction may be life-long, as exposure to pathogens circulating among juveniles have been found to be strongly associated with those experienced by adults (Fountain-Jones *et al*., 2019). Such interactions can also play a role in the consequences of pathogen emergence (e.g., emerging bacterial infection increasing susceptibility to an endemic virus (Beechler *et al*., 2015)). Henceforth, and through the advent of sequencing technologies in particular, it is now possible and essential to investigate disease emergence from a multi-host / multi-pathogen perspective (Galan *et al*., 2016), considering the potential influence of pathogen interactions on current and future disease distributions (Cattadori *et al*., 2008; Jolles *et al*., 2008; Budischak *et al*., 2015; Abbate *et al*., 2018).

Rodent communities are relevant models for developing this community ecology approach to disease distribution and emergence. They harbor a wide variety of pathogenic taxa (Bordes *et al*., 2013; Pilosof *et al*., 2015; Koskela *et al*., 2016; Diagne *et al*., 2017) and are important reservoir hosts of agents of zoonoses that have severe implications for human health. Han et al. (2015) have revealed that about 10% of the 2277 extant rodent species are reservoirs of 66 agents of zoonoses, including viruses, bacteria, fungi, helminths, and protozoa. They also described 79 hyper-reservoir rodent species that could be infected by multiple zoonotic agents. Strong ecological interactions, such as facilitation and competition, have been shown in wild rodent populations among some of these zoonotic agents (Telfer *et al*., 2010), as well as between non-zoonotic agents and zoonotic agents (e.g., helminthes and bacteria (Carvalho-Pereira *et al*., 2019); helminthes and viruses (Guivier *et al*., 2014; Sweeny *et al*., 2020); helminthes and protozoa (Knowles *et al*., 2013)). In addition, rodents share a number of habitats with humans, including urban settings, agricultural lands, and forests, providing opportunities for human-rodent contact and pathogen transmission (Davis *et al*., 2005). Describing the distribution and composition of natural pathogen communities in rodent populations, and determining the drivers behind pathogen associations, is imperative for understanding the risks they may pose for public health.

In this study, we analyzed the pathogen communities carried by rodent communities in a rural area of northern France, a region known to be endemic for several rodent-borne diseases including nephropathia epidemica (Puumala orthohantavirus (Sauvage *et al*., 2002)) and borreliosis (Razzauti *et al*., 2015). We investigated exposure histories (via the presence of antiviral antibodies) for several viruses (hantaviruses, cowpox virus, lymphocytic choriomeningitis virus, Tick-borne encephalitis virus) and current or recent exposure to bacterial pathogens (using high-throughput 16S metabarcoding of host splenic tissue). We described the pathogens detected, their prevalence in the community and their individual distributions among host populations. We then tested the role of extrinsic factors (e.g., habitat, host species, host age) in explaining variation in pathogen distributions, and for associations (non-random co-infection frequencies) between pathogens that might indicate intrinsic drivers (e.g., competition, facilitation) of pathogen community composition. We expected that host species and habitat would be the most important factors structuring pathogen community composition because most pathogens are largely host-specific, but those sharing habitats should also share opportunities for transmission (Davies & Pedersen, 2008). After accounting for extrinsic factors, we expected to retrieve several pathogen-pathogen associations previously identified in the literature. This included i) positive associations between cowpox virus and *Bartonella* infections (*Microtus agrestis,* (Telfer *et al*., 2010)); ii) positive associations between distinct *Mycoplasma* species in mammalian hosts (Sykes *et al*., 2008; Tagawa *et al*., 2012; Fettweis *et al*., 2014; Volokhov *et al*., 2017)); iii) associations between *Bartonella* and hemotropic *Mycoplasma* species (both positive and negative associations, as well as experimental demonstration of dynamic interactions, have been described in *Gerbillus andersonii* (Eidelman *et al*., 2019)). Lastly, we also expected to find previously undescribed associations due to the large bacteria and rodent dataset included in our study. All these associations were likely to differ between host species, as differences in host specificity are also likely to be accompanied by differences in transmission dynamics and host responses to infection (Davies & Pedersen, 2008; Singer, 2010; Dallas *et al*., 2019).

## 2. Materials and methods

### 2.1. Study area and host sampling

Rodent sampling was conducted over two years (Autumn 2010 & 2011) in rural habitats surrounding two villages (Boult-aux-Bois and Briquenay) in the Ardennes region of northern France (previously described in (Gotteland *et al*., 2014)). Sex and age class (based on specific body measurements and classed as ‘adult’ for sexually mature animals and ‘juvenile’ for both juveniles and sexually immature sub-adults) were recorded for each animal, a blood sample was taken for serological analyses, and animals were then euthanized using isofluorane inhalation. Spleens were taken and stored in RNA*later* Stabilizing Solution (Invitrogen) at -20°C. Species captured from the two sites included (family: Cricetidae) 195 *Arvicola scherman* (montane water vole), 10 *Microtus agrestis* (field vole), 66 *Microtus arvalis* (common vole), 203 *Myodes glareolus* (bank vole); and (family: Muridae) 43 *Apodemus flavicollis* (yellow-necked mouse), 156 *Apodemus sylvaticus* (wood mouse), 32 *Rattus norvegicus* (brown rat). These seven focal host species were collected from traps placed in distinct landscapes (henceforth referred to as host ‘habitats’) (Supplemental Materials Figure S1): *R. norvegicus* were found uniquely on farms, *Ar. scherman* and *Mi. arvalis* were found almost entirely in meadows, and the five remaining species occupied both forests and hedgerows. Demographic differences between host species were observed for sex (e.g., male bias in *Ap. sylvaticus*; Figure S2A) and age classes (e.g., relative abundance of juveniles in *Mi. arvalis* and *Ap. sylvaticus* hosts; Figure S2B). Five *Microtus subterraneus* (European pine vole) and one each of three additional host species (one cricetid one echimyid and one murid) were also found in these communities, but excluded from analyses due to their rarity; these rare (non-focal) hosts and their pathogens are described in Supplemental Materials Appendix 1.

### 2.2. Detecting virus exposure and bacterial infection

Among the 713 rodents sampled for this study, indirect fluorescent antibody tests (IFATs; see for details (Kallio-Kokko *et al*., 2006)) were successfully performed on 677 animals to detect immunoglobulin G (IgG) specific to or cross-reacting with cowpox virus (CPXV, *Orthopoxvirus*), Puumala or Dobrava-Belgrade virus (respectively PUUV and DOBV, *Orthohantavirus*, collectively referred to henceforth as “hantavirus”), lymphocytic choriomeningitis virus (LCMV, *Mammarenavirus*), and Tick-borne encephalitis virus (TBEV, *Flavivirus*). We refer to these antiviral antibody tests as indicating a history of past exposure, but antibodies against hantavirus and LCMV also likely indicate continued chronic infection. In contrast, current or very recent exposure to bacterial infection was tested via 16S rRNA gene amplicon sequencing of splenic tissue, giving no indication of past exposure history. The spleen was chosen because this organ is known to filter microbial cells in mammals, allowing the detection of a wide array of pathogenic and zoonotic agents. Funding was available to test for bacteria in just half of the animals, chosen haphazardly to equally represent all host species, study sites and years, resulting in successful analysis for 332 rodents (see Figure 1 for a breakdown of number of individuals sampled per focal host species). For each individual animal, the DNA from splenic tissue was extracted using the DNeasy Blood & Tissue kit (Qiagen) following the manufacturer recommendations. Each DNA extraction was analyzed twice independently. We followed the method described in Galan *et al*. (2016) to perform PCR amplification, indexing, pooling, multiplexing, de-multiplexing, taxonomic identification using the SILVA SSU Ref NR 119 database as a reference (http://www.arb-silva.de/projects/ssu-ref-nr/). Briefly, DNA samples were amplified by PCR using universal primers targeting the hyper variable region V4 of the 16S rRNA gene (251 bp) and sequencing via Illumina MiSeq. The V4 region has been proven to have reasonable taxonomic resolution at the genus level (Claesson *et al*., 2010). A multiplexing strategy enabled the identification of bacterial genera in each individual sample (Kozich *et al*., 2013). Data filtering was performed as described in Galan *et al*. (2016) to determine presence/absence of bacterial infections (summarized in Figure S3). Briefly, we discarded all bacterial OTUs containing fewer than 50 reads in the entire dataset and animals for which one or both individual PCR samples produced fewer than 500 reads. A bacterial OTU was considered present in an animal if the two independent PCR samples were both above a threshold number of reads, defined as the greater of either 0.012% of the total number of reads in the run for that OTU (i.e., filtering using the rate of indexing leak) or the maximum number of reads for that OTU in any negative control sample (i.e., filtering using the presence of reads in the negative controls due to contaminations) (Galan *et al*., 2016). We removed chimera (Ashelford *et al*., 2006), using the *Uchime* program implemented in *mothur*, and manually checked OTUs representing suspected chimera not identified by the program. For each OTU suspected as pathogenic, Basic Local Alignment Search Tool (BLAST) searches of the most common sequences were conducted to infer species identity where possible. Given that rare, low abundance taxa tend to show high dissimilarity between sample replicates (Smith & Peay, 2014) we assumed that only OTUs with at least 500 reads across all animals in the dataset were considered reliably detectable, allowing us to assign absent status to these OTUs in animals failing to meet the criteria for OTU presence. Thus, only OTUs for which there were at least 500 reads across all animals in the dataset (for which present and absent statuses could be assigned), and where reasonable certainty of pathogenicity could be established from the literature (see Table 1), were considered in analyses of the pathogen community.

**Figure 1.**
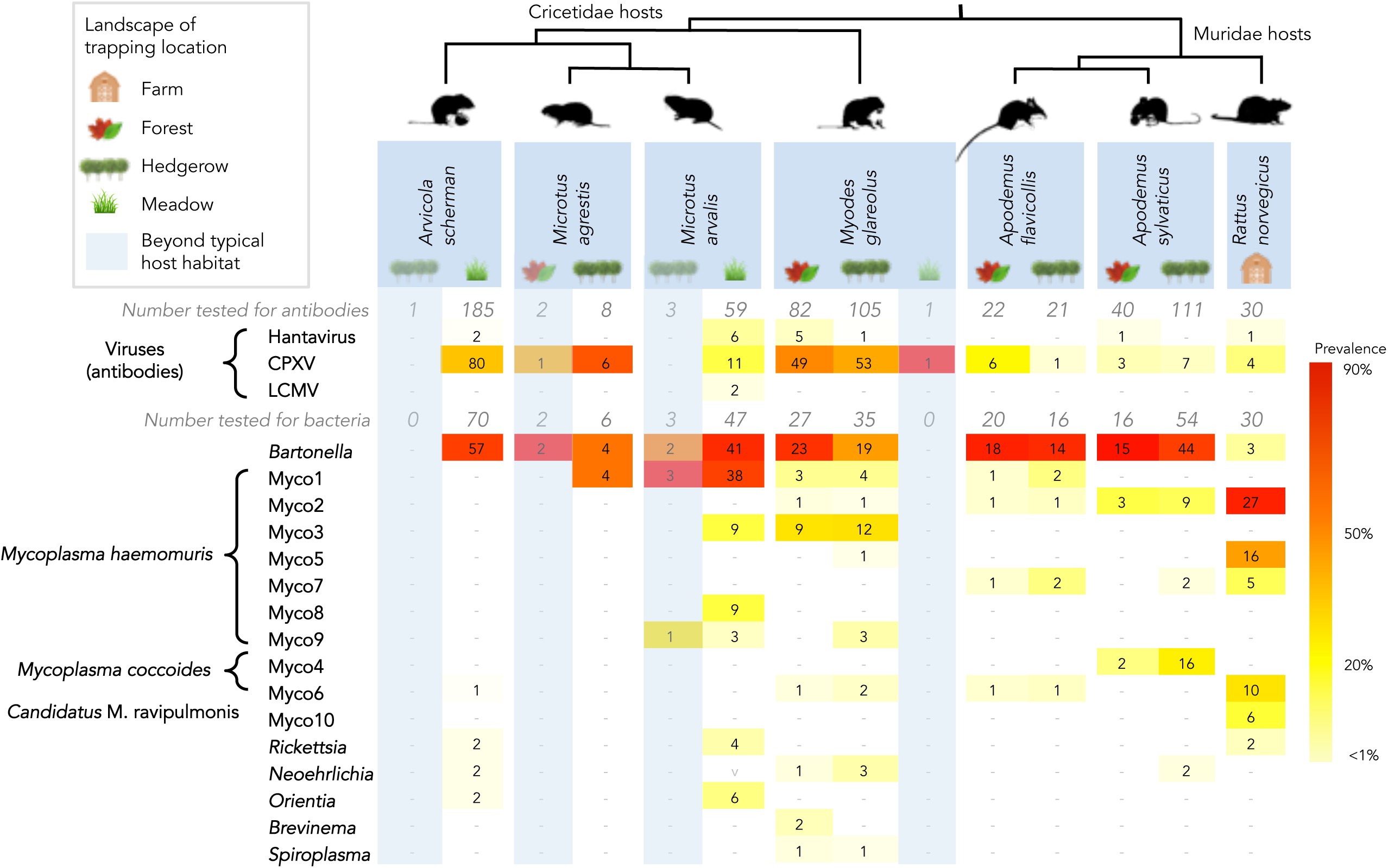
Pathogen occurrence across the rodent species community.

**Table 1.** BLAST search results for OTUs suspected of belonging to pathogenic genera. References are available in Supplemental Materials Appendix 3.

| Infesting species identity | OTU Number | Number of Reads | Genbank Accession Number | BLAST results (% identity) | Pathogen Code | Reference |
| --- | --- | --- | --- | --- | --- | --- |
| **Pathogenic taxa, reliably detectable** |  |  |  |  |  |  |
| *Bartonella* spp. | Otu00001 | 6353372 | MT027154 | 100% *Bartonella grahamii* (AB426637) from wild North America rodents; 99%-100% identity to many other pathogenic *Bartonella* species. | Bartonella | Deng et al., 2012 |
| *Brevinema* spp. | Otu00123 | 5603 | MT027155 | 97% *Brevinema andersonii* (NR_104855) type sequence, infectious spirochaete of short-tailed shrew and white-footed mouse in North America | Brevinema | Defosse et al., 1995 |
| *Candidatus Neoehrlichia mikurensis* | Otu00039 | 18358 | MT027156 | 100% *Candidatus Neoehrlichia mikurensis* (KF155504) tick-borne rodent disease, opportunistic in humans | Neoehrlichia | Andersson & Raberg, 2011 |
| *Mycoplasma ravigulmonis* | Otu00054 | 6086 | MT027164 | 100% *Mycoplasma ravigulmonis* (AF001173), grey lung disease agent in *Mus musculus*; 88% *M. orale* from humans (LR214940) | Myco10 | Petterson et al., 2000 |
| *Mycoplasma* spp. | Otu00004 | 845971 | MT027157 | 99% identity to uncultured *Mycoplasma* species (KU697344) from small rodents in Senegal and uncultured eubacterium (AJ292461) from Ixodes ticks; 95% (KM538694) and 94% (MK353834) identity to uncultured hemotropic *Mycoplasma* species in European and South American bats | Myco1 | Galan et al., 2016 |
| *Mycoplasma* spp. | Otu00003 | 426034 | MT027158 | 99% *Mycoplasma haemomuris* (AB758439) from *Rattus rattus* | Myco2 | Conrado et al., 2016 |
| *Mycoplasma* spp. | Otu00006 | 106443 | MT027159 | 99% uncultured *Mycoplasma* spp. (KP843892) from leopards in South Korea; 99% identity to uncultured *Mycoplasma* (KT215637) from rodents in Brazil | Myco3 | Goncalves et al., 2015 |
| *Mycoplasma* spp. | Otu00010 | 72724 | MT027160 | 99% *Mycoplasma coccoides* comb. nov. (AY171918); 97% *Candidatus Mycoplasma turicensis* (KJ530704) from Indian mongoose | Myco4 | Conrado et al., 2015 |
| *Mycoplasma* spp. | Otu00005 | 165095 | MT027161 | 100% *Mycoplasma haemomuris*-like undescribed species (KJ739312) from *Rattus norvegicus* | Myco5 | Alabi et al., 2020 |
| *Mycoplasma* spp. | Otu00007 | 92237 | MT027162 | 99% uncultured *Mycoplasma* a spp. (KC863983) from *Micromys minutus* (eurasian harvest mouse) in Hungary; 98% *M. coccoides* comb. nov. (AY171918) | Myco6 | Conrado et al., 2015 |
| *Mycoplasma* spp. | Otu00012 | 39767 | MT027163 | 93% uncultured *Mycoplasma* species (KU697341) of mice in Senegal; 92% *Mycoplasma* spp. (KP843892) from leopards in South Korea, 91% *Mycoplasma haemomuris* (AB820289) in rats | Myco7 | Galan et al., 2016 |
| *Mycoplasma* spp. | Otu00015 | 31528 | MT027165 | 98% uncultured *Mycoplasma* spp. (KT215632) from wild rodent spleen in Brazil; 95% uncultured *Mycoplasma* spp. (KF713538) in little brown bats | Myco8 | Ricardo Gonçalves et al., 2015 |
| *Mycoplasma* spp. | Otu00049 | 40125 | MT027166 | 96% uncultured *Mycoplasma* spp. from Brazilian rodents (KT215638) and S. Korean leopard (KP843892) | Myco9 | Ricardo Gonçalves et al., 2015 |
| *Orientia* spp. | Otu00111 | 876 | MT027167 | 97% *Orientia tsutsugamushi* (KY583502) from humans in India, zoonotic Rickettsial pathogen (causes scrub typhus) | Orientia | Paris et al., 2013 |
| *Rickettsia* spp. | Otu00008 | 72098 | MT027168 | 98% *Rickettsia japonica* (MF496166) which causes Japanese spotted fever, *R. canadensis* (NR_029155) & *R. rhipicephali* (NR_074473) type strains | Rickettsia | Yamamoto et al., 1992 |
| *Spiroplasma* spp. | Otu00093 | 4738 | MT027169 | 95% uncultured *Spiroplasma* spp. (KT983901) from Ixodes tick on a dog; 94% identity to type strain of *Spiroplasma mirum* (NR_121794), the agent of suckling mouse cataract disease; *Spiroplasma ixodetis* causes similar disease in humans. | Spiroplasma | Cisak et al., 2015 |
| **Pathogenic taxa, not reliably detectable** |  |  |  |  |  |  |
| *Arcobacter cryaerophilus* | Otu00296 | 403 | MT027170 | 100% *Arcobacter cryaerophilus* (CP032825) emerging enteropathogen in humans, zoonotic, pathogenic in rats | Arcobacter | Vandamme et al., 1992 |
| *Borrelia miyamotoi* | Otu00318 | 419 | MT027171 | 100% *Borrelia miyamotoi* (CP010308) in humans and Ixodes, zoonotic pathogen | Borrelia1 | Krause et al., 2015 |
| *Borrelia* spp. | Otu00514 | 206 | MT027172 | 96% *Borrelia* sp. nov "Lake Gaillard" in *Peromyscus leucopus* (AY536513), 95% *B. hermsii* (MF066892) from tick (*Ornithodoros hermsii*) bites in humans | Borrelia2 | Bunikis et al., 2005 |
| *Borrelia afzelii* | Otu00071 | 78 | MT027173 | 100% *Borrelia afzelii* (CP009058) human pathogen closely related (98%) to *B. burgdorferii* (positive control sequence) | Borrelia3 | Schuler et al., 2015 |
| *Leptospira* spp. | Otu01015 | 257 | MT027174 | 100% several pathogenic *Leptospira* species from mammals, e.g., *L. interrogans* (LC474514) in humans | Leptospira | Vincent et al., 2019 |
| *Mycoplasma pulmonis* | Otu00771 | 255 | MT027175 | 100% *Mycoplasma pulmonis* (NR_041744) chronic respiratory pathogen of mice and rats | Myco0771 | Piasecki et al., 2017 |
| *Mycoplasma* spp. | Otu04125 | 164 | MT027176 | 90% *Mycoplasma ravigulmonis* (AF001173), grey lung disease agent in *Mus musculus*; 80% *M. phocidae* from California sea lions (DQ521594) | Myco4125 | Petterson et al., 2000 |
| Eukaryotic family Sarcocystidae | Otu00056 | 8684 | Not submitted* | 97% similar to plastid small ribosomal unit of *Hyaloklossia lieberkuehni* (AF297120), a parasitic protazoa of European green frog; 96% *Neospora caninum* (MK770339) & *Sarcocystis muris* (AF255924); 95% *Toxoplasma gondii* (TGU28056) | Sarcocystidae1 | Calarco & Ellis, 2020 |
| Eukaryotic family Sarcocystidae | Otu00191 | 3678 | Not submitted* | 92% *Neospora caninum* (MK770339) parasite; 90% *Toxoplasma gondii* (U87145) zoonotic pathogen | Sarcocystidae2 | Calarco & Ellis, 2020 |
| Eukaryotic family Sarcocystidae | Otu00254 | 1219 | Not submitted* | 98% *Sarcocystis muris* (AF255924) coccidian parasite first found in mice | Sarcocystidae3 | Orosz, 2015 |
| * No reference specimen for identification of Eukaryote |  |  |  |  |  |  |

Table 1 (continued). BLAST search results for OTUs suspected of belonging to pathogenic genera. References are available in Supplemental Materials Appendix 3.
| Infecting species identity | OTU Number | Number of Reads | Genbank Accession Number | BLAST results (% identity) | Pathogen Code | Reference |
| --- | --- | --- | --- | --- | --- | --- |
| **Uncertain pathogenicity, reliably detectable** |  |  |  |  |  |  |
| *Corynebacterium xerosis* | Otu00050 | 1853 | MT027177 | 100% *Corynebacterium xerosis* (MH141477), only opportunistic infections identified | Corynebacterium | Bernard, 2012 |
| *Dietzia* spp. | Otu00102 | 2626 | MT027178 | 100% *Dietzia* spp. e.g., *D. aurantiaca* (MK25331); common contaminant; opportunistic in humans; thought to out-compete *Trypanosomes* | Dietzia | Kampfer et al., 2012 |
| *Helicobacter* spp. | Otu00013 | 34894 | MT027179 | 96% homology to *Helicobacter suncus* (AB006147) isolated from shrews with chronic gastritis; 95% identity to type specimen for *H. mustelae* (NR_029169) which causes gastritis in ferrets; but could be normal gut flora | Helico1 | Goto et al., 1998 |
| *Helicobacter* spp. | Otu00025 | 8702 | MT027180 | 97% similar to *Helicobacter trogonum* (AY686609) and *H. suncus* (AB006147), both enterohepatic *Helicobacter* spp. associated with intestinal diseases | Helico2 | Goto et al., 1998 |
| *Helicobacter* spp. | Otu00087 | 2303 | MT027181 | 99% identical to *Helicobacter aurati* (NR_025124.1), a pathogen of Syrian hamsters; 98% identical to *H. fennelliae* (GQ867176), a human pathogen | Helico3 | Patterson et al., 2000 |
| *Helicobacter* spp. | Otu00128 | 1178 | MT027182 | 99% *Helicobacter winthamensis* (AF363063), associated with gastroenteritis in humans; however, minor sequences were 100% identical to *H. rodentium* (AY631957) which is only associated to gastritis in rodents when coinfecting with other *Helicobacter* strains | Helico4 | Melito et al., 2001 |
| *Neisseria* spp. | Otu00612 | 780 | MT027183 | 97% uncultured *Neisseria* spp. associated with human prostatitis (HM080767) and cataracts (MG696979), but undistinguishable from environmental samples and healthy flora (e.g., JF139578) | Neisseria1 | Genbank, unpublished |
| Pasteurellaceae | Otu00129 | 1430 | MT027184 | 100% uncultured bacterium (MN095269) of mouse oral flora; 99% *Muribacter muris* (KP278064) of unknown pathogenicity, water fowl pathogen *Avibacterium gallinarum* (AF487729), and cattle respiratory disease agent *Mannheimia haemolytica* (CP017491) | Pasteurella1 | Nicklas et al., 2015 |
| Pasteurellaceae | Otu00203 | 521 | MT027185 | 99% *Aggregatibacter aphrophilus* (LR134327) and *Haemophilus parainfluenzae* (CP035368) opportunistic pathogens but otherwise part of normal flora | Pasteurella2 | Genbank, unpublished |
| *Rickettsiella* spp. | Otu00187 | 592 | MT027186 | 99%-100% identity to several endosymbionts of insects, eg. uncultured *Diplorickettsia* spp. in sand flies (KX363696), *Rickettsiella* spp. in Ixodes ticks (KP994859); 99% identity to *Rickettsiella agriotidis* (HQ640943) pathogen of wireworms | Rickettsiella | Duron et al., 2015 |
| *Streptococcus* spp. | Otu00115 | 1681 | MT027187 | 99% *Streptococcus hyointestinalis* from intestines of swine (KR819489) | Streptococcus | Genbank, unpublished |
| *Yersinia* spp. | Otu00041 | 7420 | MT027188 | A heterogeneous OTU some major sequences 100% *Yersinia* spp. and *Serratia* spp., including pathogenic zoonotic bacteria (e.g., *Y. pestis* NR_025160) and non-pathogenic endosymbionts of plants (NR_157762); some major sequences 100% *Pantoea agglomerans* (MN515098) opportunists | Yersinia | Kim et al., 2003 |
| **Uncertain pathogenicity, not reliably detectable** |  |  |  |  |  |  |
| *Fusobacterium* spp. | Otu00791 | 108 | MT027189 | 100% *Fusobacterium ulcerans* (CP028105) from tropical foot ulcers in humans; but also 100% identity with other fecal isolates of unknown pathogenicity in mammals (e.g., *F. varium* LR134390) | Fusobacterium | Genbank, unpublished |
| *Neisseria* spp. | Otu00454 | 148 | MT027190 | 98% uncultured microbiota of bat mating organs (KY300287), 98% *Simonsiella muelleri* commensal from human saliva (AF328145); 97% *Kingella kingae* (MF073277) pathogen in humans | Neisseria2 | Genbank, unpublished |
| *Treponema* spp. | Otu00235 | 348 | MT027191 | 93% uncultured rumen *Treponema* spp. (AB537611) | Treponema | Bekele et al., 2011 |
| *Williamsia* spp. | Otu00614 | 274 | MT027192 | 100% *Williamsia phyllosphaerae* (MG205541) and *Williamsia maris* (NR_024671), closely related to opportunistic pathogens in humans | Williamsia | Genbank, unpublished |

### 2.3. Statistical Methods

All statistical analyses were implemented in R version 3.2.2 (R Core Team, 2015). Throughout our analyses, we refer to simultaneous bacterial infections as “co-infection”, while analyses involving simultaneous presence of antiviral antibodies and bacterial infection are referred to as evaluating “co-exposure”. While we can only be sure that antiviral antibodies represent past exposure, we cannot rule out simultaneous viral and bacterial “co-infection”, particularly for viruses known to cause chronic infections. Likewise, detection of current (or very recent) bacterial infection, particularly for taxa known to cause chronic infections, cannot tell us how long the animal has carried the infection. Thus, we refrain from assuming sequence of infection for statistical tests unless specified by an *a priori* hypothesis from the literature.

#### 2.3.1. Testing for extrinsic drivers of pathogen community composition across the rodent community

We analyzed pathogen community composition across the whole rodent community. We use the term “pathogen community” to refer to the group of viruses and pathogenic bacteria for which we had the means to include, which was not exhaustive; thus measures of diversity are to be considered relative and not absolute. We first estimated pathogen community richness using the Shannon diversity index (alpha diversity) considering pathogenic bacterial OTUs and antiviral antibodies found in each study year, study site, habitat, host species, host sex and host age group (default options (natural logarithm Shannon index) in ‘diversity’ function from the *vegan* package). We evaluated a linear regression model using analysis of deviance (‘lm’ and ‘drop1’ functions from the basic *stats* package) to test for significance of differences in pathogen diversity due to the fixed factors listed above after first correcting for all other factors in the model (marginal error tested against the *F-*distribution). Post-hoc comparisons and correction for multiple tests were performed using function ‘TukeyHSD’ from package *stats* and ‘HSD.test’ from package *agricolae* to group factor levels that were not significantly different. An additional Akaike Information Criterion (AIC)-based model selection analysis was performed to assess any qualitative influence of spurious predictors on these comparisons using the ‘glmulti’ and ‘weightable’ function from package *glmulti*, The impact of host species diversity (Shannon diversity with Chao’s estimator correction using ‘Shannon’ in package *entropart*) on pathogen species diversity was tested for by correlation (‘cor.test’, package *stats*) across each year x site x habitat community.

We next tested for differences in pathogen community composition (beta diversity) between host species, habitats, study sites, years, age classes, and sexes by applying a permutational multivariate analysis of variance (PERMANOVA) on a Bray-Curtis dissimilarity matrix (‘adonis2’ function in the *vegan* package). To explore how intrinsic factors (pathogen-pathogen associations) contributed to the structure of the pathogen community, we used multiple correspondence analysis (MCA) to reduce variance in presence/absence of each bacterial pathogen species and antiviral antibody, implemented with the function ‘MCA’ in the *FactoMineR* package and visualization tools found in the *factoextra* package. This produces a set of quantitative and orthogonal descriptors (dimensions) describing the pathogen community composition, revealing correlated variables. With each MCA dimension as a continuous dependent response variable, we then evaluated linear regression models using analysis of deviance with post-hoc comparisons (as detailed above) to understand how the variation described by each MCA dimension was influenced by the extrinsic factors.

#### 2.3.2. Testing for associations between co-circulating pathogens

Because we identified a large number of pathogens, the number of potential association combinations to consider was excessively high, especially with regard to the relatively small number of rodents sampled. We therefore decided to test the significance only of those associations (i) clearly suggested by the community-wide MCA or (ii) previously described in the literature: positive association between *Bartonella* spp. and CPXV (Telfer *et al*. 2010), positive associations between *Mycoplasma* species (Sykes *et al*., 2008; Tagawa *et al*., 2012; Fettweis *et al*., 2014; Volokhov *et al*., 2017), and both positive (Kedem *et al*., 2014; Eidelman *et al*., 2019) and negative (Cohen *et al*., 2015) associations between *Bartonella* spp. and hemotropic *Mycoplasma* species. Given the *a priori* assumption that associations would differ between host species, we analyzed each host species separately; where evidence suggested no significant differences between host species (non-significant variation in the MCA dimension among host species or non-significant host species identity x explanatory pathogen term in logistic regressions), we pooled individuals into a single analysis to gain statistical power.

We tested the significance of each association using both association screening (SCN) analysis (Vaumourin *et al*., 2014) and multiple logistic regression analysis (GLMs, modeling the binomial ‘presence/absence’ status of each pathogen as a function of the occurrence of other pathogens) on the subset of host species in which the pathogens were found to circulate. We first performed SCN analysis, as this approach is among the most suitable for detecting pathogen associations in cross-sectional studies (Vaumourin *et al*. 2014). Briefly, given the prevalence of each pathogen species in the study population, SCN analysis generates a simulation-based 95% confidence envelope around the expected frequency of each possible combination of concurrent infection status (a total of 2^NP^ combinations, where NP = the number of pathogen species) under the null hypothesis of random pathogen associations. Observed frequencies of co-infection combinations falling above or below this envelope are considered to occur more or less frequently, respectively, than in 95% of the random simulations. Significance of the association is given as a *p*-value, calculated as the number of instances in which the simulated co-infection frequency differed (above or below the upper or lower threshold, respectively) from the observed frequency divided by the total number of simulations (Vaumourin *et al*., 2014).

The benefit of the SCN approach is a relatively high level of statistical power and the ability to identify precisely which combinations of pathogens occur outside the random expectations (Vaumourin *et al*. (2014)). However, the SCN is sensitive to heterogeneity in the data due to extrinsic factors (e.g., host specificity, or structuring in space, time, age or sex), which can both create and mask true associations. A multiple logistic regression approach was thus also systematically employed, as it has the benefit of explicitly taking into account potentially confounding extrinsic factors. Binomial exposure (presence/absence of either bacterial infection or antiviral antibodies) to a single pathogen was set as the dependent variable with exposure to the hypothetically associated pathogen(s) treated as independent explanatory variable(s) and extrinsic factors (host sex, host age, study year, study site, and where appropriate, habitat) were specified as covariates using function ‘glm’ in the *stats* package with a binomial logit link. When the multiple host species were involved, we tested an interaction term (host species identity x explanatory pathogen), and either (if *p* < 0.05) performed separate analyses for each host species or (if *p* ≥ 0.05) simply added host species identity as another covariate in the model. When there was no *a priori* assumption concerning timing of exposure (e.g., antiviral antibody presence is more likely to affect current acute bacterial infection than the reverse), the occurrence of each pathogen involved in a given association was set as the dependent variable in reciprocal GLMs. In contrast to the MCA dimensions above, a model selection step was first performed using the ‘glmulti’ and ‘weightable’ function from package *glmulti* to find the best model among top-ranking models (small-sample size corrected Akaike Information Criterion (AICc) score less than 2 + lowest AICc) that retained all explanatory pathogens of interest, in an effort to limit the appearance of associations due to the inclusion of spurious predictors. Statistical significance of the association was then assessed after first correcting for all remaining covariates in the best model using the ‘drop1’ function (likelihood ratio tests via single-term deletions compared to the full model). Despite a large number of *a priori* hypotheses, we regarded a p-value of < 0.05 as significant due to the very low number of individuals of each host species sampled. Though conceivably important, we also did not have sufficient power to test for additional interaction terms.

#### 2.3.3. Evaluating false discovery

Given the large number of significance tests performed on this single dataset, we compiled all relevant p-values (N=77) and applied a Benjamini-Hochberg correction procedure to estimate how many of the significant results may fall within the false discovery zone (using function ‘p.adjust’ in the *stats* package). Where model selection was performed, p-values were taken from full models prior to model selection. Among tests of positively correlated hypotheses (e.g., pairwise tests of intrinsic pathogen-pathogen associations N=20), only one of the p-values from the pair was included in calculating the false discovery rate (selected randomly). SCN analysis results were not included, because they were also expected to be positively correlated with the logistic regressions, and because the method inherently performs correction for multiple tests. Between hypotheses, the data were often composed of different non-overlapping subsets of varying sizes, and sample sizes varied widely with some being very small. Thus, application of this procedure likely indicates a conservative (low) estimate for how many null hypotheses should truly be rejected.

## 3. Results

### 3.1. Taxonomic identification and prevalence of pathogens

#### 3.1.1. Viral exposure

The most abundant virus detected was CPXV, with 222 (32.8%) positive sera of the 677 animals tested for anti-CPXV antibodies. It was detected in all focal host species. However, significant variation in prevalence was observed between focal host species (highly prevalent (43-70%) in *Ar. scherman*, *Mi. agrestis*, and *My. glareolus*; Figure 1; *χ^2^* = 119.5, *df* = 6, *p* < 10^-15^). Anti-hantavirus antibodies were detected in 16 animals (2.4%), and were significantly structured among host species (with exposure highest in *Mi. arvalis* (9.7%), *R. norvegicus* (3.3%) and *My. glareolus* (3.1%); Figure 1; *χ^2^* = 19.4, *df* = 6, *p* = 0. 0036). Anti-LCMV antibodies were detected in two *Mi. arvalis* individuals (Figure 1). No animals were positive for anti-TBE antibodies.

#### 3.1.2. Bacterial pathogens

Out of 952 bacterial OTUs represented by at least 50 reads in the dataset, 498 were considered positive in at least one animal after data filtering (presented in Supplemental Materials Table S1). Two OTUs (00024 & 00037) identified as *Bartonella* with low bootstrap values (74 and 92 respectively) appeared to represent chimeric sequences between the two highly amplified genera (*Bartonella* and *Mycoplasma*) in co-infected samples. Two OTUs (00009 & 00117) which were unclassified but which had a large number of reads in positive animals were also found to represent chimeric sequences between the two genera, despite high bootstrap values (100). Three additional chimeric *Mycoplasma* OTUs with under 500 reads were also excluded (OTUs 00076, 00159, and 00316). Two OTUs (00002 & 00059) were found to be redundant with OTUs Myco1 and Myco3, respectively, and two more (00134 & 00220) were chimera between Myco OTUs. These 11 OTUs were manually removed from the database, and are not included in Table S1.

We identified 43 OTUs belonging to bacterial genera with members known or thought to be pathogenic in mammals (Table 1). After BLAST queries, we found 16 of these OTUs (representing 7 distinct genera) which could be considered as reliably detectable pathogens in the focal host species (Figure 1). An additional 24 OTUs were considered potentially pathogenic but excluded from analyses because they were only observed in rare host species, because presence-absence could not be reliably established due to a low total number of reads (<500 in the dataset, e.g., *Borrelia* spp. and *Leptospira* spp.), because we could not rule out contamination by natural sources of non-pathogenic flora during dissection (e.g., *Helicobacter* spp., *Streptococcus* spp.) or by known contaminants of sequencing reagents (e.g., *Williamsia* spp.; (Salter *et al*., 2014)), or because their identity to a pathogenic species was uncertain due to insufficient genetic variation at the 16S rRNA locus (e.g., *Yersinia* spp.) (Table 1). We also identified three OTUs belonging to the eukaryotic family Sarcocystidae (98% sequence similarity to the coccidian parasite *Sarcocystis muris*); though each OTU was represented by >500 reads, there are currently no data on the reliability of this method for detection (Table 1). Individual infection status for each of these OTUs is given in Table S2.

The 16 reliably detectable pathogenic OTUs included *Bartonella* spp., 10 *Mycoplasma* spp. OTUs, *Rickettsia canadensis*, “*Candidatus* Neoehrlichia mikurensis”, *Orientia* spp., *Brevinema andersonii*, and *Spiroplasma* spp. Phylogenetic analysis including published sequences from BLAST queries revealed that the 10 *Mycoplasma* spp. OTUs belonged to three distinct species: *Myco. haemomuris* (Myco1-3,5,7-9), *Myco. coccoides* (Myco4 and Mco6), and “*Candidatus* Myco. ravipulmonis” (Myco10) (Figure S4). In general, these bacterial infections were present in all but 30 of the 332 animals tested (91.0% prevalent), and were not concentrated in any particular focal host species (*χ^2^* = 9.7, *df* = 6, *p* = 0.139). Prevalence of each pathogen in each focal host species is presented in Figure 1.

### 3.2. Extrinsic drivers of pathogen community diversity and composition within rodent community

#### 3.2.1. Analyses of pathogen diversity

We found evidence for the co-circulation of between 3 (in *Mi. agrestis*) and 12 (in *My. glareolus*) pathogen taxa per host species across the rodent community (Figure 1). Using multiple regression analysis on the Shannon diversity index, we found that marginal mean pathogen diversity differed significantly between host species (*F_5,69_* = 6.86, *p* < 10^-4^) and habitats (*F_2,69_* = 4.97, *p* = 0.0096), and it was significantly higher in adults than in juveniles (*F_1,69_* = 21.43, *p* < 10^-4^). Pathogen diversity did not, however, differ between study sites, years, or host sexes (Figure 2, Table S3). Post-hoc Tukey tests showed that after correcting for all other factors in the model, meadow habitats had higher diversity of pathogen exposure than forest habitats, and the diversity of pathogen communities in host species fell along a continuum between *My. glareolus* (high) and *Ar. scherman* (low) extremes (Figure 2, Table S3). The results were qualitatively identical when non-significant predictors were removed from the model following model selection (Figure S5). Host species diversity in each community (year x site x habitat) was positively correlated with pathogen diversity (*r* = 0.62, *t* = 2.5, df = 10, *p* = 0.032).

**Figure 2:**
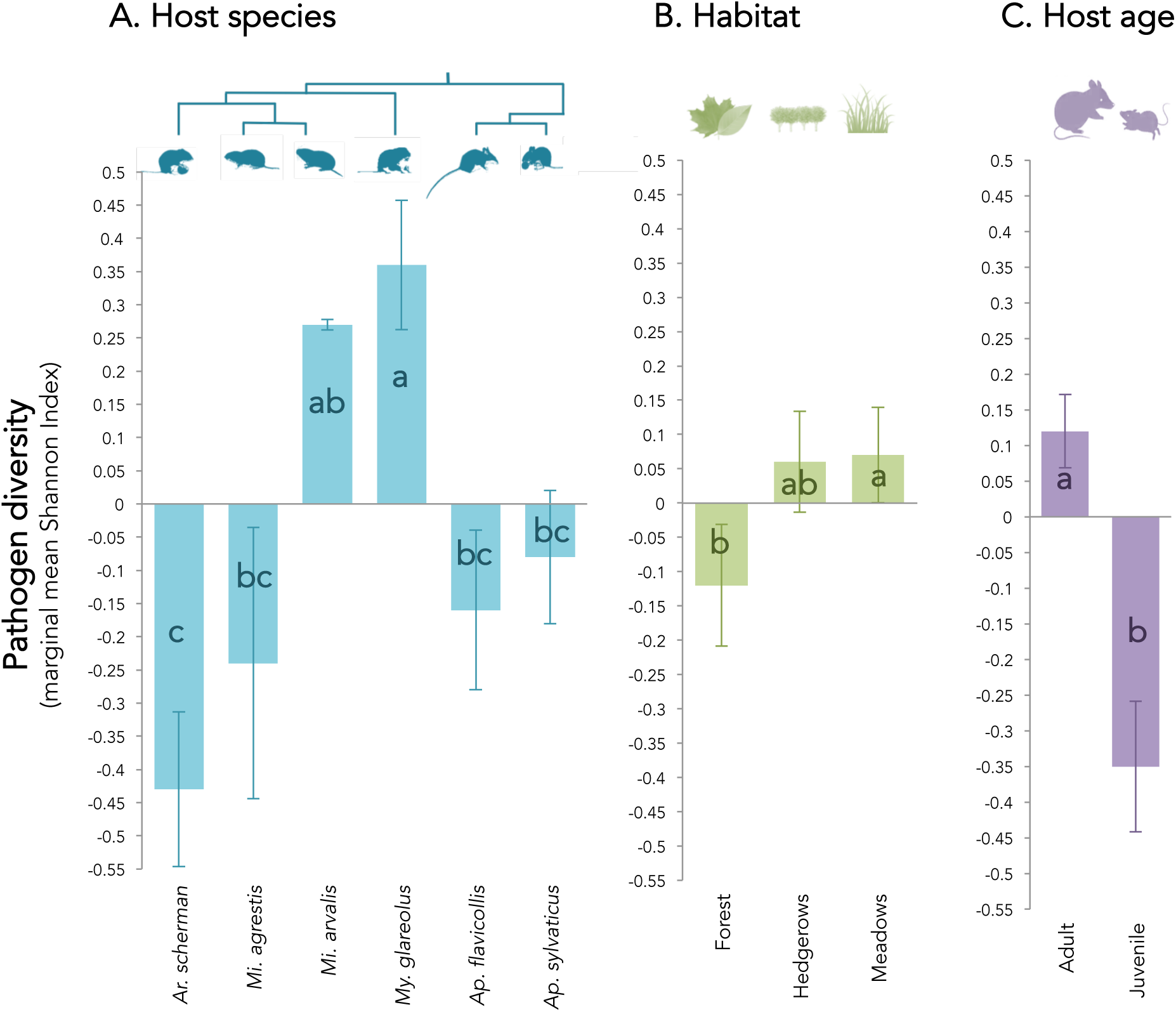
Extrinsic drivers of pathogen diversity in a rodent species community. Differences in Shannon diversity index was tested on marginal means for each factor in the multiple regression model. Different letters signify statistically significant differences at *p* < 0.05, with post-hoc Tukey adjustments for multi-level factors.

To understand the relative pathogen diversity of *R. norvegicus* hosts, not included in the above analysis because they were entirely confounded with farm habitats, we analyzed a modified model excluding host habitat and including all seven focal host species. Post-hoc Tukey tests from this model showed that *R. norvegicus* hosts had the second most diverse pathogen community (Table S4).

We also found an enormous amount of both bacterial co-infections and concurrent history of viral exposures (Figures 3A, 3B). The percentage of animals co-infected with two or more reliably detectable pathogenic bacterial OTUs among all those infected in each host species ranged between 84.4% (in *Mi. arvalis*) and 10.5% (in *Ar. scherman*). This co-infection frequency was significantly correlated with the diversity (Shannon Index) of bacteria circulating in each rodent species (Figure 3C; analysis of deviance *Pseudo-R^2^*= 0.69, *p* < 10^-11^, calculated using logistic regression weighted by the number of infected animals per species). Bacterial co-infections were more frequent than expected in *Mi. arvalis*, and less frequent than expected in *My. glareolus* (according to Cook’s Distance, Figure S6A). Results were similar when co-occurrence of antiviral antibodies was considered along with bacterial OTU exposure (Figure 3D; *Pseudo-R^2^* = 0.65, *p* < 10^-7^). While *Mi. arvalis* had both more bacterial co-infections and slightly more pathogen co-exposures than expected based on pathogen diversity, other outliers differed between the two measures (Figure S6B): *My. glareolus* co-exposure frequencies were not lower than expected, and both *Apodemus* species had lower than expected co-exposures. Host species diversity in each community did not correlate with bacterial co-infection (*r* = 0.33, *t* = 1.11, df = 10, *p* = 0.29) or pathogen co-exposure (*r* = 0.022, *t* = 0.071, df = 10, *p* = 0.95) frequencies.

**Figure 3:**
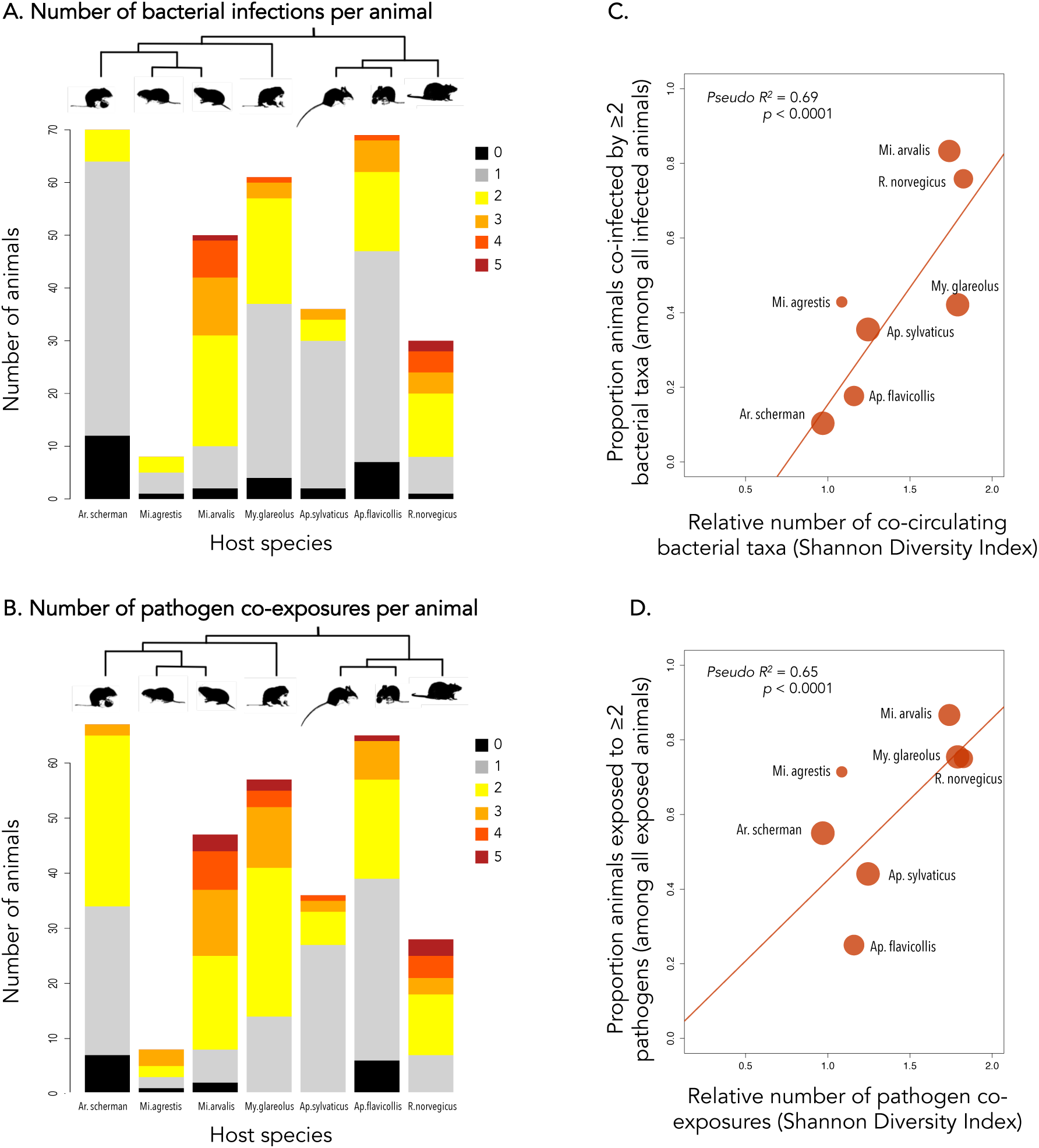
Bacterial co-infection and co-exposure patterns among host species.

#### 3.2.2. Analyses of pathogen community composition

Many pathogen taxa were found only in a single host species (*Mycoplasma haemomuris* OTU Myco8, “*Candidatus* Mycoplasma ravipulmonis*”* (Myco10), *Brevinema* spp., *Spiroplasma* spp., LCMV), and each host species had a unique combination of co-circulating pathogens (Figure 1). In order to best identify extrinsic and intrinsic factors potentially driving the composition of pathogen communities within the rodent community, we reduced the dataset to limit biases. We excluded *R. norvegicus* individuals due to competing *a priori* hypotheses that host species and habitat would be important factors (as this host species was the only one found in farm habitats, confounding these two variables; but see MCA results when *R. norvegicus* was included in Figures S7, S8). Likewise, we excluded pathogens that occurred only in one habitat type of one host species (not including *R. norvegicus*). Two additional individuals were excluded due to missing sex and age information. Analyses were performed on the remaining 280 individuals from six host species and their 14 pathogens (*Bartonella*, Myco1, Myco2, Myco3, Myco4, Myco6, Myco7, Myco9, *Rickettsia*, *Neoehrlichia*, *Orientia*, *Spiroplasma*, and antibodies against CPXV and hantaviruses) (Figure 4).

**Figure 4:**
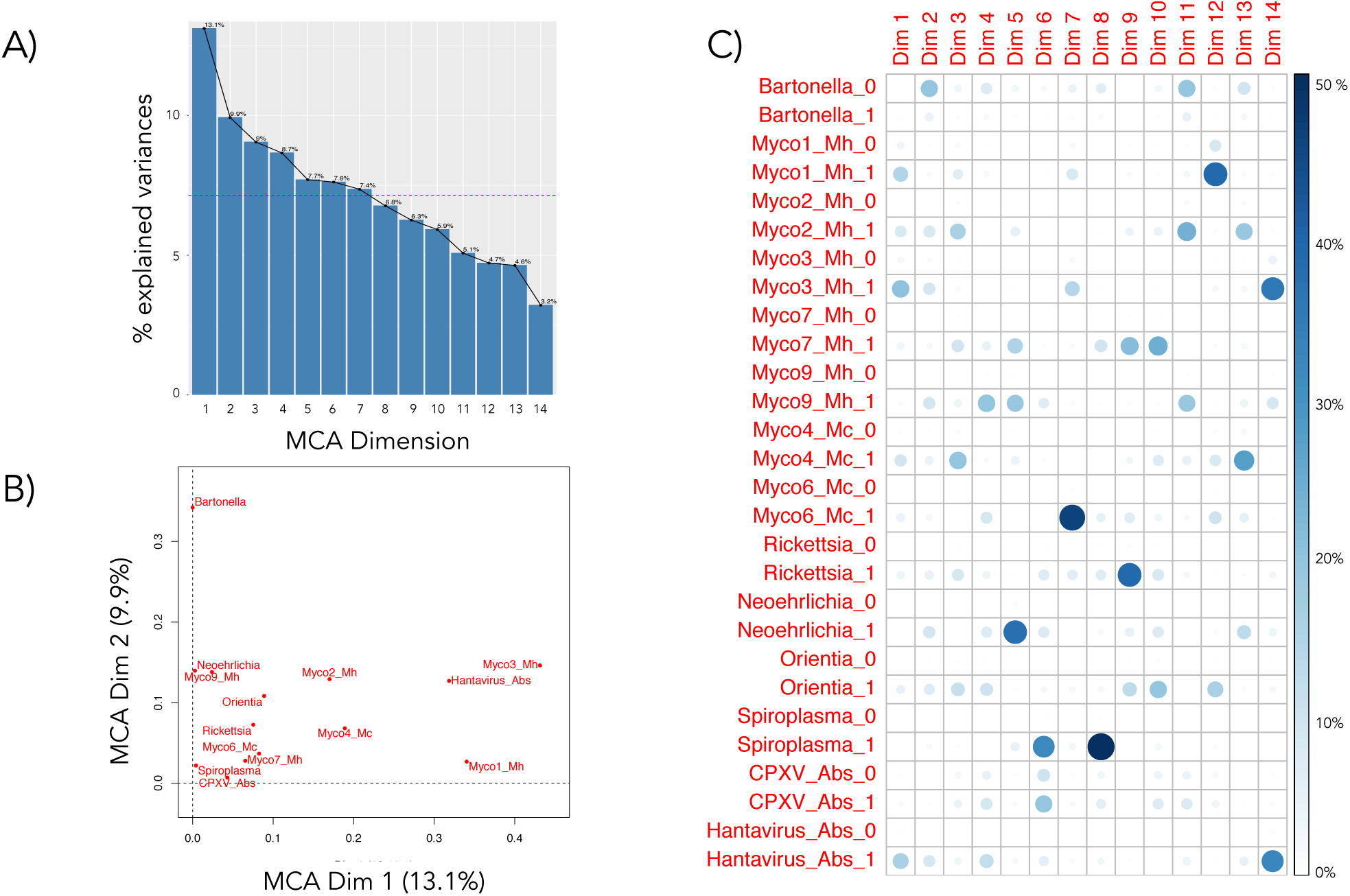
Results of multiple correspondence analysis (MCA) for pathogen community composition in rodents (excluding *R. norvegicus*) are described by (A) the contribution of each dimension to the overall variance in the data, (B) variable correlations with the first two dimensions of the MCA, and (C) variable contributions to each orthogonal MCA dimension. Horizontal line in (A) represents the percent variance expected due to chance (100/14) = 7.14%.

Overall, pathogen species composition was significantly structured by host species identity (*F_5,252_* = 16.23, p = 0.001) and habitat type (*F_2,252_* = 2.51, *p* = 0.024; Table S5). Out of 14 orthogonal dimensions returned by the MCA, the first two captured 23.0% of the variation in pathogen and antibody occurrence, and the first seven explained a cumulative 63.4% of the total variance (Figure 4A). Further dimensions captured less variance than would be expected if all dimensions contributed equally to overall inertia in the data. Dimension 1 (MCA Dim1; explaining 13.1% of the variation in pathogen community and loading heavily with the presence of Myco1, Myco3 and anti-hantavirus antibodies) differed significantly between host species (*F_5,268_* = 23.83, *p* < 0.0001) and age classes (*F_1,268_* = 6.27, *p* = 0.013; Table S6). Dimension 2 (MCA Dim2; explaining 9.9% of the variation in pathogen community and primarily describing the occurrence of *Bartonella*) was also structured significantly by host species (*F_5,268_* = 3.89, *p* = 0.002; Table S6). While these first two dimensions varied by host species (Figure S10A) and host age class (Figure S10B), variance in host habitats (Figure S10C) was not significant after accounting for the other factors. Host species was the most consistently important extrinsic driver of pathogen community composition, significantly explaining variation captured in six of the first seven dimensions, MCA Dim1 – MCA Dim7 (except for MCA Dim 5; Table S6).

### 3.3. Associations between pathogens

#### 3.3.1. Validation of the associations detected by MCA

We applied SCN and GLM analyses to further characterize patterns detected using MCA. Strong and relatively equal loading of MCA Dim1 with Myco1, Myco3, and anti-hantavirus antibody presence indicated that these three pathogens were positively associated with one-another. Indeed, the six animals with anti-hantavirus antibodies were found exclusively in animals infected with Myco3, and Myco1 was found in 2/3 of hantavirus-exposed animals but in just 1/3 of those without hantavirus exposure. The MCA also revealed significant differences among host species and host age classes for Dim 1; hantavirus and Myco3 only circulated in two host species (*Mi. arvalis* and *My. glareolus*) and 34 of those 35 occurrences were in adults. To exclude positive associations arising from mutual host specificity and age-related accumulation of exposure probability, we focused our analyses on the dataset restricted to adults of the two host species in which all three pathogens co-circulated (*Mi. arvalis* and *My. glareolus*). Individual SCN analyses performed on adults of each host species revealed no associations (Table S7A). However, since values of MCA Dim 1 did not differ between the two host species (according to post-hoc tests given in Table S6), we also ran a single SCN analysis on the pooled data from adults of both species to improve statistical power (Table S7A). According to this pooled SCN analysis, the three-way co-occurrence of Myco1, Myco3 and anti-hantavirus antibodies was significantly more frequent than would be expected by random chance (*p* = 0.008), with a trend for anti-hantavirus antibodies occurring by themselves more rarely than expected (sitting on the lower bound at zero; *p* = 0.13; Table S7A, Figure S11). We also investigated this association using GLM. However, given the small number of hantavirus exposures and perfect association with Myco3 infection, there was insufficient statistical power to explicitly test for an association between all three pathogens and extrinsic factors. We therefore ran three reciprocal GLM models on the restricted dataset, one for each pathogen as a function of extrinsic factors to control for heterogeneous host groups (host species, host sex, study site, habitat, and year sampled) and exposure to the two other pathogens (Table S7B). The number of extrinsic variables in each model was reduced using model selection (results in Figure S12). These models showed that there remained significant unexplained positive associations between hantavirus exposure and Myco1 infection (anti-hantavirus antibodies ∼ Myco1: *χ^2^* = 5.80, p = 0.016) and between hantavirus exposure and Myco3 infection (Myco3 ∼ anti-hantavirus antibodies: *χ^2^* = 13.66, *p* < 0.001), but that there was no evidence of direct association between Myco1 and Myco3 infections (Myco1 ∼ Myco3: *χ^2^* = 0.01, *p* = 0.94; Myco3 ∼ Myco1: *χ^2^*< 0.37, *p* = 0.54; Figure 5A).

**Figure 5:**
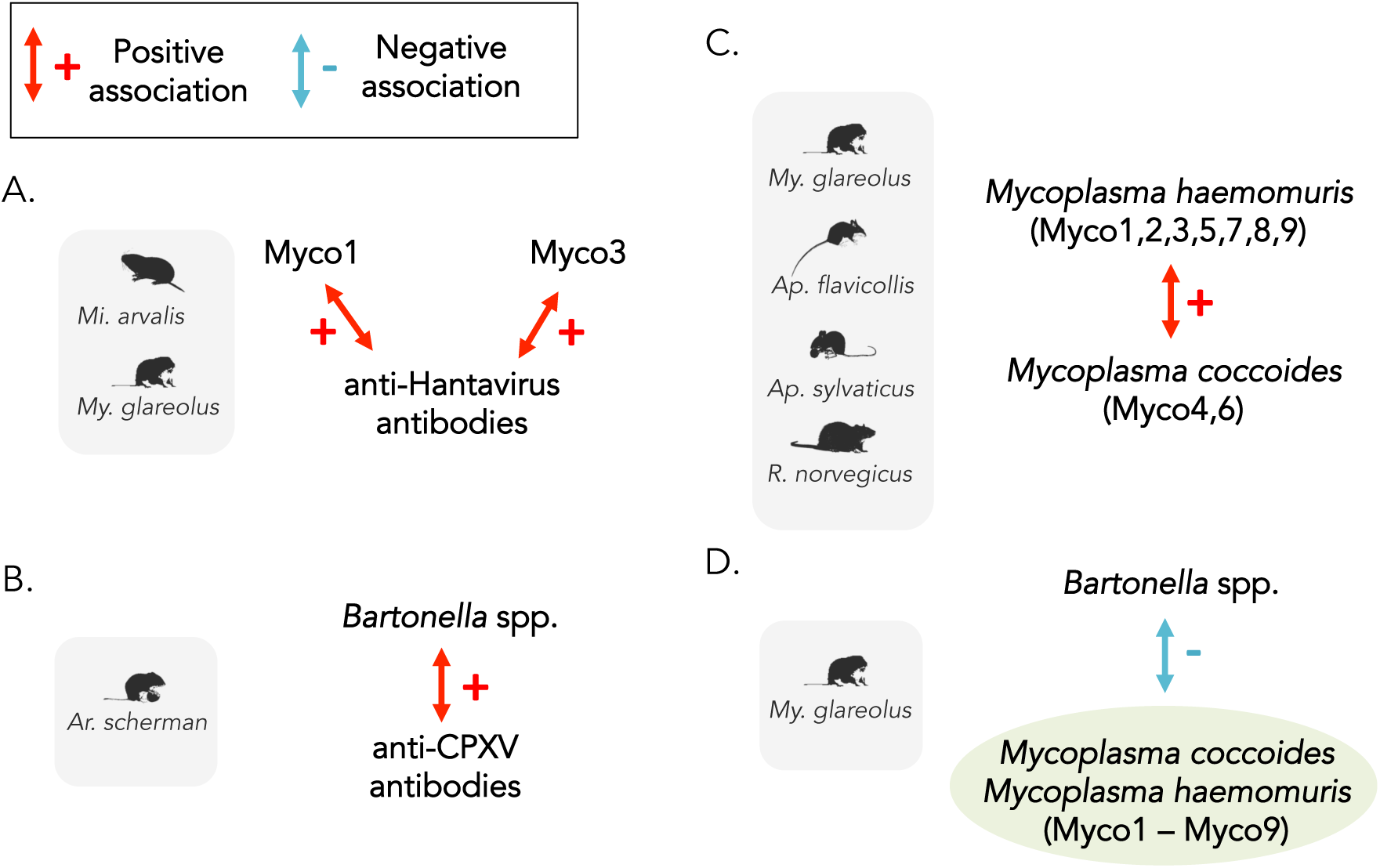
Associations between pathogens in a community of rodents. Association hypotheses were generated by multiple correspondence analysis (A) or previously noted in the literature (B, C, D). Only associations supported by significant statistical tests (*p* < 0.05) are illustrated. Red arrows represent positive associations, blue arrows represent negative associations.

The third MCA dimension also presented a clear hypothesis with sufficient statistical power to be tested. MCA Dim3 was characterized by co-variation in Myco2 (*Myco. haemomuris*) and Myco4 (*Myco. coccoides*) infections suggesting a positive association between members of these two *Mycoplasma* species. Myco2 and Myco4 OTUs co-circulated only in *Ap. sylvaticus* hosts, thus we limited our analysis to this host species. There was no significant association between the two OTUs detected by SCN analysis (Table S8A), and after correcting for extrinsic factors remaining in the models after model selection (Figure S13), there remained only a non-significant trend (Myco2 ∼ Myco4: *χ^2^* = 2.79, p = 0.12; Myco4 ∼ Myco2: *χ^2^* = 2.34, *p* = 0.13; Table S8B) for a positive association between the two OTUs. No additional associations with sufficient variance for statistical tests were clearly suggested by the MCA analysis.

#### 3.3.2. Validation of associations described in the literature

We tested the *a priori* hypothesis that seropositivity to CPXV would be positively associated with *Bartonella* infection, previously detected in *Mi. agrestis* (Telfer *et al*., 2010). The whole dataset was considered as these two pathogens co-circulated in all host species (Figure 1). SCN analyses performed independently for each host species revealed no associations (Table S9A), and the MCA results suggested that pooling data across host species would be inappropriate. After correcting for extrinsic factors using GLM, we found reciprocal evidence for a positive association in *Ar. scherman* hosts (*Bartonella ∼* anti-CPXV antibodies: χ^2^ = 5.07, *p* = 0.024; anti-CPXV antibodies ∼ *Bartonella*: χ^2^ = 5.07, *p* = 0.024; Figure 5B), but not in any other host species (Figure S14; Table S9B). It is of note that there were only eight *Mi. agrestis* individuals, rendering statistical power to test for the association while controlling for extrinsic factors insufficient in this host species where the association was previously described. While prevalence of both pathogens in *Mi. agrestis* was relatively high compared to other host species, one of the two animals without *Bartonella* infection was positive for anti-CPXV antibodies, also precluding evidence for a within-species trend.

We next focused on the potential associations between OTUs identified as belonging to two different species of hemotropic *Mycoplasma, Myco. haemomuris* (HM) and *Myco. coccoides* (HC), within the four host species in which they both circulated (*My. glareolus*, *Ap. flavicollis*, *Ap. sylvaticus*, *R. norvegicus*; Figure 1; Figure S4). We found no significant associations using independent SCN analyses for each host species (Table S10A). However, after controlling for extrinsic factors using GLM, a significant positive association was detected (HM ∼ HC: χ^2^ = 9.5, *p* = 0.0021; HC ∼ HM: χ^2^ = 9.59, *p* = 0.002), and did not differ between host species (non-significant interaction term between host species by explanatory pathogen occurrence in each reciprocal model, Figure S15; Table S10B; Figure 5C). We note that only one *R. norvegicus* animal was uninfected with *Myco. haemomuris*, and that animal also had no *Myco. coccoides* infection; thus the trend for the association in this host species was also positive but lacked sufficient variance for independent statistical analysis.

Finally, we tested for associations between *Bartonella* spp. and hemotropic *Mycoplasma* species, grouping the occurrence of different OTUs of the latter (Myco1 – Myco9) into a single presence-absence variable. There was no association detected by SCN analyses (Table S11A), and marginal evidence that any association may differ by host species after correcting for extrinsic factors using GLM with model selection (Figure S16; Table S11B). After controlling for extrinsic factors using independent GLMs for each host species (where possible), we found a negative association between the two pathogen groups only in *My. glareolus* hosts (*Bartonella* ∼ *Mycoplasma*: χ^2^ = 4.14, *p* = 0.042; *Mycoplasma* ∼ *Bartonella*: χ^2^ = 6.59, *p* = 0.010, Table S11B, Figure 5D).

### 3.4. Evaluating false discovery

Benjamini-Hochberg correction of p-values from hypothesis tests throughout the study suggested that those above ∼0.01 may lie above the false discovery cutoff for statistical significance (Figure S17), and that null hypotheses rejected with smaller p-values have been rejected with confidence.

## 4. Discussion

Rodents have long been recognized as important reservoirs of infectious agents, with a high transmission potential to humans and domestic animals (Kruse *et al*., 2004). Europe is identified as a hotspot of rodent reservoir diversity and one third of rodent species are considered hyper-reservoirs, carrying up to 11 zoonotic agents (Han *et al*., 2015). Nevertheless, associations between these pathogens have still only rarely been investigated (but see, for example, studies from field voles in the UK (Telfer *et al*., 2010) and in Poland (Pawelczyk *et al*., 2004), gerbils in Israel (Cohen *et al*., 2015), across a rodent community in North America (Dallas *et al*., 2019), and co-infection frequencies of zoonotic pathogens from rodents in Croatia (Tadin *et al*., 2012)).

In this study, we confirmed that rodent communities in northern France may harbor a large diversity of potential zoonotic pathogens, with at least 10 bacterial genera and antibodies against at least four genera of viruses. Some of these pathogens have already been reported in the study region or in geographic proximity, including viruses (*Orthohantavirus*, *Orthopoxvirus*, *Mammarenavirus* (Charbonnel *et al*., 2008; Salvador *et al*., 2011)) and bacteria (e.g., *Bartonella, Mycoplasma, Rickettsia, “Candidatus* Neoehrlichia*”, Orientia, Spiroplasma, Treponema, Leptospira, Borrelia, Neisseria, Pasteurella*; see (Vayssier-Taussat *et al*., 2012; Razzauti *et al*., 2015)). A previously undetected relative of the putatively pathogenic spirochaete *Brevinema andersonii* that infects short-tailed shrews and white-footed mice in North America (Defosse *et al*., 1995) was among our findings, and TBEV is not known to circulate this far east (Lindquist & Vapalahti, 2008). The high prevalence of anti-hantavirus antibodies in *Mi. arvalis* is likely explained by cross-reactivity between the anti-PUUV antibodies used in our assay and those elicited against the related *Tula orthohantavirus* (TULA) virus common to European voles (Deter *et al*., 2007; Tegshduuren *et al*., 2010).

Three zoonotic pathogens were particularly prevalent: *Orthopoxvirus*, *Bartonella* spp., and *Mycoplasma* spp. The wide range of hosts with anti-*Orthopoxvirus* antibodies corroborates prior evidence that cowpox virus could be widespread in European rodents, particularly voles (Bennett *et al*., 1997; Essbauer *et al*., 2010; Forbes *et al*., 2014). An astounding 77% of all individuals in the study were infected by *Bartonella* spp., a diverse group of hemotrophs known to commonly infect rodents and other mammals (Breitschwerdt & Kordick, 2000; Bai *et al*., 2009) and which have also been implicated in both zoonotic and human-specific disease (Iralu *et al*., 2006; Breitschwerdt, 2014; Vayssier-Taussat *et al*., 2016). We could not assess the specific diversity of *Bartonella* spp. circulating in these rodent communities because accurate resolution in this genus requires additional genetic markers (Matar *et al*., 1999; Guy *et al*., 2013). Hemotropic and pneumotropic *Mycoplasma* spp. were also highly prevalent across all host species, though surprisingly nearly absent from *Ar. scherman* despite expectations from related *Ar. terrestris* (Villette *et al*., 2017). These *Mycoplasma* species are also known pathogens of humans and rodents (Harwick *et al*., 1972; Baker, 1998). Here, we found two distinct hemotropic *Mycoplasma* species (*Myco. haemomuris* and *Myco. coccoides*) and the pneumotropic *Mycoplasma* species *Myco. pulmonis* and “*Candidatus* Myco. ravipulmonis”. The former two are both hemotropic mycoplasmas responsible for vector-transmitted infectious anaemia of wild mice, rats, and other rodent species (Neimark *et al*., 2001, 2005; Messick, 2004). In contrast, *Myco. pulmonis* and “*Candidatus* Myco. ravipulmonis” cause respiratory infections, are more closely related to other pneumotropic mycoplasmas, and “*Candidatus* Myco. ravipulmonis” has only ever before been described in laboratory mice (formerly termed Grey Lung virus (Andrews & Glover, 1945; Neimark *et al*., 1998; Graham & Schoeb, 2011; Piasecki *et al*., 2017)).

Our results also corroborated the status of hyper-reservoir (more than two zoonotic pathogens carried by a reservoir species) for all seven of the focal rodent species studied here (Han *et al*., 2015). Even the rare host species *Mi. subterraneous* also carried two potentially zoonotic pathogens (*Bartonella* spp. and *Brevinema* spp.; Appendix 1). Overall, we found a high variability in the number of pathogens circulating in each species despite correction for sampling effort, with low levels observed in *Apodemus* species and *Arvicola scherman*, and high levels detected in *Mi. arvalis, My. glareolus*, and *R. norvegicus*. While physiology, genetics, and behavior can contribute to the number of pathogen species able to infect a given host species, larger geographic range size is highly correlated with higher pathogen species diversity (Morand, 2015); this explanation matches the pattern among hosts in the communities sampled here (i.e., *Ar. scherman* and *Apodemus* spp. have small geographic ranges compared to those of *Mi. arvalis*, My*. glareolus*, and *R. norvegicus*).

Several studies have emphasized the influence of host habitat specialization on parasite species richness, low habitat specialization being associated with both high species richness of macro- and micro-parasites (e.g., (Morand & Bordes, 2015)). Our results did not fully corroborate this association; while the grassland-specific *Ar. scherman* had the lowest pathogen diversity and the multi-habitat spanning *My. glareolus* had the highest pathogen diversity, entirely farm-dwelling *R. norvegicus* had high pathogen diversity nearly equal to that of *My. glareolus*, and the two *Apodemus* hosts (neither with significantly higher pathogen diversity than *Ar. scherman*) were found across both meadows and hedgerows. Instead, we found that more diverse host species communities hosted more diverse pathogen communities. However, the implications of that result are complex because while exposure to diverse (i.e., potentially novel) pathogens is a risk for disease emergence, diverse host species communities are thought to keep individual pathogen prevalence low due to the dilution effect – which should limit risk of zoonoses (Keesing *et al*., 2010).

The search for factors that drive parasite species richness, diversity and community composition has been at the core of numerous studies (Poulin, 1995; Poulin & Morand, 2000; Nunn *et al*., 2003; Mouillot *et al*., 2005; Krasnov *et al*., 2010; Sallinen *et al*., 2020). Here, we emphasized that both pathogen diversity and community composition was mainly structured by host species identity, despite both shared habitats and shared pathogen taxa. Pathogen beta diversity was also structured by habitat, which could result from particular environmental suitability (e.g., for vectors) or opportunities for cross-species transmission. We found no evidence that any specific pathogen-pathogen associations were likely to be as important as host species identity in determining pathogen distributions across the community of rodents. The strong influence of host characteristics (Cohen *et al*., 2015) and host species identity (Dallas *et al*., 2019) on pathogen community composition has recently been described in comparison to intrinsic pathogen-pathogen associations in other rodents. Moreover, the pathogen community composition provided a unique signature for each rodent species, even among those most closely related (e.g., *Ap. flavicolis* and *Ap. sylvaticus*). This result is in line with the conclusions of meta-analyses showing that phylogeny, over other host traits, has a minimal impact on pathogen diversity in rodent species (Luis *et al*., 2013; Guy *et al*., 2019).

The importance of host species identity in shaping pathogen community composition may not stem from strict host-pathogen specificity, as most pathogens were found to infect multiple host species – a broad result echoed across animal communities (Cleaveland *et al*., 2001; Taylor *et al*., 2001; Woolhouse *et al*., 2001; Pedersen *et al*., 2005; Streicker *et al*., 2013). However, we might be cautious as more precise molecular analyses are necessary to test whether different species of a bacteria genus or divergent populations of the same bacteria species may circulate independently in different rodent host species, with little or no transmission. For example, two genera seemed to be largely shared among the rodent species studied here, *Bartonella* and *Mycoplasma*. But previous studies have shown strong host-specificity when considering the genetic variants of *Bartonella* (Buffet *et al*., 2013; Withenshaw *et al*., 2016; Brook *et al*., 2017). Evidence in the literature for host specificity of *Mycoplasma* species has led to a mix of conclusions (Pitcher & Nicholas, 2005), as cases of cross-species transmission are commonly reported – particularly in humans – despite a general consensus that most species are highly host-specific. We found that some *Mycoplasma* taxa were dominant contributors to prevalence in a single host species, and that when shared, they were shared with just a few other specific host species. Rare infections in unexpected host species (e.g., Myco6 in *Ar. scherman* and Myco1 in *Ap. flavicollis*) were represented by fewer sequence reads compared to positive samples in host species where they were more prevalent, suggesting a low potential for amplification and sustained transmission from these occasional hosts (Figure S4). On the other hand, while the Cricetidae appeared to be susceptible only – with rare exception – to taxa within the *Myco. haemomuris* group, host species in the Muridae family were susceptible to all three distinct *Mycoplasma* species detected. The biggest exception to this pattern was that three of 62 *Myodes glareolus* (sister to all other sampled Cricetidae in the study) animals were found to be infected by both hemotropic *Mycoplasma* species. These results both support the observation that cross-species transmission naturally occurs among wild rodents and suggest that the degree of host specificity may be driven by both host and pathogen factors.

Concurrent exposure to multiple pathogens within individuals was also frequent, as high as 89 % (in *Mi. arvalis* hosts), in line with recent studies that have shown that co-infections by multiple pathogens are common in natural populations (e.g., in mammals, birds, amphibians, ticks, humans (Telfer *et al*., 2010; Griffiths *et al*., 2011; Clark *et al*., 2016; Moutailler *et al*., 2016; Stutz *et al*., 2018)). Variation in the frequency of pathogen co-exposure was highly correlated to the diversity of pathogens circulating in each host species, suggesting the dominance of a random process of pathogen exposure for each individual. However, there were a few intriguing outliers: *My. glareolus* hosts were less co-infected than expected based on diversity of bacterial taxa, but not when viral antibodies were included; conversely, *Ar. scherman* hosts were more co-exposed when viruses were considered, but not when only bacteria were considered; and *Mi. arvalis* hosts had consistently higher proportions of co-exposures whether viruses were or were not considered along with bacteria. The non-random grouping of pathogen exposures within individuals (as in *Mi. arvalis*) may result from heterogeneity in extrinsic transmission, environmental, or susceptibility factors (Cattadori *et al*., 2006; Swanson *et al*., 2006; Beldomenico *et al*., 2008; Beldomenico & Begon, 2010; Fenton *et al*., 2010) or from intrinsic interactions between pathogens (e.g., facilitation mediated by hosts immune response). Differences in the pattern of co-exposure frequencies when including or excluding antiviral antibodies (as with *My. glareolus* and *Ar. scherman*) could result from different mechanisms (e.g., bacterial manipulation of innate immunity (Diacovich & Gorvel, 2010)) affecting pathogen community assemblage. However, a lack of deviance from the expected co-exposure frequency does not exclude the possibility that both extrinsic and intrinsic processes may be occurring.

We found evidence in support of three previously identified pathogen-pathogen associations (positive association between *Myco. haemomuris* and *Myco. coccoides* infections; positive association between *Bartonella* spp. infection and the presence of anti-CPXV antibodies; negative association between *Bartonella* spp. and hemotropic *Mycoplasma* spp. infections) and characterized one set of associations not previously described (positive associations between the presence of anti-hantavirus antibodies and infections by two specific *Myco. haemomuris* OTUs) – each in a unique subset of host species. *Mycoplasma* spp. blood infections are likely transmitted through bites of blood-sucking arthropod vectors (Volokhov *et al*., 2017), meaning vectors could prefer some individuals over others (Malmqvist *et al*., 2004). Positive associations detected between *Myco. haemomuris* and *Myco. coccoides* could also result from similarities in rodent susceptibility. Indeed *Mycoplasma* spp. infection can lead to acute or chronic infection, and the establishment of chronic bacteremia seems to occur in immunosuppressed or immunocompromised individuals (Cohen *et al*., 2018). Co-infections with multiple *Mycoplasma* spp. might therefore be more likely to be detected in these immunocompromised rodents with chronic infections. The existence of chronic infections might also lead to additional co-infections and positive associations as a result of disease-induced changes in population dynamics, immune system function, or through direct pathogen-pathogen interactions (Fenton, 2008; Aivelo & Norberg, 2018; Fountain-Jones *et al*., 2019).

Whether through the accumulation of exposure probabilities or increased susceptibility, the previously-undocumented positive association we found here between *Myco. haemomuris* OTUs (Myco1 and Myco3) and anti-hantavirus antibodies may similarly be explained by the chronic nature of both *Mycoplasma* spp. and hantavirus infections in rodents (e.g., for Puumala hantavirus in bank voles (Yanagihara *et al*., 1985; Meyer & Schmaljohn, 2000; Vaheri *et al*., 2013)). This positive association was found in both host species where the majority of hantavirus exposures occurred (*Microtus arvalis* and *Myodes glareolus*), consistent with the generality of association between *Mycoplasma* species across host taxa detailed above, suggesting the intrinsic ecology of these pathogens contributes to shaping variation in the pathogen community. Curiously, we found no evidence for direct associations between OTUs of the same *Mycoplasma* species, thus facilitation interactions are unlikely to explain the high diversity of *Mycoplasma* taxa both within and between host species.

Infections by *Bartonella* species are also known to often result in subclinical and persistent bacteremia in mammals, including rodents (Birtles *et al*., 2001; Kosoy *et al*., 2004). The positive association detected in *Ar. scherman* between *Bartonella* spp. and anti-CPXV antibodies might therefore be explained by, for example, joint accumulation of both chronic bacterial infections and long-lived antiviral antibodies used to test for prior exposure to relatively short-lived CPXV infections. However, if the same processes governing association of the chronic infections described above were at play here, we would have expected to find both pathogens implicated in positive associations (i) with other chronic infections, and (ii) across host species given their ubiquitous prevalence. While the failure to recover the association in *Mi. agrestis* (previously described (Telfer *et al*., 2010)) was likely due to low statistical power, the lack of a general pattern in other host species despite adequate sampling suggests a more specific, and potentially immune-mediated, ecological process between these two pathogens. Indeed, pox virus infections, including CPXV, have been shown to induce immunomodulation that increases host susceptibility to other parasites (Johnston & McFadden, 2003). These interactions could be of variable intensities according to the rodent species considered, due to potential differences in impacts of CPXV infection on immunity among host species, or to the influence of other infections not examined here on host immune responses during pox infections (e.g., helminths (Cattadori *et al*., 2007), protozoa (Telfer *et al*., 2010)). Furthermore, *Bartonella* spp. infection was negatively associated with *Mycoplasma* spp. infections in *My. glareolus*, corroborating negative interactions reported in co-infection experiments in gerbils (Eidelman *et al*., 2019). This association may therefore originate from an interaction mediated by specific (immune) genetic features of *My. glareolus*, and not ecological conditions as proposed by Eidelman et al. (2019). The antagonistic and host-specific nature of this association lends further support to the interpretation that *Bartonella* spp. infections do not behave in ways similar to other chronic infections in the community. However, few studies have investigated the robustness of within-host interactions across different host species (e.g., (Lello *et al*., 2018)), and this question deserves further investigation.

Our results suggest that intrinsic ecological interactions could help shape the composition of the pathogen community within hosts. However, this suggestion provides only a hypothesis that requires further investigation. Interpretation of associations can be misleading, as they may arise from unmeasured co-factors such as exposure to shared transmission routes, and may even run counter to the underlying ecological process (Fenton *et al*., 2014). The associations we found here were not visible (or even misleading, in the case of a 3-way interaction between hantavirus, Myco1 and Myco3), for instance, when ignoring extrinsic factors using the SCN analysis, despite the increased statistical power it offered. Our decision to screen only the spleen means we could have missed evidence of exposure to pathogens that can only be found by screening the liver, kidney or brain (Mangombi *et al*., 2021). Evidence for interactions between pathogens within hosts initially came from laboratory studies (e.g., in the development of vaccines, reviewed in (Casadevall & Pirofski, 2000)), and until recently, many studies conducted in the wild could not detect such interactions (e.g., (Behnke, 2008)). Developments in statistical approaches have contributed to improve sampling designs and analyses, in particular by better controlling for confounding factors, enabling the detection of associations resulting from these within-host interactions (e.g., (Lello *et al*., 2004; Telfer *et al*., 2010; Galen *et al*., 2019)). A structural equation modeling approach, for instance, could be applied to identify potential influence of unmeasured (latent) variables (Carver *et al*., 2015). Experiments conducted in semi-controlled environments have been used to confirm the importance of interactions suggested by associations (e.g., (Knowles *et al*., 2013)). Both facilitation mediated by immune responses (e.g., (Ezenwa *et al*., 2010)) and competition mediated by shared resources (e.g., (Brown, 1986; Budischak *et al*., 2018)) have been emphasized.

There remain additional important limits to the interpretation of snapshot observational studies from wild populations such as ours. For instance, they cannot provide information about the sequence or duration of infection, although these features strongly affect the outcome of within-host interactions (Eidelman *et al*., 2019). Moreover, both the 16S metabarcoding approach and serological antibody tests can only be interpreted in terms of presence/absence of exposure to pathogens, although co-infection may rather impact parasite abundance (e.g., (Thumbi *et al*., 2013; Gorsich *et al*., 2014)). Other extrinsic factors, such as seasonal variation in pathogen community composition, could also impact both interpretation and year-round generality of our results due to adherence to autumn sampling dates (Maurice *et al*., 2015; Villette *et al*., 2020). Lastly, we also acknowledge several caveats to consider with our methods. We removed animals from which fewer than 500 reads were amplified in one or both bacterial metabarcoding PCR replicates. While 16 of these samples removed were due to random failure of PCR amplification from just one of the two replicates, 12 of the animals had poor amplification in both PCR replicates. In the absence of an internal positive control, e.g., a spike-in standard (Zemb *et al*., 2020), we were unable to verify whether a lack of reads was due to poor DNA extraction or a true lack of infections. Although this has a risk of artificially inflating prevalence rates by selectively removing uninfected individuals, it is unlikely to have had a qualitative effect on our results. Similarly, limiting our analyses to OTUs with 500 reads or more in the entire dataset may select against detection of very rare or low-burden infections. We also removed many OTUs corresponding to bacteria normally occurring in external or internal microbiomes of healthy animals, some of which were represented by a high abundance of reads in positive animals. This was due mainly to the fact that 16S data cannot often distinguish between pathogenic and commensal taxa of many such genera. We know that, for instance, *Helicobacter* species are naturally found in the digestive tract, but can also cause pathogenic infections. Parasitism can affect host microbiome composition (Gaulke *et al*., 2019), and this in turn can have impacts on host health and disease susceptibility (reviewed in (Murall *et al*., 2017; Rosshart *et al*., 2017)). Thus, our choice to ignore OTUs corresponding to microbes typical of healthy flora contributes to the problem of missing data, such as information on intestinal helminth infections or other viruses, which may explain or alter the associations we were able to detect. Furthermore, the evaluation of diversity measures (e.g., Shannon diversity index) based only on a selection of taxa violates the assumption that all species are represented in the sample; thus, patterns of diversity could also be influenced by missing data. These caveats are common problems for disease surveillance and community ecology studies, irrespective of the diagnostic methods, and it is difficult to speculate about their overall impacts on the present study. Finally, it is well-understood that this bias towards detection of common pathogens and difficulty in interpreting evidence for the absence of a pathogen in a given individual or population can make testing for negative associations driven by antagonistic ecological interactions incredibly difficult, if not impossible (Weiss *et al*., 2016; Cougoul *et al*., 2019).

## Conclusions

Our results add to a growing number of studies finding that (i) rodents host many important zoonotic human pathogens and (ii) pathogen communities are shaped primarily by host species identity. We also detected a number of previously undescribed associations among pathogens within these rodent communities, and we also confirmed previously identified associations, sometimes in other rodent species than those in which they were previously described. These associations can be considered in the future as hypotheses for pathogen-pathogen interactions within rodent hosts, and that participate in shaping the community of pathogens in rodent communities. Long-term survey and experimental studies are now required to confirm these interactions and understand the mechanisms underlying the patterns of co-infection detected. In addition to these biological results, we have identified several methodological caveats, with regard to both pathogen and association detection, that deserve further investigation to improve our ability to make robust inference of pathogen interactions.

## Acknowledgements

We would like to thank all collaborators and research assistants that previously helped with the field work (Yannick Chaval, Cécile Gotteland, Marie-Lazarine Poulle). This study was partly funded by the INRA Metaprogram MEM Hantagulumic, and by the European programme FP7-261504 EDENext. The manuscript is registered with the EDENext Steering Committee as EDENext409. None of the rodent species investigated here has protected status (see list of the International Union for Conservation of Nature). All procedures and methods were carried out in accordance with relevant regulations and official guidelines from the American Society of Mammalogists. All protocols presented here were realized with prior explicit agreement from relevant institutional committee (Centre de Biologie pour la Gestion des Populations (CBGP): 34 169 003). We thank Hélène Vignes of the Cirad Genotyping platform for her assistance with the MiSeq sequencing, Sylvain Piry, Alexandre Dehne-Garcia and Marie Pagès for their help with the bioinformatic analysis, and Gael Kergoat for helpful comments and proofreading. We also thank the reviewers and recommenders at PCI Ecology for their careful reviews, insights, and patience. Version 4 of this preprint has been peer-reviewed and recommended by Peer Community In Ecology (PCI Ecology) (https://doi.org/10.24072/pci.ecology.100071).

## Data Accessibility

Supplementary data deposited in Zenodo (https://doi.org/10.5281/zenodo.7092812) include the following 16S metabarcoding data: (i) raw sequence reads (fastq format), (ii) raw output files generated by the mothur program (iii) raw abundance table and (iv) filtered occurrence table, as well as (v) scripts and data files for statistical analyses. Items iii-v are also provided in Supplemental Materials Appendix 2 to directly accompany this publication.

