## Supplementary material for "Pathogen community composition and co-infection patterns in a wild community of rodents": Table S1

Supplementary Materials Table S1. Catalogue of bacterial genera from splenic tissue in all rodents sampled. Table includes only genera deemed positive in at least one animal. OTU = operational taxanomic unit ; NC = negative controls ; Bold genera include OTUs deemed potentially pathogenic ; % reads in positive animals = Nb reads - NCs - Nb reads in animals not meeting criteria for positive infection (specified in the main text).

| Phylum | Class | (sub-)Order | Family | Genus | Minimum Bootstrap | Biology | Nb of OTUs | Total Nb Reads (incl. NCs) | % reads in NC | Nb Positive Infections | % Reads in Positive Animals |
|---|---|---|---|---|---|---|---|---|---|---|---|
| Acidobacteria | Acidobacteria | Acidobacteriales | Acidobacteriaceae | *unclassified Acidobacteriaceae* | 100 | Common soil microbes, known contaminant of molecular biology reagents. | 1 | 53 | 0 | 1 | 38 % |
| | | Pseudonocardiales | Pseudonocardiaceae | Pseudonocardia | 79 | | 1 | 344 | 0 | 1 | 14 % |
| Actinobacteria | Actinobacteria | Bifidobacteriales | Bifidobacteriaceae | Bifidobacterium | 100 | Long thought to be fungi, most are soil or aquatic microbes. Some can cause opportunistic infections in humans, such as *Corynebacterium*, *Dietzia and Williamsia* species. *Mycabacteria* in the sub-order Corynebacteriales are pathogens that cause tuberculosis and leprosy in humans and other mammals, and one species of *Rhodococcus* is an important pathogen of young horses. However, many of these have also been recognized as common contaminants of molecular biology equipment and reagents (indicated here with an asterisk *). | 2 | 649 | 0 | 5 | 28 % |
| | | | Corynebacteriaceae | **Corynebacterium*** | **100** | | **1** | **1853** | **0** | **3** | **32 %** |
| | | | Dietziaceae | **Dietzia*** | **100** | | **1** | **2553** | **0** | **9** | **98 %** |
| | | Corynebacteriales | Nocardiaceae | Rhodococcus* | 100 | | 1 | 240 | 0 | 2 | 47 % |
| | | | | **Williamsia** | **100** | | **1** | **274** | **0** | **2** | **54 %** |
| | | Micrococcales | Brevibacteriaceae | Brevibacterium* | 100 | | 1 | 1251 | 0 | 6 | 33 % |
| | | | Dermabacteraceae | Brachybacterium* | 100 | | 1 | 4787 | 0 | 9 | 94 % |
| | | | Microbacteriaceae | Microbacterium* | 80 | | 1 | 392 | 6.12 % | 1 | 8 % |
| | | | Micrococcaceae | Kocuria | 63 | | 1 | 632 | 0 | 4 | 22 % |
| | | Propionibacteriales | Nocardioidaceae | Marmoricola | 91 | | 1 | 329 | 0 | 1 | 5 % |
| | Coriobacteriia | Coriobacteriales | Coriobacteriaceae | Adlercreutzia | 100 | | 1 | 139 | 0 | 2 | 36 % |
| | | | | Enterorhabdus | 100 | | 1 | 127 | 0 | 1 | 10 % |
| | | | | *unclassified Coriobacteriaceae* | 100 | | 3 | 878 | 0 | 8 | 60 % |
| Bacteroidetes | Bacteroidia | Bacteroidales | Bacteroidaceae | Bacteroides | 100 | Most are normal flora or environmental microbes, rarely associated with opportunistic infections (e.g., *Prevotella* species can cause infection from animal bites). The Flavobacteriaceae family includes a number of important fish pathogens, some respiratory pathogens of birds, and more recently discovered isolates from human sewage and clinical specimens (such as *Elizabethkingia meningoseptica*). One rodent respiratory pathogen has been recently described (*Filobacterium rodentium* gen. nov., sp. nov.) but was not found here. | 5 | 581 | 0 | 7 | 70 % |
| | | | *Candidatus* Homeothermaceae | *unclassified Candidatus* Homeothermaceae | 98 | | 110 | 33495 | 0 | 283 | 68 % |
| | | | Porphyromonadaceae | Odoribacter | 100 | | 1 | 153 | 0 | 1 | 30 % |
| | | | Prevotellaceae | Alloprevotella | 100 | | 5 | 999 | 0 | 5 | 94 % |
| | | | | Prevotella | 100 | | 6 | 2967 | 0 | 11 | 77 % |
| | | | Rikenellaceae | Alistipes | 100 | | 4 | 885 | < 1 % | 6 | 60 % |
| | | | | Rikenella | 100 | | 1 | 65 | 0 | 1 | 65 % |
| | | | unclassified Bacteroidales | *unclassified Bacteroidales* | 94 | | 4 | 997 | 0 | 9 | 73 % |
| | | | | *unclassified Bacteroidales* | 100 | | 1 | 264 | 0 | 1 | 31 % |
| | Flavobacteriia | Flavobacteriales | Flavobacteriaceae | Chryseobacterium | 100 | | 1 | 438 | 0 | 2 | 36 % |
| | | | | Cloacibacterium | 100 | | 1 | 2760 | 0 | 2 | 2 % |
| | | | | Flavobacterium | 100 | | 1 | 673 | 0 | 3 | 46 % |
| | | | | *unclassified Flavobacteriaceae* | 100 | | 3 | 668 | 0 | 4 | 86 % |
| | Sphingobacteriia | Sphingobacteriales | Chitinophagaceae | Sediminibacterium | 100 | | 1 | 10426 | 7.6 % | 1 | 4 % |
| | | | Sphingobacteriaceae | Pedobacter* | 92 | | 2 | 531 | 0 | 2 | 75 % |
| | | | | Sphingobacterium | 100 | | 1 | 80 | 0 | 1 | 20 % |
| | | | | *unclassified Sphingobacteriaceae* | 69 | | 1 | 341 | 0 | 2 | 27 % |
| | unclassified Bacteroidetes | | | *unclassified Bacteroidetes* | 100 | | 3 | 1074 | 0 | 9 | 73 % |
| Cyanobacteria | Melainabacteria | Gastranaerophilales | unclassified Gastranaerophilales | *unclassified Gastranaerophilales* | 100 | Most likely natural gut flora. | 1 | 202 | 0 | 3 | 68 % |
| Firmicutes | Bacilli | Bacillales | Bacillaceae | Geobacillus | 99 | While most Firmicutes species are abundant in normal flora or environmental samples, a number are associated with important diseases and opportunistic infections in animals and humans (naso-pharyngeal diseases from *Streptococcus* spp. infections; among the *Bacillales*: anthrax from infection by *Bacillus anthracis*, and Listeriosis disease of the central nervous system from *Listeria* species; *Clostridium* species infections are probably the most abundant, with clinical relevancy for cases of tetaus, botulism and food poisoning, gangrene, and for diarrhea in patients following antibiotic therapy. | 1 | 306 | 0 | 1 | 13 % |
| | | | Listeriaceae | Brochothrix | 100 | | 1 | 379 | 0 | 1 | 10 % |
| | | | unclassified Bacillales | *unclassified Bacillales* | 90 | | 3 | 1651 | 0 | 4 | 30 % |
| | | Lactobacillales | Carnobacteriaceae | Trichococcus | 59 | | 1 | 53 | 0 | 1 | 43 % |
| | | | Enterococcaceae | Enterococcus | 59 | | 1 | 355 | 0 | 2 | 29 % |
| | | | Lactobacillaceae | Lactobacillus | 96 | | 5 | 7645 | 0 | 30 | 76 % |
| | | | | *unclassified Lactobacillaceae* | 75 | | 1 | 265 | 0 | 2 | 20 % |
| | | | Leuconostocaceae | Leuconostoc | 100 | | 1 | 862 | 0 | 2 | 57 % |
| | | | Streptococcaceae | Lactococcus | 100 | | 1 | 5499 | 0 | 3 | 12 % |
| | | | | **Streptococcus** | **98** | | **5** | **2604** | **0** | **16** | **72 %** |
| | Clostridia | Clostridiales | Anaerococcus | Anaerococcus | 100 | | 1 | 147 | 0 | 1 | 86 % |
| | | | Clostridiaceae | *Candidatus* Arthromitus | 100 | | 1 | 98 | 0 | 1 | 93 % |
| | | | | Clostridium *sensu stricto* 12 | 97 | | 1 | 59 | 0 | 1 | 56 % |
| | | | Defluviitaleaceae | *Incertae sedis* | 100 | | 1 | 68 | 0 | 2 | 62 % |
| | | | Defluviitaleaceae | *unclassified Defluviitaleaceae* | 100 | | 1 | 189 | 0 | 1 | 38 % |
| | | | Lachnospiraceae | Acetatifactor | 82 | | 5 | 1676 | 0 | 19 | 64 % |
| | | | | Anaerostipes | 100 | | 1 | 87 | 0 | 1 | 23 % |
| | | | | Blautia | 51 | | 19 | 10445 | 0 | 81 | 64 % |
| | | | | Dorea | 69 | | 1 | 64 | 0 | 1 | 28 % |
| | | | | *Incertae sedis* | 70 | | 1 | 68 | 0 | 1 | 28 % |
| | | | | Roseburia | 66 | | 6 | 1381 | 0 | 12 | 62 % |

| Phylum | Class | (sub-)Order | Family | Genus | Minimum Bootstrap | Biology | Nb of OTUs | Total Nb Reads (incl. NCs) | % reads in NC | Nb Positive Infections | % Reads in Positive Animals |
|---|---|---|---|---|---|---|---|---|---|---|---|
| Firmicutes (cont.) | | | | *unclassified Lachnospiraceae* | 52 | | 72 | 23886 | 0 | 189 | 68 % |
| | | | Peptostreptococcaceae | *Incertae sedis* | 100 | | 1 | 161 | 0 | 1 | 52 % |
| | | | Ruminococcaceae | Anaerotruncus | 77 | | 1 | 98 | 0 | 2 | 40 % |
| | | | | Faecalibacterium | 100 | | 1 | 356 | 0 | 2 | 93 % |
| | | | | *Incertae sedis* | 57 | | 5 | 1892 | 0 | 21 | 61 % |
| | | | | Intestinimonas | 79 | | 3 | 618 | 0 | 6 | 25 % |
| | | | | Oscillibacter | 83 | | 4 | 2081 | < 0.1 % | 19 | 55 % |
| | | | | Ruminococcus | 89 | | 11 | 3779 | < 1 % | 34 | 70 % |
| | | | | *unclassified Ruminococcaceae* | 59 | | 24 | 16720 | 0 | 116 | 74 % |
| | | | vadinBB60 | *unclassified Clostridiales vadinBB60 genus* | 72 | | 13 | 2560 | 0 | 32 | 57 % |
| | | | unclassified Clostridiales | *unclassified Clostridiales* | 100 | | 2 | 9590 | 0 | 32 | 82 % |
| | | | | *unclassified Clostridiales* | 100 | | 2 | 956 | 0 | 8 | 72 % |
| | | unclassified Clostridia | | *unclassified Clostridia* | 100 | | 1 | 516 | 0 | 2 | 27 % |
| | Erysipelotrichia | Erysipelotrichales | Erysipelotrichaceae | Allobaculum | 99 | | 2 | 911 | 0 | 6 | 90 % |
| | | Erysipelotrichales | | Catenibacterium | 100 | | 1 | 256 | 0 | 1 | 96 % |
| | Negativicutes | Selenomonadales | Acidaminococcaceae | Phascolarctobacterium | 100 | | 1 | 157 | 0 | 2 | 83 % |
| | | Selenomonadales | Veillonellaceae | Veillonella | 100 | | 1 | 268 | 0 | 1 | 24 % |
| | unclassified Firmicutes | | | *unclassified Firmicutes* | 100 | | 1 | 53 | 0 | 1 | 49 % |
| Fusobacteria | Fusobacteriia | Fusobacteriales | Fusobacteriaceae | **Fusobacterium** | **100** | Fusobacterium species are associated with a range of gastro-intestinal diseases and skin ulcerations, but it is not clear to what degree this association is opportunistic. | **1** | **108** | **0** | **2** | **94 %** |
| Proteobacteria | Alphaproteobacteria | Caulobacterales | Caulobacteraceae | Brevundimonas | 100 | Many *Proteobacteria* are important animal pathogens, including those with sustained zoonotic transmission to humans. | 1 | 275 | 0 | 1 | 28 % |
| | | Rhizobiales | Bartonellaceae | **Bartonella** | **100** | *Bartonella* species are thought to have undergone a recent expansion and now commonly infect rodent species around the world. Many of these species are implicated in causing zoonotic human infections. | **1** | **6327077** | **0.01** | **245** | **99%** |
| | | | Methylobacteriaceae | Methylobacterium* | 100 | The remaining *Rhizobiales* and *Rhodobacterales* are important soil microbes. | 2 | 7113 | 1.8 % | 4 | 7 % |
| | | | Phyllobacteriaceae | Phyllobacterium | 70 | | 1 | 120 | 0 | 1 | 39 % |
| | | | Rhizobiaceae | Rhizobium | 100 | | 1 | 968 | 1.96 % | 1 | 40 % |
| | | Rhodobacterales | Rhodobacteraceae | Paracoccus | 89 | | 1 | 1229 | 1.46 % | 1 | 3 % |
| | | Rickettsiales | Anaplasmataceae | ***Candidatus* Neoehrlichia** | **100** | Causes anaplasmosis in humans and domestic animals, carried by a diverse array of rodents as well as *Ixodes* tick vectors. 100% sequence identity to *Candidatus N. mukarensis*. | **1** | **18358** | **0** | **8** | **99 %** |
| | | | Rickettsiaceae | **Orientia** | **100** | The only species in this order, *Orientia tsutsugamushi*, causes scrub typhus in humans and is transmitted by chiggers. | **1** | **876** | **0** | **8** | **91 %** |
| | | | | **Rickettsia** | **100** | Closest sequence identities (98%) were to the spotted fever group of *Rickettsia* species, responsible for a number of important vector-transmitted human and animal diseases. | **1** | **57226** | **0** | **8** | **99 %** |
| | | Sphingomonadales | Sphingomonadaceae | Novosphingobium | 95 | The Sphingomonads have been implicated in nosocomial infections, but are widly distributed in nature with a diverse array of biological characteristics. However, in our dataset, positive results apppeared more likely to be contaminants. | 1 | 235 | 0 | 1 | 8 % |
| | | | | Sphingobium | 100 | | 1 | 574 | 0 | 2 | 34 % |
| | | | | Sphingomonas | 100 | | 1 | 593 | 0 | 1 | 31 % |
| | Betaproteobacteria | Burkholderiales | Burkholderiaceae | Lautropia | 100 | Though some *Burkholderiaceae* are well-known pathogens (e.g., whooping cough agent *Bordatella pertussis*), this is not expected for any of the *Burkholderales* found in our study. | 1 | 58 | 0 | 1 | 71 % |
| | | | | Ralstonia* | 97 | | 1 | 1624 | 0 | 2 | 4 % |
| | | | Comamonadaceae | *unclassified Comamonadaceae* | 100 | | 1 | 3407 | < 1 % | 7 | 17 % |
| | | | Oxalobacteraceae | Duganella | 65 | | 1 | 266 | 0 | 2 | 39 % |
| | | Hydrogenophilales | Hydrogenophilaceae | Hydrogenophilus | 100 | Among the remaining Betaproteobacteria, the only recognized pathogenic species are a few of many in the genus *Neisseria*. One infection carrying the bulk of reads shared 98% sequence identity to bacteria found in healthy human nasal passages. | 1 | 2617 | 2.98 % | 1 | 6 % |
| | | Neisseriales | Neisseriaceae | ***unclassified Neisseriaceae*** | **97** | | **2** | **928** | **0** | **2** | **95 %** |
| | | Rhodocyclales | Rhodocyclaceae | Dechloromonas | 94 | | 1 | 302 | 0 | 1 | 95 % |
| | | unclassified Betaproteobacteria | | *unclassified Betaproteobacteria* | 100 | | 1 | 143 | 1.4 % | 1 | 8 % |

| Phylum | Class | (sub-)Order | Family | Genus | Minimum Bootstrap | Biology | Nb of OTUs | Total Nb Reads (incl. NCs) | % reads in NC | Nb Positive Infections | % Reads in Positive Animals |
|---|---|---|---|---|---|---|---|---|---|---|---|
| Proteobacteria (continued) | Deltaproteobacteria | Desulfovibrionales | Desulfovibrionaceae | Desulfovibrio | 74 | *Deltaproteobacteria* live in extreme environments, and only one pathogenic species is currently known (*Lawsonia intercellularis* in horses). | 8 | 1690 | 0 | 22 | 78 % |
| | | | | *unclassified Desulfovibrionaceae* | 61 | | 5 | 962 | 0 | 10 | 72 % |
| | | unclassified Deltaproteobacteria | | *unclassified Deltaproteobacteria* | 100 | | 1 | 52 | 0 | 1 | 100 % |
| | Epsilonproteobacteria | Campylobacterales | Campylobacteraceae | **Arcobacter** | **100** | Very little is known about the Arcobacter species, but their potential for pathogenicity in animals has been recently described (Houf & Stephan 2007). We noted 100% sequence homology to *Arcobacter cryaerophilus*, which has been associated with disease in humans and laboratory rats. | **1** | **403** | **0** | **2** | **85 %** |
| | | | Helicobacteraceae | **Helicobacter** | **100** | While *Helicobacter* are common among gut flora, dysbiosis resulting in systemic infection is associated with pathogenesis. The two main OTUs shared closest (96-97%) sequence identity to *H. suncus* (isolated from shrews with chronic gastritis, Helico1) and *H. trogontum* (another enterohepatic *Helicobacter* spp. associated with intestinal disease, Helico2) | **5** | **47394** | **0** | **56** | **93 %** |
| | Gammaproteobacteria | Alteromonadales | Pseudoalteromonadaceae | Pseudoalteromonas | 100 | Marine bacteria. | 1 | 116 | 0 | 1 | 32 % |
| | | | Shewanellaceae | Shewanella | 100 | | 1 | 78 | 0 | 1 | 23 % |
| | | Enterobacteriales | Enterobacteriaceae | Escherichia-Shigella | 91 | | 1 | 979 | 3.17 % | 2 | 29 % |
| | | | | Providencia | 96 | | 1 | 600 | 0 | 1 | 88 % |
| | | | | **Yersinia** | **85** | Shigellosis, plague, and typhoid fever are just a few pathogenic *Enterobacteriaceae*, many known to also be carried by rodents. However, our methods did not allow for sufficient distinction between pathogenic and non-pathogenic occurrence or contamination from gut flora. | **1** | **7414** | **0** | **14** | **85 %** |
| | | | | *unclassified Enterobacteriaceae* | 93 | | 1 | 964 | 0 | 3 | 50 % |
| | | Legionellales | Coxiellaceae | **Rickettsiella** | **76** | Abundant endosymbiont in *Ixodes* ticks, thought to have lead to emergence of Q-fever (*Coxiella burnetii*); 100% sequence identity to *Rickettsiella grylli* and other entomopathic isolates. One animal with a relatively high number of reads (128,46, BR10117) | **1** | **592** | **0** | **3** | **45 %** |
| | | Oceanospirillales | Oceanospirillaceae | Marinospirillum | 100 | Marine bacterium. | 1 | 263 | 0 | 1 | 97 % |
| | | Pasteurellales | Pasteurellaceae | ***unclassified Pasteurellaceae*** | **81** | 99% sequence similarity to the newly reclassified Muribacter muris, described among samples taken from mice and rats, whose closest ancestor is the epizootic Haemophilus influenzae-murium. While some species of *Pasteurella* are zoonotic and pathogenic to humans, this is not the case for other mammalian hosts. | **2** | **1951** | **0** | **5** | **54 %** |
| | | Pseudomonadales | Moraxellaceae | Acinetobacter | 97 | Among the *Pseudomonadales*, the *Pseudomonas* (nosocomial superbug) and *Psychrobacter* (endocarditis-causing) species have the largest implications for pathogenicity. However, along with *Acinetobacter*, disease is mostly limited to opportunistic infection. | 4 | 11290 | 2.3 % | 14 | 32 % |
| | | | | Alkanindiges | 100 | | 1 | 184 | 0 | 1 | 9 % |
| | | | | Enhydrobacter | 100 | | 1 | 8497 | 1.06 % | 1 | 7 % |
| | | | | Psychrobacter | 100 | | 3 | 1033 | < 0.1 % | 5 | 62 % |
| | | | Pseudomonadaceae | Pseudomonas | 100 | | 2 | 18473 | 1.72 % | 20 | 56 % |
| | | | | *unclassified Pseudomonadaceae* | 100 | | 2 | 179 | 0 | 2 | 64 % |
| | | Vibrionales | Vibrionaceae | Vibrio | 96 | *Vibrio* species are important human pathogens (e.g., causing cholera, food poisoning), common aquatic bacteria. | 1 | 182 | 0 | 2 | 64 % |
| | | Xanthomonadales | Xanthomonadaceae | Stenotrophomonas | 100 | Human opportunists, particularly in nosocomial settings. | 1 | 1123 | 0 | 2 | 14 % |
| | | unclassified Gammaproteobacteria | | *unclassified Gammaproteobacteria* | 100 | Unknown | 1 | 63 | 0 | 1 | 76 % |

| Phylum | Class | (sub-)Order | Family | Genus | Minimum Bootstrap | Biology | Nb of OTUs | Total Nb Reads (incl. NCs) | % reads in NC | Nb Positive Infections | % Reads in Positive Animals |
|---|---|---|---|---|---|---|---|---|---|---|---|
| Proteobacteria (continued) | unclassified Proteobacteria | | | *unclassified Proteobacteria* | 100 | Unknown | 4 | 970 | 0 | 7 | 68 % |
| Spirochaetae | Spirochaetes | Spirochaetales | Brevinemataceae | **Brevinema** | **100** | 96% sequence identity to *Brevinema andersonii*, an infectious spirochaete of the short-tailed shrew and white footed mouse in North America. | **1** | **5603** | **0** | **3** | **100 %** |
| | | | Leptospiraceae | **Leptospira** | **100** | 100% sequence identity to human pathogenic *Leptospira borgpetersenii* and *L. interrogans*; rodents are considered the primary reservoir species | **1** | **257** | **0** | **2** | **95 %** |
| | | | Spirochaetaceae | **Borrelia** | **100** | Borrelia1: 100% sequence identity to *Borrelia miyamotoi* in humans and *Ixodes* ticks, zoonotic fever that may relapse (co-infects with but is not Lyme disease agent *B. burgdorferi*); Borrelia2: 96% sequence identity to *Borrelia* sp. *Nov* in *Peromyscus leucopus;* Borrelia3: positive in two animals beyond positive control reads | **3** | **703** | **< 1 %** | **5** | **90 %** |
| | | | | Treponema | 99 | *Treponema* species are pathogenic in humans (e.g., syphilis), but many occur in natural gut flora such as the rumen of cows. The closest sequence identity (92%) of our samples is to these non-pathogenic species, and they were found only in animals with several gut microbiota (probable contaminated dissection). | 2 | 985 | 0 | 7 | 77 % |
| Tenericutes | Mollicutes | Anaeroplasmatales | Anaeroplasmataceae | Anaeroplasma | 100 | Rumen microbes. | 1 | 137 | 0 | 1 | 22 % |
| | | Entomoplasmatales | Spiroplasmataceae | **Spiroplasma** | **100** | 95% species identity with pathogenic Spiroplasma species isolated from Ixodes ticks, 94% identity with samples pulled from ticks on dogs (Spiroplasma eriocheiris); The pathogenicity of these species is disputed, with questions revolving around its role in transmissible spongiform encephylopathies in humans and animals. 1 animal very infected (BR10049 >2000 reads per sample), second animal low copy numbers (BR10114 <15 copies per sample) | **1** | **4738** | **0** | **2** | **100 %** |
| | | Mycoplasmatales | Mycoplasmataceae | **Mycoplasma** | **100** | *Mycoplasmae* species are pathogenic in both humans and animals, implicated in respiratory and pelvic infections. Some OTUs were overlapping, and more detail is provided in the text. | **11** | **1786662** | **2.56 %** | **204** | **99%** |
| | | | | ***unclassified Mycoplasma species*** | **100** | | **1** | **39767** | **0** | **10** | **100%** |
| Verrucomicrobia | Spartobacteria | Chthoniobacterales | unclassified Chthoniobacterales | *unclassified Chthoniobacterales* | 100 | Not pathogenic | 1 | 786 | 0 | 2 | 11 % |
| | Verrucomicrobiae | Verrucomicrobiales | Verrucomicrobiaceae | Luteolibacter | 98 | | 1 | 147 | 0 | 1 | 24 % |
| *entirely unclassified Bacteria* | | | | | 100 | Unknown | 25 | 4390 | < 1 % | 37 | 65 % |
| *Eukaryote* | | | | | | | | | | | |
| Apicomplexa | Conoidasida | Eucoccidiorida | Sarcocystidae | ***unclassified Sarcocystidae*** | **100** | Protazoan intracellular organisms in the family Sarcocystidae cause a variety of diseases in animals and humans, such as Toxoplasmosis *(Toxoplasma gondii* 90-96% similarity) and reproductive failure (*Neospora caninum* 92-97% similarity). 98% sequence similarity to Sacrocystis muris, a coccidian parasite closely related to Toxoplasmae and Neurosporae, first found in mice. | **3** | **13581** | **0** | **15** | **95 %** |

* Indicates taxa found to be frequent contaminants of molecular biology reagents and consumables
