## Supplementary material for "Pathogen community composition and co-infection patterns in a wild community of rodents": Table S2

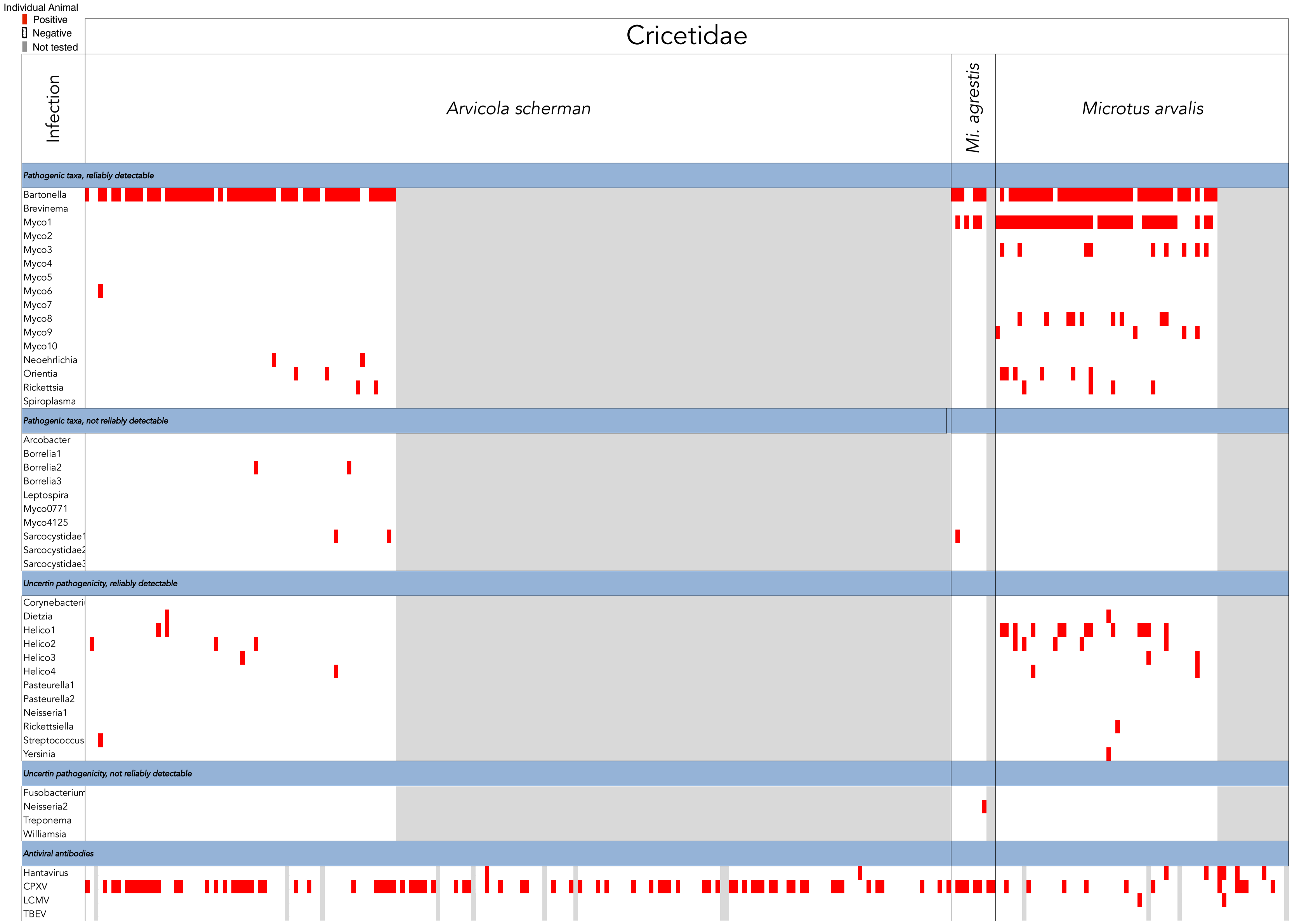
Individual Animal
Positive
Negative
Not tested
Cricetidae
Infection
Arvicola scherman
Mi. agrestis
Microtus arvalis
Pathogenic taxa, reliably detectable
Bartonella
Brevinema
Myco1
Myco2
Myco3
Myco4
Myco5
Myco6
Myco7
Myco8
Myco9
Myco10
Neoehrlichia
Orientia
Rickettsia
Spiroplasma
Pathogenic taxa, not reliably detectable
Arcobacter
Borrelia1
Borrelia2
Borrelia3
Leptospira
Myco0771
Myco4125
Sarcocystidae1
Uncertin pathogenicity, reliably detectable
Dietzia
Helico1
Helico2
Helico3
Helico4
Pasteurella1
Pasteurella2
Neisseria1
Rickettsiella
Streptococcus
Yersinia
Uncertin pathogenicity, not reliably detectable
Neisseria2
Treponema
Williamsia
Antiviral antibodies
Hantavirus
CPXV
LCMV
TBEV

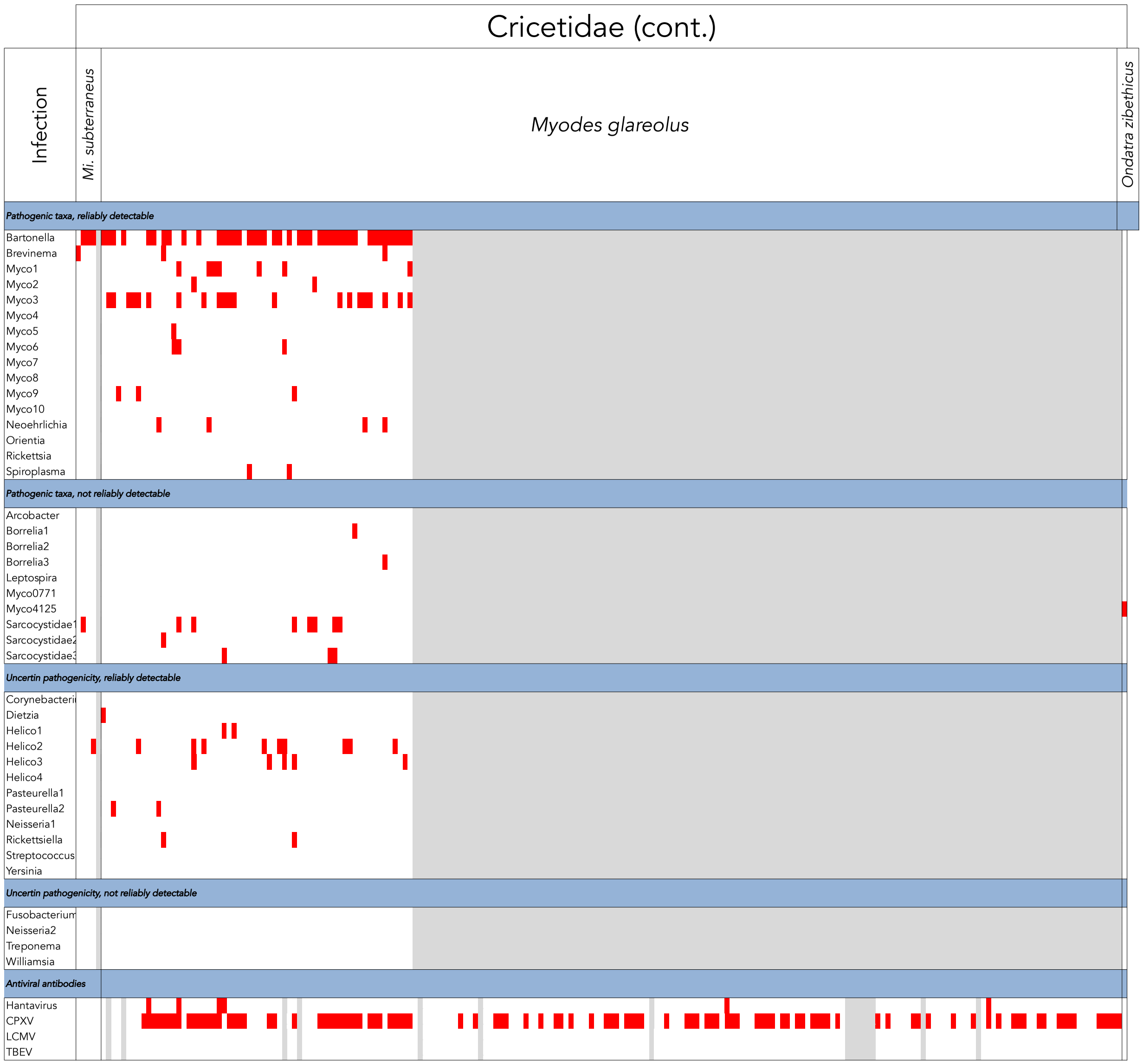
Cricetidae (cont.)
Infection
Mi. subterraneus
Myodes glareolus
Ondatra zibethicus
Pathogenic taxa, reliably detectable
Bartonella
Brevinema
Myco1
Myco2
Myco3
Myco4
Myco5
Myco6
Myco7
Myco8
Myco9
Myco10
Neoehrlichia
Orientia
Rickettsia
Spiroplasma
Pathogenic taxa, not reliably detectable
Arcobacter
Borrelia1
Borrelia2
Borrelia3
Leptospira
Myco0771
Myco4125
Sarcocystidae1
Sarcocystidae2
Sarcocystidae3
Uncertin pathogenicity, reliably detectable
Corynebacteri
Dietzia
Helico1
Helico2
Helico3
Helico4
Pasteurella1
Pasteurella2
Neisseria1
Rickettsiella
Streptococcus
Yersinia
Uncertin pathogenicity, not reliably detectable
Fusobacterium
Neisseria2
Treponema
Williamsia
Antiviral antibodies
Hantavirus
CPXV
LCMV
TBEV

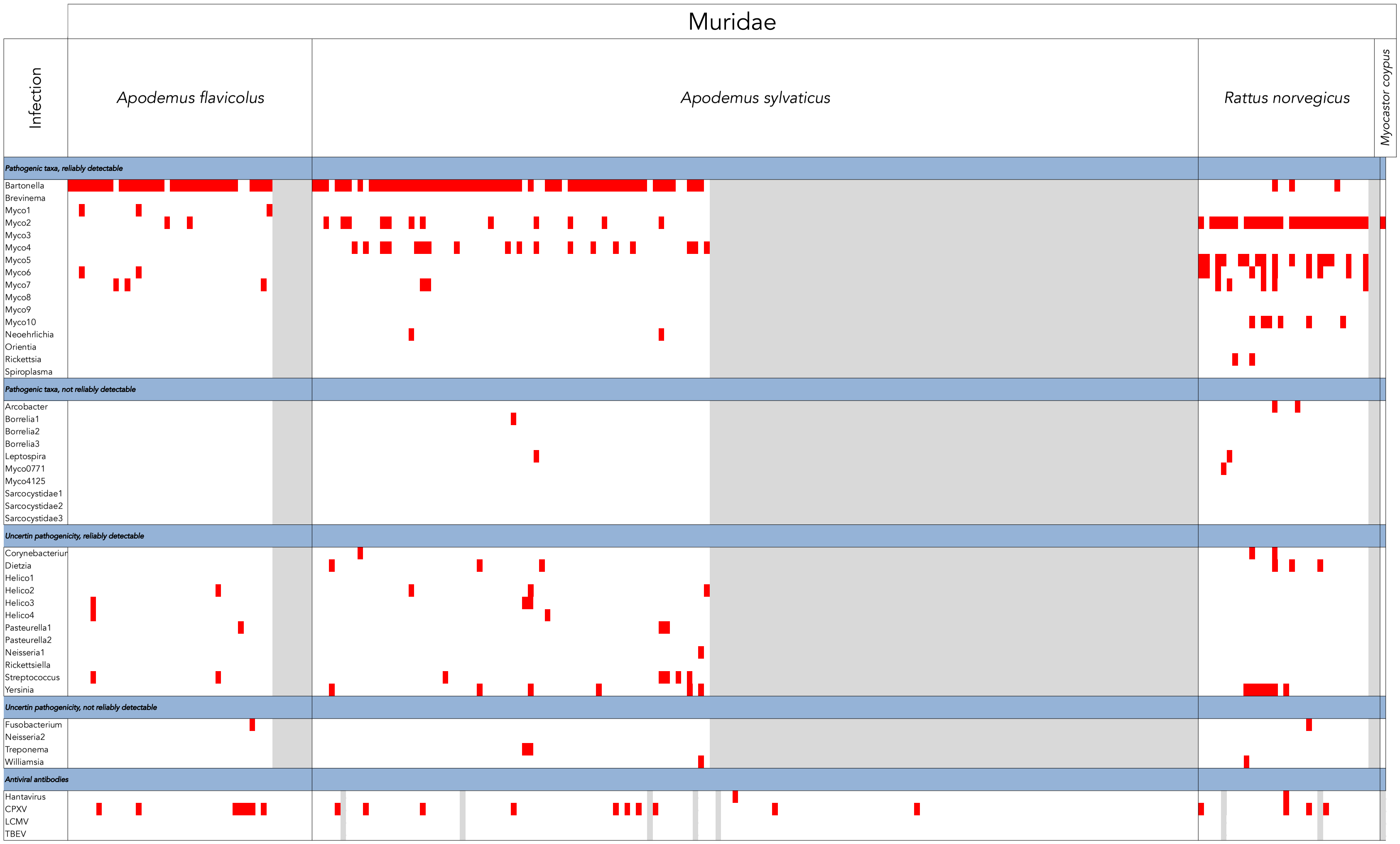
Muridae
Infection
Apodemus flavicolus
Apodemus sylvaticus
Rattus norvegicus
Myocastor coypus
Pathogenic taxa, reliably detectable
Bartonella
Brevinema
Myco1
Myco2
Myco3
Myco4
Myco5
Myco6
Myco7
Myco8
Myco9
Myco10
Neoehrlichia
Orientia
Rickettsia
Spiroplasma
Pathogenic taxa, not reliably detectable
Arcobacter
Borrelia1
Borrelia2
Borrelia3
Leptospira
Myco0771
Myco4125
Sarcocystidae1
Sarcocystidae2
Sarcocystidae3
Uncertin pathogenicity, reliably detectable
Corynebacteriur
Dietzia
Helico1
Helico2
Helico3
Helico4
Pasteurella1
Pasteurella2
Neisseria1
Rickettsiella
Streptococcus
Yersinia
Uncertin pathogenicity, not reliably detectable
Fusobacterium
Neisseria2
Treponema
Williamsia
Antiviral antibodies
Hantavirus
CPXV
LCMV
TBEV

### Summaries

| Infection | Total Nb Positive Animals | NbReads PC | NbReads NC | Total NbReads minusPC | NbReads Positive Animals | % Reads in Positive Animals | Bacterial OTU |
|---|---|---|---|---|---|---|---|
| *Pathogenic taxa, reliably detectable* | | | | | | | |
| Bartonella | 245 | 26295 | 575 | 6327077 | 6245151 | 99% | Otu00001 |
| Brevinema | 3 | 0 | 0 | 5603 | 5597 | 100% | Otu00123 |
| Myco1 | 55 | 0 | 453 | 845971 | 833501 | 99% | Otu00004 |
| Myco2 | 44 | 0 | 2 | 426034 | 423543 | 99% | Otu00003 |
| Myco3 | 30 | 0 | 0 | 106443 | 102028 | 96% | Otu00006 |
| Myco4 | 18 | 0 | 0 | 72724 | 72414 | 100% | Otu00010 |
| Myco5 | 17 | 0 | 1 | 165095 | 164713 | 100% | Otu00005 |
| Myco6 | 16 | 0 | 1 | 92237 | 92010 | 100% | Otu00007 |
| Myco7 | 10 | 0 | 0 | 39767 | 39580 | 100% | Otu00012 |
| Myco8 | 9 | 0 | 0 | 31528 | 31448 | 100% | Otu00015 |
| Myco9 | 7 | 0 | 0 | 40125 | 39661 | 99% | Otu00049 |
| Myco10 | 6 | 0 | 0 | 6086 | 6043 | 99% | Otu00054 |
| Neoehrlichia | 8 | 0 | 0 | 18358 | 18266 | 99% | Otu00039 |
| Orientia | 8 | 0 | 0 | 876 | 799 | 91% | Otu00111 |
| Rickettsia | 8 | 14872 | 0 | 57226 | 56859 | 99% | Otu00008 |
| Spiroplasma | 2 | 0 | 0 | 4738 | 4723 | 100% | Otu00093 |
| *Pathogenic taxa, not reliably detectable* | | | | | | | |
| Arcobacter | 1 | 0 | 0 | 403 | 342 | 85% | Otu00296 |
| Borrelia1 | 2 | 0 | 0 | 419 | 419 | 100% | Otu00318 |
| Borrelia2 | 2 | 0 | 0 | 206 | 158 | 77% | Otu00514 |
| Borrelia3 | 1 | 9706 | 3 | 78 | 53 | 68% | Otu00071 |
| Leptospira | 2 | 0 | 0 | 257 | 243 | 95% | Otu01015 |
| Myco0771 | 1 | 0 | 0 | 255 | 255 | 100% | Otu00771 |
| Myco4125 | 1 | 0 | 0 | 164 | 164 | 100% | Otu04125 |
| Sarcocystidae1 | 11 | 0 | 0 | 8684 | 8010 | 92% | Otu00056 |
| Sarcocystidae2 | 1 | 0 | 0 | 3678 | 3657 | 99% | Otu00191 |
| Sarcocystidae3 | 3 | 0 | 0 | 1219 | 1196 | 98% | Otu00254 |
| *Uncertin pathogenicity, reliably detectable* | | | | | | | |
| Corynebacterium | 3 | 0 | 0 | 1853 | 600 | 32% | Otu00050 |
| Dietzia | 9 | 73 | 0 | 2553 | 2513 | 98% | Otu00102 |
| Helico1 | 17 | 0 | 0 | 34894 | 34678 | 99% | Otu00013 |
| Helico2 | 22 | 0 | 0 | 8702 | 6158 | 71% | Otu00025 |
| Helico3 | 11 | 0 | 0 | 2303 | 2153 | 93% | Otu00087 |
| Helico4 | 5 | 0 | 0 | 1178 | 874 | 74% | Otu00128 |
| Pasteurella1 | 3 | 0 | 0 | 1430 | 801 | 56% | Otu00129 |
| Pasteurella2 | 2 | 0 | 0 | 521 | 246 | 47% | Otu00203 |
| Neisseria1 | 1 | 0 | 0 | 780 | 745 | 96% | Otu00612 |
| Rickettsiella | 3 | 0 | 0 | 592 | 264 | 45% | Otu00187 |
| Streptococcus | 8 | 0 | 0 | 1681 | 1324 | 79% | Otu00115 |
| Yersinia | 14 | 6 | 0 | 7414 | 6295 | 85% | Otu00041 |
| *Uncertin pathogenicity, not reliably detectable* | | | | | | | |
| Fusobacterium | 2 | 0 | 0 | 108 | 101 | 94% | Otu00791 |
| Neisseria2 | 1 | 0 | 0 | 148 | 132 | 89% | Otu00454 |
| Treponema | 2 | 0 | 0 | 348 | 127 | 36% | Otu00235 |
| Williamsia | 2 | 0 | 0 | 274 | 148 | 54% | Otu00614 |
| *Antiviral antibodies* | | | | | | | |
| Hantavirus | 16 | | | | | | |
| CPXV | 222 | | | | | | |
| LCMV | 2 | | | | | | |
| TBEV | 0 | | | | | | |
