## Supplemental Materials for "Pathogen community composition and co-infection patterns in a wild community of rodents"

Updated: 19 June 2023

#### Supplemental Materials

**TABLES 3**

**Table S1: Catalogue of 16S OTUs detected. (separate PDF file) 3**

**Table S2: Occurrence data for each potentially pathogenic OTU and antiviral antibody in each individual rodent. (separate PDF file) 4**

**Table S3: Extrinsic drivers of pathogen diversity in a rodent community. 5**

**Table S4. The role of Rattus norvegicus hosts as extrinsic drivers of pathogen diversity in a rodent community. 6**

**Table S5. Results of permutational multivariate analysis of variance on extrinsic contributors to pathogen community composition. 7**

**Table S6. Extrinsic drivers of pathogen community composition in a rodent community. 8**

**Table S7: Association analyses between Myco1, Myco3, and Hantavirus exposure in *Mi. arvalis* and *My. glareolus*. 9**

**Table S8. Association between Myco2 and Myco4 infections in Ap. sylvaticus. 10**

**Table S9: Association between Bartonella spp. infection and cowpox virus (CPXV) exposure across rodent species. 11**

**Table S10: Association between *Mycoplasma haemomuris* (MH) and *Myco. coccoides* (MC) infections in the four rodent species where they both circulate. 13**

**Table S11: Association between *Bartonella* spp. and hemotropic *Mycoplasma* spp. infections across rodent species. 14**

**FIGURES 17**

**Figure S1. Landscape of trapping locations for focal host species. 17**

**Figure S2. Host species demographic distributions (sex and age class). 18**

**Figure S3. Data filtering process for 16S meta-barcoding identification of pathogenic bacteria. 19**

**Figure S4. Phylogeny of Mycoplasma species and OTUs. 20**

**Figure S5. Results of Akaike Information Criterion (AIC)-based model selection analysis of the effect of extrinsic factors on pathogen diversity. 21**

**Figure S6. Cook's distances for co-infection outliers. 22**

**Figure S7. Results of multiple correspondence analysis (MCA) for pathogen community composition in rodents (including *R. norvegicus*)** 23

**Figure S8. Variance in pathogen community structure among (A) host species and (B) habitats including *R. norvegicus* from farms.** 24

**Figure S9. Data used for pathogen community composition multiple correspondence analysis (MCA).** 25

**Figure S10: Pathogen community structure among extrinsic factors** 26

**Figure S11. Association Screening (SCN) analysis results for Myco1, Myco3, and Hantavirus antibodies in *Mi. arvalis* and *My. glareolus* host species.** 27

**Figure S12. Results of Akaike Information Criterion (AIC)-based model selection analysis of associations between occurrence of Myco1, Myco3 and hantavirus exposure.** 28

**Figure S13. Results of Akaike Information Criterion (AIC)-based model selection analysis of associations between occurrence of Myco2 and Myco4.** 29

**Figure S14. Results of Akaike Information Criterion (AIC)-based model selection analysis of associations between occurrence of Bartonella and CPXV exposure.** 30

**Figure 15: Results of Akaike Information Criterion (AIC)-based model selection analysis of associations between occurrence of *Mycoplasma haemomuris* (MH) and *Mycoplasma coccoides* (MC).** 33

**Figure 16: Results of Akaike Information Criterion (AIC)-based model selection analysis of associations between occurrence of *Bartonella* spp. and hemotropic *Mycoplasma* spp. infections across rodent species.** 34

**Figure S17. Benjamini-Hochberg correction for multiple tests.** 36

**APPENDICES** **37**

**Appendix 1: Description of rare host species, their pathogens, and other under-detected pathogens.** 37

**Appendix 2: Scripts and data for statistical analyses** 39

**Appendix 3: References for Table 1** 40

### Tables

#### Table S1: Catalogue of 16S OTUs detected. (separate PDF file)

See PDF: Table_S1.pdf

#### Table S2: Occurrence data for each potentially pathogenic OTU and antiviral antibody in each individual rodent. (separate PDF file)

See PDF: Table_S2.pdf

Table S3: Extrinsic drivers of pathogen diversity in a rodent community. Significant differences in Shannon diversity index were tested on residual variance of the multiple regression model controlling for all other extrinsic factors. Bold factors and different group letters (Tukey-adjusted for multi-level factors) signify statistically significant differences at $p < 0.05$.

| *Residual df = 78* | | Pathogen Diversity (Shannon Index) | | | | | |
|---|---|---|---|---|---|---|---|
| Extrinsic Factor | | residual mean [sd] | n | group | df | *F* | *p*-value |
| Host species | | | | | **5** | **6.86** | **< 0.0001** |
| | *Ar. scherman* | **-0.43 [0.35]** | **9** | **C** | | | |
| | *Mi. agrestis* | **-0.24 [0.50]** | **6** | **BC** | | | |
| | *Mi. arvalis* | **0.27 [0.03]** | **14** | **AB** | | | |
| | *My. glareolus* | **0.36 [0.39]** | **16** | **A** | | | |
| | *Ap. flavicollis* | **-0.16 [0.48]** | **16** | **BC** | | | |
| | *Ap. sylvaticus* | **-0.08 [0.45]** | **20** | **BC** | | | |
| Habitat type | | | | | **2** | **4.97** | **0.0096** |
| | Forest | **-0.12 [0.46]** | **27** | **B** | | | |
| | Hedgerows | **0.06 [0.43]** | **34** | **AB** | | | |
| | Meadows | **0.07 [0.31]** | **20** | **A** | | | |
| Study Site | | | | | 1 | 0.42 | 0.51 |
| | Boult-aux-Bois | *ns* | | *ns* | | | |
| | Briquenay | *ns* | | *ns* | | | |
| Year | | | | | 1 | 1.38 | 0.24 |
| | 2010 | *ns* | | *ns* | | | |
| | 2011 | *ns* | | *ns* | | | |
| Age group | | | | | **1** | **21.43** | **< 0.0001** |
| | Adult | **0.12 [0.40]** | **60** | **A** | | | |
| | Juvenile | **-0.35 [0.42]** | **21** | **B** | | | |
| Sex | | | | | 1 | 1.23 | 0.27 |
| | Male | *ns* | | *ns* | | | |
| | Female | *ns* | | *ns* | | | |

Table S4. The role of Rattus norvegicus hosts as extrinsic drivers of pathogen diversity in a rodent community. Significant differences in Shannon diversity index were tested on residual variance of the multiple regression model controlling for study site, study year, host age class, and host sex. Different group letters signify statistically significant differences at $p < 0.05$ with post-hoc Tukey's adjustment for multiple tests.

Pathogen Diversity (Shannon Index)

| Host Species | residual mean [sd] | n | group |
|---|---|---|---|
| *Ar. scherman* | **-0.27 [0.35]** | **9** | **BC** |
| *Mi. agrestis* | **-0.31 [0.54]** | **6** | **C** |
| *Mi. arvalis* | **0.41 [0.36]** | **14** | **A** |
| *My. glareolus* | **0.21 [0.42]** | **16** | **AB** |
| *Ap. flavicollis* | **-0.28 [0.49]** | **16** | **C** |
| *Ap. sylvaticus* | **-0.18 [0.48]** | **20** | **BC** |
| *R. norvegicus* | **0.32 [0.36]** | **10** | **AB** |

Table S5. Results of permutational multivariate analysis of variance on extrinsic contributors to pathogen community composition. Results of permutational multivariate analysis of variance (PERMANOVA) for the effect of extrinsic factors on pathogen community composition defined via a Bray-Curtis dissimilarity matrix.

| | **Pathogen community composition (Bray-Curtis dissimilarity matrix)** | | | |
|---|---|---|---|---|
| *Residual df = 252* | df | *Sum of Squares* | *F* | *p-value* |
| Extrinsic Factor | | | | |
| **Host species** | **5** | **10.55** | **16.23** | **0.001** |
| **Habitat type** | **2** | **0.65** | **2.51** | **0.024** |
| Study site | 1 | 0.18 | 1.41 | 0.23 |
| Year | 1 | 0.24 | 1.81 | 0.15 |
| Age group | 1 | 0.23 | 1.78 | 0.14 |
| Sex | 1 | 0.075 | 0.58 | 0.67 |

Table S6. Extrinsic drivers of pathogen community composition in a rodent community. Orthogonal response variables MCA Dim1 - MCA Dim7, summarizing the majority of variation in pathogen occurrence, were tested individually. Each linear multiple regression model included all six extrinsic factors and analysis of deviance was performed to evaluate significance of each factor after first controlling for all other factors in the model. Tukey's HSD (at $p < 0.05$) was performed to determine the significance of post-hoc tests between factor levels. Factor levels with different letters were significantly different from one-another. Bold factors indicate factors with significant differences between levels following post-hoc tests. Figures in grey showed non-significant trends ($p \leq 0.1$).

*Residual df = 268*

| Exctrinsic Factor | MCA Dim1 residual mean [sd] | group | df | *F* | p-value | MCA Dim2 residual mean [sd] | group | df | *F* | p-value | MCA Dim3 residual mean [sd] | group | df | *F* | p-value |
|---|---|---|---|---|---|---|---|---|---|---|---|---|---|---|---|
| Host species | | | **5** | **23.83** | **< 0.0001** | | | **5** | **3.89** | **0.002** | | | **5** | **10.60** | **< 0.0001** |
| *Ar. scherman* | **-0.14 [0.13]** | **B** | | | | **0.01 [0.20]** | **AB** | | | | **-0.09 [0.18]** | **C** | | | |
| *Mi. agrestis* | **0.11 [0.22]** | **AB** | | | | **-0.17 [0.19]** | **B** | | | | **-0.15 [0.12]** | **C** | | | |
| *Mi. arvalis* | **0.23 [0.38]** | **A** | | | | **-0.02 [0.36]** | **AB** | | | | **0.12 [0.35]** | **AB** | | | |
| *My. glareolus* | **0.23 [0.40]** | **A** | | | | **0.12 [0.39]** | **A** | | | | **-0.11 [0.21]** | **C** | | | |
| *Ap. flavicollis* | **-0.09 [0.24]** | **B** | | | | **-0.10 [0.20]** | **B** | | | | **-0.04 [0.19]** | **BC** | | | |
| *Ap. sylvaticus* | **-0.19 [0.28]** | **B** | | | | **-0.03 [0.27]** | **AB** | | | | **0.14 [0.37]** | **A** | | | |
| Habitat type | | | 2 | 0.67 | 0.51 | | | 2 | 2.60 | 0.08 | | | 2 | 0.71 | 0.49 |
| Forest | *ns* | *ns* | | | | -0.05 [0.29] | *ns* | | | | *ns* | *ns* | | | |
| Hedgerows | *ns* | *ns* | | | | 0.04 [0.30] | *ns* | | | | *ns* | *ns* | | | |
| Meadows | *ns* | *ns* | | | | -0.01 [0.28] | *ns* | | | | *ns* | *ns* | | | |
| Study Site | | | 1 | 1.61 | 0.21 | | | 1 | 2.65 | 0.10 | | | **1** | **4.03** | **0.05** |
| Boult-aux-Bois | *ns* | *ns* | | | | 0.03 [0.29] | *ns* | | | | **0.03 [0.29]** | **A** | | | |
| Briquenay | *ns* | *ns* | | | | -0.03 [0.29] | *ns* | | | | **-0.03 [0.24]** | **B** | | | |
| Year | | | 1 | 0.24 | 0.63 | | | 1 | < 0.01 | 0.97 | | | **1** | **4.65** | **0.032** |
| 2010 | *ns* | *ns* | | | | *ns* | *ns* | | | | **-0.04 [0.23]** | **A** | | | |
| 2011 | *ns* | *ns* | | | | *ns* | *ns* | | | | **0.03 [0.29]** | **B** | | | |
| Age group | | | **1** | **6.27** | **0.013** | | | 1 | 0.41 | 0.52 | | | 1 | 0.46 | 0.50 |
| Adult | **0.02 [0.29]** | **A** | | | | *ns* | *ns* | | | | *ns* | *ns* | | | |
| Juvenile | **-0.11 [0.26]** | **B** | | | | *ns* | *ns* | | | | *ns* | *ns* | | | |
| Sex | | | 1 | 0.36 | 0.55 | | | 1 | 0.87 | 0.35 | | | 1 | < 0.01 | 0.97 |
| Male | *ns* | *ns* | | | | *ns* | *ns* | | | | *ns* | *ns* | | | |
| Female | *ns* | *ns* | | | | *ns* | *ns* | | | | *ns* | *ns* | | | |

| | MCA Dim4 residual mean [sd] | group | df | *F* | p-value | MCA Dim5 residual mean [sd] | group | df | *F* | p-value | MCA Dim6 residual mean [sd] | group | df | *F* | p-value |
|---|---|---|---|---|---|---|---|---|---|---|---|---|---|---|---|
| Host species | | | **5** | **5.97** | **< 0.0001** | | | 5 | 1.44 | 0.21 | | | **5** | **3.67** | **0.0032** |
| *Ar. scherman* | **-0.09 [0.22]** | **B** | | | | *ns* | *ns* | | | | **-0.06 [0.20]** | **B** | | | |
| *Mi. agrestis* | **-0.01 [0.11]** | **AB** | | | | *ns* | *ns* | | | | **-0.11 [0.14]** | **B** | | | |
| *Mi. arvalis* | **0.15 [0.37]** | **A** | | | | *ns* | *ns* | | | | **0.10 [0.29]** | **A** | | | |
| *My. glareolus* | **-0.01 [0.37]** | **B** | | | | *ns* | *ns* | | | | **-0.05 [0.45]** | **AB** | | | |
| *Ap. flavicollis* | **-0.01 [0.20]** | **B** | | | | *ns* | *ns* | | | | **0.01 [0.15]** | **AB** | | | |
| *Ap. sylvaticus* | **-0.01 [0.20]** | **B** | | | | *ns* | *ns* | | | | **0.04 [0.14]** | **AB** | | | |
| Habitat type | | | **2** | **4.77** | **0.009** | | | 2 | 2.00 | 0.14 | | | 2 | 0.83 | 0.44 |
| Forest | **-0.07 [0.25]** | **B** | | | | *ns* | *ns* | | | | *ns* | *ns* | | | |
| Hedgerows | **0.05 [0.29]** | **A** | | | | *ns* | *ns* | | | | *ns* | *ns* | | | |
| Meadows | **-0.01 [0.27]** | **AB** | | | | *ns* | *ns* | | | | *ns* | *ns* | | | |
| Study Site | | | 1 | 1.83 | 0.18 | | | 1 | 0.32 | 0.57 | | | 1 | 0.29 | 0.59 |
| Boult-aux-Bois | *ns* | *ns* | | | | *ns* | *ns* | | | | *ns* | *ns* | | | |
| Briquenay | *ns* | *ns* | | | | *ns* | *ns* | | | | *ns* | *ns* | | | |
| Year | | | 1 | 3.42 | 0.065 | | | 1 | 0.87 | 0.35 | | | 1 | 1.92 | 0.17 |
| 2010 | 0.03 [0.27] | *ns* | | | | *ns* | *ns* | | | | 0.02 [0.31] | *ns* | | | |
| 2011 | -0.03 [0.27] | *ns* | | | | *ns* | *ns* | | | | -0.02 [0.22] | *ns* | | | |
| Age group | | | 1 | 0.40 | 0.053 | | | 1 | 0.80 | 0.37 | | | 1 | 0.21 | 0.64 |
| Adult | 0.01 [0.28] | *ns* | | | | *ns* | *ns* | | | | *ns* | *ns* | | | |
| Juvenile | -0.02 [0.17] | *ns* | | | | *ns* | *ns* | | | | *ns* | *ns* | | | |
| Sex | | | 1 | 0.23 | 0.63 | | | 1 | 0.51 | 0.47 | | | 1 | 0.01 | 0.94 |
| Male | *ns* | *ns* | | | | *ns* | *ns* | | | | *ns* | *ns* | | | |
| Female | *ns* | *ns* | | | | *ns* | *ns* | | | | *ns* | *ns* | | | |

| | MCA Dim7 residual mean [sd] | group | df | *F* | p-value |
|---|---|---|---|---|---|
| Host species | | | **5** | **3.49** | **0.0045** |
| *Ar. scherman* | **-0.06 [0.20]** | **B** | | | |
| *Mi. agrestis* | **-0.11 [0.14]** | **B** | | | |
| *Mi. arvalis* | **0.10 [0.29]** | **A** | | | |
| *My. glareolus* | **-0.05 [0.45]** | **AB** | | | |
| *Ap. flavicollis* | **0.01 [0.15]** | **AB** | | | |
| *Ap. sylvaticus* | **0.04 [0.14]** | **AB** | | | |
| Habitat type | | | 2 | 1.34 | 0.26 |
| Forest | *ns* | *ns* | | | |
| Hedgerows | *ns* | *ns* | | | |
| Meadows | *ns* | *ns* | | | |
| Study Site | | | 1 | 3.37 | 0.068 |
| Boult-aux-Bois | 0.03 [0.32] | *ns* | | | |
| Briquenay | -0.03 [0.16] | *ns* | | | |
| Year | | | **1** | **11.88** | **< 0.001** |
| 2010 | **0.06 [0.29]** | | | | |
| 2011 | **-0.05 [0.22]** | | | | |
| Age group | | | 1 | 0.57 | 0.45 |
| Adult | *ns* | *ns* | | | |
| Juvenile | *ns* | *ns* | | | |
| Sex | | | 1 | 1.21 | 0.27 |
| Male | *ns* | *ns* | | | |
| Female | *ns* | *ns* | | | |

Table S7: Association analyses between Myco1, Myco3, and Hantavirus exposure in *Mi. arvalis* and *My. glareolus*. (A) Association Screening (SCN) analyses between Myco1, Myco3, and Hantavirus exposure in *Mi. arvalis*, *My. glareolus* and both host species combined. The 95% confidence envelope against which observed frequencies were tested is given by its computationally-generated lower limit (LL) and upper limit (UL). Only co-exposure frequencies that are more frequent or more rare than expected by chance based on the confidence envelope are considered statistically significant (in bold). (B) Multiple logistic regression (GLM) models testing for intrinsic associations after first accounting for extrinsic factors structuring pathogen exposures within the two host species (based on the results of model selection provided in Figure S12). Bold rows indicate statistical significance at ($p < 0.05$).

A. Association Screening (SCN) analysis results

| Pathogen co-exposure status | | | *Mi. arvalis* adult hosts | | | | | | | *My. glareolus* adult hosts | | | | | | | Both *Mi. arvalis* and *My. glareolus* adult hosts | | | |
|---|---|---|---|---|---|---|---|---|---|---|---|---|---|---|---|---|---|---|---|---|
| Myco1 | Myco3 | Hanta-virus | Obs. Freq | LL | UL | Beyond 95% CI | SCN p-value | Direction | Obs. Freq | LL | UL | Beyond 95% CI | SCN p-value | Direction | Obs. Freq | LL | UL | Beyond 95% CI | SCN p-value | Direction |
| ✔ | ✔ | ✔ | 2 | 0 | 3 | No | 0.17 | - | 2 | 0 | 2 | No | 0.016 | - | 4 | 0 | 3 | Yes | 0.0084 | Frequent |
| ✔ | ✔ | - | 5 | 1 | 12 | No | 0.83 | - | 1 | 0 | 6 | No | 0.82 | - | 6 | 3 | 17 | No | 0.43 | - |
| ✔ | - | ✔ | 0 | 0 | 5 | No | 0.63 | - | 0 | 0 | 2 | No | 1 | - | 0 | 0 | 5 | No | 0.51 | - |
| - | ✔ | ✔ | 0 | 0 | 2 | No | 1 | - | 2 | 0 | 5 | No | 0.71 | - | 2 | 0 | 5 | No | 0.78 | - |
| ✔ | - | - | 15 | 7 | 21 | No | 0.95 | - | 3 | 0 | 9 | No | 0.71 | - | 15 | 7 | 21 | No | 0.91 | - |
| - | ✔ | - | 1 | 0 | 5 | No | 0.75 | - | 15 | 8 | 26 | No | 0.75 | - | 16 | 9 | 27 | No | 0.87 | - |
| - | - | ✔ | 0 | 0 | 2 | No | 1 | - | 0 | 0 | 7 | No | 0.19 | Rare | 0 | 0 | 7 | No | 0.13 | Rare |
| - | - | - | 4 | 0 | 8 | No | 0.83 | - | 33 | 20 | 40 | No | 0.48 | - | 37 | 22 | 45 | No | 0.56 | - |

B. Multiple logistic regression analysis

| | Myco1 infection in adults | | | Myco3 infection in adults | | | Hantavirus exposure in adults | | |
|---|---|---|---|---|---|---|---|---|---|
| | df | Adjusted $\chi 2$ | p-value | df | Adjusted $\chi 2$ | p-value | df | Adjusted $\chi 2$ | p-value |
| Extrinsic factors | | | | | | | | | |
| Year | - | - | - | 1 | 3.69 | 0.05 | - | - | - |
| Study Site | 1 | 3.46 | 0.06 | - | - | - | - | - | - |
| Host Sex | - | - | - | - | - | - | - | - | - |
| Host Habitat | - | - | - | - | - | - | **2** | **9.61** | **0.0082** |
| Host Species | **1** | **34.83** | **<0.0001** | - | - | - | - | - | - |
| Intrinsic factors | | | | | | | | | |
| Myco1 infection | - | - | - | 1 | 0.37 | 0.54 | **1** | **5.80** | **0.016** |
| Myco3 infection | 1 | 0.01 | 0.94 | - | - | - | - | - | - |
| Hantavirus exposure | 1 | 2.60 | 0.11 | **1** | **13.66** | **0.0002** | - | - | - |

#### Table S8. Association between Myco2 and Myco4 infections in Ap. sylvaticus.

(A) Association Screening (SCN) analysis between Myco2 and Myco4. The 95% confidence envelope against which observed frequencies were tested is given by its computationally-generated lower limit (LL) and upper limit (UL). Only co-exposure frequencies that are more frequent or more rare than expected by chance based on the confidence envelope are considered statistically significant. (B) Reciprocal multiple logistic regression analyses were performed to test for the association after first controlling for all extrinsic factors in the best model (based on the results of model selection provided in Figure S13). Bold factors indicate statistical significance at $p < 0.05$. Odds ratios are given for both trends ($p < 0.1$) and significant association results, and are adjusted for all other factors in the model.

A. Association Screening (SCN) analysis results

| Pathogen co-exposure status | | *Ap. sylvaticus* hosts | | | | | |
|---|---|---|---|---|---|---|---|
| Myco2 | Myco4 | Obs. Freq | LL | UL | Beyond 95% CI | SCN p-value | Direction |
| | | 5 | 0 | 8 | No | 0.38 | - |
| | - | 7 | 3 | 16 | No | 0.64 | - |
| - | | 13 | 7 | 23 | No | 0.68 | - |
| - | - | 45 | 33 | 52 | No | 0.73 | - |

B. Multiple logistic regression analysis results

| | **Myco2 infection** | | | **Myco4 infection** | | |
|---|---|---|---|---|---|---|
| *Residual df* = 62 | df | $\chi^2$ | *p*-value | df | $\chi^2$ | *p*-value |
| Extrinsic Factor | | | | | | |
| Habitat type | - | - | - | - | - | - |
| Study Site | - | - | - | - | - | - |
| Year | - | - | - | - | - | - |
| Age group | **1** | **4.07** | **0.044** | - | - | - |
| Sex | - | - | - | - | - | - |
| Intrinsic Factor | | | | | | |
| Myco4 infection | 1 | 2.79 | 0.12 | - | - | - |
| | OR 3.33 [0.80 – 13.73] | | | | | |
| Myco2 infection | - | - | - | 1 | 2.34 | 0.13 |
| | | | | OR 2.92 [0.73 – 11.43] | | |

Table S9: Association between Bartonella spp. infection and cowpox virus (CPXV) exposure across rodent species. (A) Independent Association Screening (SCN) analyses for each rodent host species. The 95% confidence envelope against which observed frequencies were tested is given by its computationally-generated lower limit (LL) and upper limit (UL). Only co-exposure frequencies that are more frequent or more rare than expected by chance based on the confidence envelope are considered statistically significant. (B) Reciprocal multiple logistic regression analyses were performed to test the association after controlling all other extrinsic factors in the best model (based on the results of model selection provided in Figure S14). Bold factors indicate statistical significance at $p < 0.05$. Odds ratios are given for both trends ($p < 0.1$) and significant association results, and are adjusted for all other factors in the model.

A. Association Screening (SCN) analysis results

| Pathogen co-exposure status | | *Ar. scherman* hosts | | | | | | *Mi. agrestis* hosts | | | | | |
|---|---|---|---|---|---|---|---|---|---|---|---|---|---|
| Bartonella infection | CPXV expsoure | Obs. Freq | LL | UL | Beyond 95% CI | SCN p-value | Direction | Obs. Freq | LL | UL | Beyond 95% CI | SCN p-value | Direction |
| ✔ | ✔ | 29 | 17 | 36 | No | 0.65 | - | 4 | 1 | 7 | No | 0.55 | - |
| ✔ | - | 27 | 20 | 39 | No | 0.66 | - | 2 | 0 | 5 | No | 0.71 | - |
| - | ✔ | 3 | 1 | 11 | No | 0.42 | - | 1 | 0 | 4 | No | 0.75 | - |
| - | - | 8 | 1 | 12 | No | 0.43 | - | 1 | 0 | 3 | No | 0.54 | - |
| | | *Mi. arvalis* hosts | | | | | | *My. glareolus* hosts | | | | | |
| Bartonella infection | CPXV expsoure | Obs. Freq | LL | UL | Beyond 95% CI | SCN p-value | Direction | Obs. Freq | LL | UL | Beyond 95% CI | SCN p-value | Direction |
| ✔ | ✔ | 5 | 1 | 11 | No | 0.58 | - | 26 | 17 | 36 | No | 0.57 | - |
| ✔ | - | 35 | 27 | 42 | No | 0.57 | - | 13 | 6 | 21 | No | 0.53 | - |
| - | ✔ | 1 | 0 | 4 | No | 0.59 | - | 13 | 5 | 21 | No | 0.52 | - |
| - | - | 6 | 1 | 12 | No | 0.59 | - | 6 | 1 | 13 | No | 0.6 | - |
| | | *Ap. flavicollis* hosts | | | | | | *Ap. sylvaticus* hosts | | | | | |
| Bartonella infection | CPXV expsoure | Obs. Freq | LL | UL | Beyond 95% CI | SCN p-value | Direction | Obs. Freq | LL | UL | Beyond 95% CI | SCN p-value | Direction |
| ✔ | ✔ | 5 | 1 | 12 | No | 0.79 | - | 7 | 2 | 13 | No | 0.51 | - |
| ✔ | - | 27 | 19 | 32 | No | 0.81 | - | 49 | 41 | 57 | No | 0.59 | - |
| - | ✔ | 2 | 0 | 3 | No | 0.37 | - | 1 | 0 | 5 | No | 0.71 | - |
| - | - | 2 | 0 | 8 | No | 0.73 | - | 9 | 3 | 16 | No | 0.54 | - |
| | | *R. norvegicus* hosts | | | | | | | | | | | |
| Bartonella infection | CPXV expsoure | Obs. Freq | LL | UL | Beyond 95% CI | SCN p-value | Direction | | | | | | |
| ✔ | ✔ | 0 | 0 | 2 | No | 1.00 | - | | | | | | |
| ✔ | - | 3 | 0 | 7 | No | 0.97 | - | | | | | | |
| - | ✔ | 4 | 0 | 8 | No | 0.98 | - | | | | | | |
| - | - | 21 | 16 | 26 | No | 0.66 | - | | | | | | |

Table S9 (cont.): Association between *Bartonella* spp. infection and cowpox virus (CPXV) exposure across rodent species.

B. Multiple logistic regression analysis results

| | *Ar. scherman* hosts (residual df = 61) | | | | | | *Mi. agrestis* hosts (residual df = 5) | | | | | |
|---|---|---|---|---|---|---|---|---|---|---|---|---|
| | *Bartonella* infection | | | CPXV exposure | | | *Bartonella* infection | | | CPXV exposure | | |
| | df | $\chi^2$ | *p*-value | df | $\chi^2$ | *p*-value | df | $\chi^2$ | *p*-value | df | $\chi^2$ | *p*-value |
| Extrinsic Factor | | | | | | | | | | | | |
| Habitat type | - | - | - | - | - | - | - | - | - | - | - | - |
| Study Site | - | - | - | - | - | - | - | - | - | - | - | - |
| Year | **1** | **7.29** | **0.0069** | **1** | **8.43** | **0.0037** | - | - | - | - | - | - |
| Age group | - | - | - | - | - | - | 1 | 3.82 | 0.051 | 1 | 2.77 | 0.096 |
| Sex | - | - | - | - | - | - | - | - | - | - | - | - |
| Intrinsic Factor | | | | | | | | | | | | |
| *Bartonella* infection | - | - | - | **1** | **5.07** | **0.024** | - | - | - | 1 | 0.74 | 0.39 |
| | | | | **OR: 5.35 [1.23-29.60]** | | | | | | | | |
| CPXV exposure | **1** | **5.07** | **0.024** | - | - | - | 1 | 0.74 | 0.39 | - | - | - |
| | **OR: 5.35 [1.23-29.60]** | | | | | | | | | | | |

| | *Mi. arvalis* hosts (residual df = 41) | | | | | | *My. glareolus* hosts (residual df = 50) | | | | | |
|---|---|---|---|---|---|---|---|---|---|---|---|---|
| | *Bartonella* infection | | | CPXV exposure | | | *Bartonella* infection | | | CPXV exposure | | |
| | df | $\chi^2$ | *p*-value | df | $\chi^2$ | *p*-value | df | $\chi^2$ | *p*-value | df | $\chi^2$ | *p*-value |
| Extrinsic Factor | | | | | | | | | | | | |
| Habitat type | - | - | - | - | - | - | **1** | **13.80** | **<0.001** | - | - | - |
| Study Site | **1** | **4.03** | **0.045** | 1 | 3.31 | 0.069 | **1** | **15.62** | **<0.001** | - | - | - |
| Year | **1** | 2.02 | 0.16 | - | - | - | - | - | - | **1** | **10.4** | **0.0013** |
| Age group | **1** | 2.46 | 0.12 | - | - | - | **1** | **10.21** | **0.0014** | - | - | - |
| Sex | - | - | - | - | - | - | - | - | - | - | - | - |
| Intrinsic Factor | | | | | | | | | | | | |
| *Bartonella* infection | - | - | - | 1 | 0.23 | 0.63 | - | - | - | 1 | 0.8 | 0.37 |
| CPXV exposure | 1 | 0.53 | 0.46 | - | - | - | 1 | 0.21 | 0.65 | - | - | - |

| | *Ap. flavicollis* hosts (residual df = 29) | | | | | | *Ap. sylvaticus* hosts (residual df = 58) | | | | | |
|---|---|---|---|---|---|---|---|---|---|---|---|---|
| | *Bartonella* infection | | | CPXV exposure | | | *Bartonella* infection | | | CPXV exposure | | |
| | df | $\chi^2$ | *p*-value | df | $\chi^2$ | *p*-value | df | $\chi^2$ | *p*-value | df | $\chi^2$ | *p*-value |
| Extrinsic Factor | | | | | | | | | | | | |
| Habitat type | - | - | - | - | - | - | - | - | - | - | - | - |
| Study Site | - | - | - | 1 | 2.96 | 0.085 | - | - | - | - | - | - |
| Year | 1 | 1.32 | 0.25 | - | - | - | 1 | 2.85 | 0.091 | - | - | - |
| Age group | - | - | - | 1 | 2.78 | 0.095 | 1 | 0.81 | 0.37 | 1 | 2.73 | 0.098 |
| Sex | 1 | 1.37 | 0.24 | 1 | 2.75 | 0.097 | 1 | 0.68 | 0.41 | 1 | 3.22 | 0.73 |
| Intrinsic Factor | | | | | | | | | | | | |
| *Bartonella* infection | - | - | - | 1 | 2.75 | 0.097 | - | - | - | 1 | 0.7 | 0.79 |
| | | | | OR: 0.07 [0.001-1.60] | | | | | | | | |
| CPXV exposure | 1 | 2.87 | 0.09 | - | - | - | 1 | 0.007 | 0.93 | - | - | - |
| | OR: 0.1 [0.0003-1.44] | | | | | | | | | | | |

| | *R. norvegicus* hosts (residual df = 25) | | | | | |
|---|---|---|---|---|---|---|
| | *Bartonella* infection | | | CPXV exposure | | |
| | df | $\chi^2$ | *p*-value | df | $\chi^2$ | *p*-value |
| Extrinsic Factor | | | | | | |
| Habitat type | - | - | - | - | - | - |
| Study Site | 1 | 2.24 | 0.13 | - | - | - |
| Year | - | - | - | - | - | - |
| Age group | - | - | - | 1 | 1.97 | 0.16 |
| Sex | - | - | - | - | - | - |
| Intrinsic Factor | | | | | | |
| *Bartonella* infection | - | - | - | 1 | 0.85 | 0.36 |
| CPXV exposure | 1 | 1.06 | 0.3 | - | - | - |

Table S10: Association between *Mycoplasma haemomuris* (MH) and *Myco. coccoides* (MC) infections in the four rodent species where they both circulate. (A) Independent Association Screening (SCN) analyses for each rodent host species. The 95% confidence envelope against which observed frequencies were tested is given by its computationally-generated lower limit (LL) and upper limit (UL). Only co-exposure frequencies that are more frequent or more rare than expected by chance based on the confidence envelope are considered statistically significant. (B) Reciprocal multiple logistic regression analyses were performed to test the association after controlling all other extrinsic factors in the best model (following results of model selection in Figure S15). Bold factors indicate statistical significance at $p < 0.05$. Odds ratios are given for both trends ($p < 0.1$) and significant association results, and are adjusted for all other factors in the model. The interaction term (in italics) is presented for completeness, but not included in the models.

#### A. Association Screening (SCN) analysis results

| Pathogen co-exposure status | | *My. glareolus* hosts | | | | | | *Ap. flavicollis* hosts | | | | | |
|---|---|---|---|---|---|---|---|---|---|---|---|---|---|
| MH | MC | Obs. Freq | LL | UL | Beyond 95% CI | SCN p-value | Direction | Obs. Freq | LL | UL | Beyond 95% CI | SCN p-value | Direction |
| ✔ | ✔ | 3 | 0 | 5 | No | 0.34 | - | 2 | 0 | 2 | No | 0.14 | - |
| ✔ | - | 27 | 20 | 38 | No | 0.79 | - | 6 | 3 | 14 | No | 0.70 | - |
| - | ✔ | 0 | 0 | 5 | No | 0.43 | - | 0 | 0 | 5 | No | 0.41 | - |
| - | - | 32 | 21 | 39 | No | 0.80 | - | 28 | 20 | 32 | No | 0.71 | - |
| | | *Ap. sylvaticus* hosts | | | | | | *R. norvegicus* hosts | | | | | |
| MH | MC | Obs. Freq | LL | UL | Beyond 95% CI | SCN p-value | Direction | Obs. Freq | LL | UL | Beyond 95% CI | SCN p-value | Direction |
| ✔ | ✔ | 6 | 0 | 8 | No | 0.23 | - | 10 | 4 | 15 | No | 0.51 | - |
| ✔ | - | 7 | 3 | 17 | No | 0.47 | - | 19 | 13 | 25 | No | 0.63 | - |
| - | ✔ | 12 | 7 | 23 | No | 0.51 | - | 0 | 0 | 2 | No | 1.00 | - |
| - | - | 45 | 33 | 52 | No | 0.59 | - | 1 | 0 | 3 | No | 0.98 | - |

#### B. Multiple logistic regression analysis results

| | ***Myco. haemomuris* (MH) infection** | | | ***Myco. coccoides* (MC) infection** | | |
|---|---|---|---|---|---|---|
| *Residual df = 186* | df | $\chi^2$ | p-value | df | $\chi^2$ | p-value |
| Extrinsic Factor | | | | | | |
| Host species | **2** | **26.57** | **<0.0001** | **2** | **25.00** | **<0.0001** |
| Habitat type | 1 | 1.28 | 0.26 | - | - | - |
| Study Site | **1** | **6.63** | **0.010** | - | - | - |
| Year | 1 | 3.08 | 0.079 | - | - | - |
| Age group | - | - | - | - | - | - |
| Sex | - | - | - | 1 | 2.35 | 0.13 |
| Intrinsic Factor | | | | | | |
| MH infection | - | - | - | **1** | **11.05** | **<0.001** |
| | | | | | **OR 6.87 [2.18-23.98]** | |
| MC infection | **1** | **8.67** | **0.0032** | - | - | - |
| | | **OR 5.71 [1.78-20.21]** | | | | |
| *Host species x [MH or MC] infection* | *3* | *3.53* | *0.32* | *3* | *3.30* | *0.34* |

#### Table S11: Association between *Bartonella* spp. and hemotropic *Mycoplasma* spp. infections across rodent species.

(A) Independent Association Screening (SCN) analyses for each rodent host species. The 95% confidence envelope against which observed frequencies were tested is given by its computationally-generated lower limit (LL) and upper limit (UL). Only co-exposure frequencies that are more frequent or more rare than expected by chance based on the confidence envelope are considered statistically significant. Reciprocal multiple logistic regression analyses were performed to test the association after controlling all other extrinsic factors in the model, performed on each individual host species with sufficient variation in the two infections (C) after a global model suggested association may vary by species (B). Bold factors indicate statistical significance at $p < 0.05$. Odds ratios are given for both trends ($p < 0.1$) and significant association results, and are adjusted for all other factors in the model.

A. Association Screening (SCN) analysis results

| Pathogen co-exposure status | | *Ar. scherman* hosts | | | | | | *Mi. agrestis* hosts | | | | | |
|---|---|---|---|---|---|---|---|---|---|---|---|---|---|
| Bartonella infection | HMyco infection | Obs. Freq | LL | UL | Beyond 95% CI | SCN p-value | Direction | Obs. Freq | LL | UL | Beyond 95% CI | SCN p-value | Direction |
| ✔ | ✔ | 1 | 0 | 3 | No | 0.56 | - | 3 | 0 | 6 | No | 0.62 | - |
| ✔ | - | 56 | 48 | 63 | No | 0.58 | - | 3 | 0 | 6 | No | 0.64 | - |
| - | ✔ | 0 | 0 | 2 | No | 1.00 | - | 1 | 0 | 3 | No | 0.65 | - |
| - | - | 13 | 6 | 21 | No | 0.53 | - | 1 | 0 | 3 | No | 0.67 | - |
| | | *Mi. arvalis* hosts | | | | | | *My. glareolus* hosts | | | | | |
| Bartonella infection | HMyco infection | Obs. Freq | LL | UL | Beyond 95% CI | SCN p-value | Direction | Obs. Freq | LL | UL | Beyond 95% CI | SCN p-value | Direction |
| ✔ | ✔ | 38 | 29 | 44 | No | 0.89 | - | 15 | 12 | 29 | No | 0.19 | Rare |
| ✔ | - | 5 | 1 | 12 | No | 0.89 | - | 27 | 12 | 31 | No | 0.21 | Frequent |
| - | ✔ | 5 | 1 | 12 | No | 0.86 | - | 15 | 3 | 17 | No | 0.095 | Frequent |
| - | - | 2 | 0 | 4 | No | 0.52 | - | 5 | 4 | 18 | No | 0.078 | Rare |
| | | *Ap. flavicollis* hosts | | | | | | *Ap. sylvaticus* hosts | | | | | |
| Bartonella infection | HMyco infection | Obs. Freq | LL | UL | Beyond 95% CI | SCN p-value | Direction | Obs. Freq | LL | UL | Beyond 95% CI | SCN p-value | Direction |
| ✔ | ✔ | 6 | 2 | 13 | No | 0.83 | - | 21 | 12 | 31 | No | 0.54 | - |
| ✔ | - | 26 | 18 | 31 | No | 0.85 | - | 38 | 28 | 48 | No | 0.54 | - |
| - | ✔ | 2 | 0 | 4 | No | 0.45 | - | 4 | 0 | 9 | No | 0.55 | - |
| - | - | 2 | 0 | 8 | No | 0.78 | - | 7 | 2 | 14 | No | 0.57 | - |
| | | *R. norvegicus* hosts | | | | | | | | | | | |
| Bartonella infection | HMyco infection | Obs. Freq | LL | UL | Beyond 95% CI | SCN p-value | Direction | | | | | | |
| ✔ | ✔ | 3 | 0 | 7 | No | 0.57 | - | | | | | | |
| ✔ | - | 0 | 0 | 1 | No | 1.00 | - | | | | | | |
| - | ✔ | 26 | 22 | 29 | No | 0.65 | - | | | | | | |
| - | - | 1 | 0 | 3 | No | 0.59 | - | | | | | | |

Table S11 (cont.): Association between *Bartonella* spp. and hemotropic *Mycoplasma* spp. infections across rodent species.

B. Multiple logistic regression analysis results (global model)

| *Residual df = 304* | ***Bartonella* spp. infection** | | | **Hemotropic *Mycoplasma* spp. infection (HM)** | | |
|---|---|---|---|---|---|---|
| | df | $\chi^2$ | *p*-value | df | $\chi^2$ | *p*-value |
| Extrinsic Factor | | | | | | |
| Host species | 4 | 4.09 | 0.39 | 4 | 4.62 | 0.33 |
| Habitat type | **2** | **10.03** | **0.0066** | 2 | 1.42 | 0.23 |
| Study Site | - | - | - | **1** | **8.34** | **0.0039** |
| Year | **1** | **8.57** | **0.0034** | 1 | 2.34 | 0.13 |
| Age group | - | - | - | - | - | - |
| Sex | - | - | - | - | - | - |
| Intrinsic Factor | | | | | | |
| Bartonella infection | - | - | - | 1 | 0.083 | 0.77 |
| HM infection | 1 | 0.09 | 0.77 | - | - | - |
| Interaction Term | | | | | | |
| Host species x [Bartonella or HM] infection | 4 | 8.74 | 0.068 | 4 | 4.55 | 0.34 |

Table S11 (cont.): Association between *Bartonella* spp. and hemotropic *Mycoplasma* spp. infections across rodent species.

C. Multiple logistic regression analysis results (individual species models)

| | *Ar. scherman* hosts (n = 69) | | | | | | *Mi. agrestis* hosts (n=8) | | | | | |
|---|---|---|---|---|---|---|---|---|---|---|---|---|
| | *Bartonella* infection | | | Hemotropic *Mycoplasma* infection | | | *Bartonella* infection | | | Hemotropic *Mycoplasma* infection | | |
| | df | $\chi^2$ | *p*-value | df | $\chi^2$ | *p*-value | df | $\chi^2$ | *p*-value | df | $\chi^2$ | *p*-value |
| Extrinsic Factor | | | | | | | | | | | | |
| Habitat type | only 1 *Mycoplasma* infection and only 1 uninfected by *Bartonella* | | | | | | - | - | - | 1 | 1.73 | 0.19 |
| Study Site | | | | | | | - | - | - | - | - | - |
| Year | | | | | | | - | - | - | - | - | - |
| Age group | | | | | | | 1 | 4.50 | 0.034 | | | |
| Sex | | | | | | | - | - | - | 1 | 1.73 | 0.19 |
| Intrinsic Factor | | | | | | | | | | | | |
| *Bartonella* infection | | | | | | | - | - | - | 1 | 0.00 | 1.00 |
| HM infection | | | | | | | 1 | 1.24 | 0.26 | - | - | - |

| | *Mi. arvalis* hosts (n = 44) Hedgerows excluded b/c all infected w/HM | | | | | | *My. glareolus* hosts (n = 57) | | | | | |
|---|---|---|---|---|---|---|---|---|---|---|---|---|
| | *Bartonella* infection | | | Hemotropic *Mycoplasma* infection | | | *Bartonella* infection | | | Hemotropic *Mycoplasma* infection | | |
| | df | $\chi^2$ | *p*-value | df | $\chi^2$ | *p*-value | df | $\chi^2$ | *p*-value | df | $\chi^2$ | *p*-value |
| Extrinsic Factor | | | | | | | | | | | | |
| Habitat type | - | - | - | - | - | - | **1** | **11.81** | **<0.001** | - | - | - |
| Study Site | 1 | 2.68 | 0.11 | 1 | 3.82 | 0.051 | **1** | **6.93** | **0.0084** | 1 | 3.30 | 0.069 |
| Year | 1 | 1.03 | 0.31 | 1 | 1.99 | 0.16 | 1 | 2.79 | 0.095 | **1** | **5.88** | **0.015** |
| Age group | 1 | 2.00 | 0.16 | - | - | - | **1** | **7.82** | **0.0052** | - | - | - |
| Sex | - | - | - | - | - | - | - | - | - | - | - | - |
| Intrinsic Factor | | | | | | | | | | | | |
| *Bartonella* infection | - | - | - | 1 | 0.31 | 0.58 | - | - | - | **1** | **6.59** | **0.01** |
| | | | | | | | | | | **OR: 0.16 [0.03 - 0.66]** | | |
| HM infection | 1 | 0.62 | 0.43 | - | - | - | **1** | **4.14** | **0.042** | - | - | - |
| | | | | | | | **OR: 0.17 [0.02 - 0.94]** | | | | | |

| | *Ap. flavicollis* hosts (n=36) | | | | | | *Ap. sylvaticus* hosts (n = 65) | | | | | |
|---|---|---|---|---|---|---|---|---|---|---|---|---|
| | *Bartonella* infection | | | Hemotropic *Mycoplasma* infection | | | *Bartonella* infection | | | Hemotropic *Mycoplasma* infection | | |
| | df | $\chi^2$ | *p*-value | df | $\chi^2$ | *p*-value | df | $\chi^2$ | *p*-value | df | $\chi^2$ | *p*-value |
| Extrinsic Factor | | | | | | | | | | | | |
| Habitat type | - | - | - | 1 | 0.48 | 0.49 | - | - | - | - | - | - |
| Study Site | - | - | - | 1 | 0.31 | 0.58 | - | - | - | - | - | - |
| Year | 1 | 2.03 | 0.15 | 1 | 0.42 | 0.51 | 1 | 3.15 | 0.076 | - | - | - |
| Age group | - | - | - | 1 | 0.24 | 0.62 | 1 | 0.89 | 0.35 | - | - | - |
| Sex | - | - | - | 1 | 1.33 | 0.25 | 1 | 0.74 | 0.39 | - | - | - |
| Intrinsic Factor | | | | | | | | | | | | |
| *Bartonella* infection | - | - | - | 1 | 1.73 | 0.19 | - | - | - | 1 | 1.71 | 0.19 |
| HM infection | 1 | 1.27 | 0.26 | - | - | - | 1 | 0.28 | 0.60 | - | - | - |

| | *R. norvegicus* hosts (residual n = 28) | |
|---|---|---|
| | *Bartonella* infection | Hemotropic *Mycoplasma* infection |
| Extrinsic Factor | only 1 animal uninfected by HM | |
| Habitat type | | |
| Study Site | | |
| Year | | |
| Age group | | |
| Sex | | |
| Intrinsic Factor | | |
| *Bartonella* infection | | |
| HM infection | | |

#### Figures

##### Figure S1. Landscape of trapping locations for focal host species.

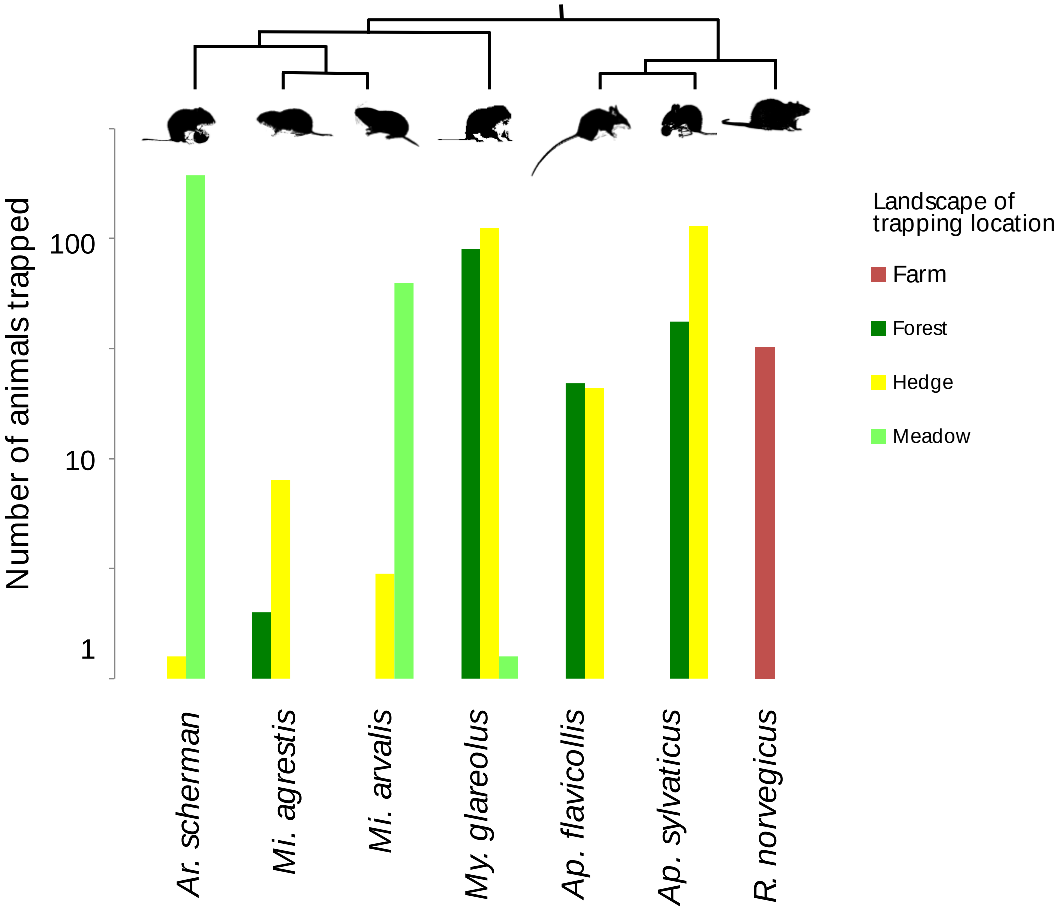

Figure S2. Host species demographic distributions (sex and age class). (A) Sex distribution in each focal host species. (B) Age class distribution in each focal host species. Adult = only sexually mature adults, juvenile = juveniles and non-breeding sub-adults.

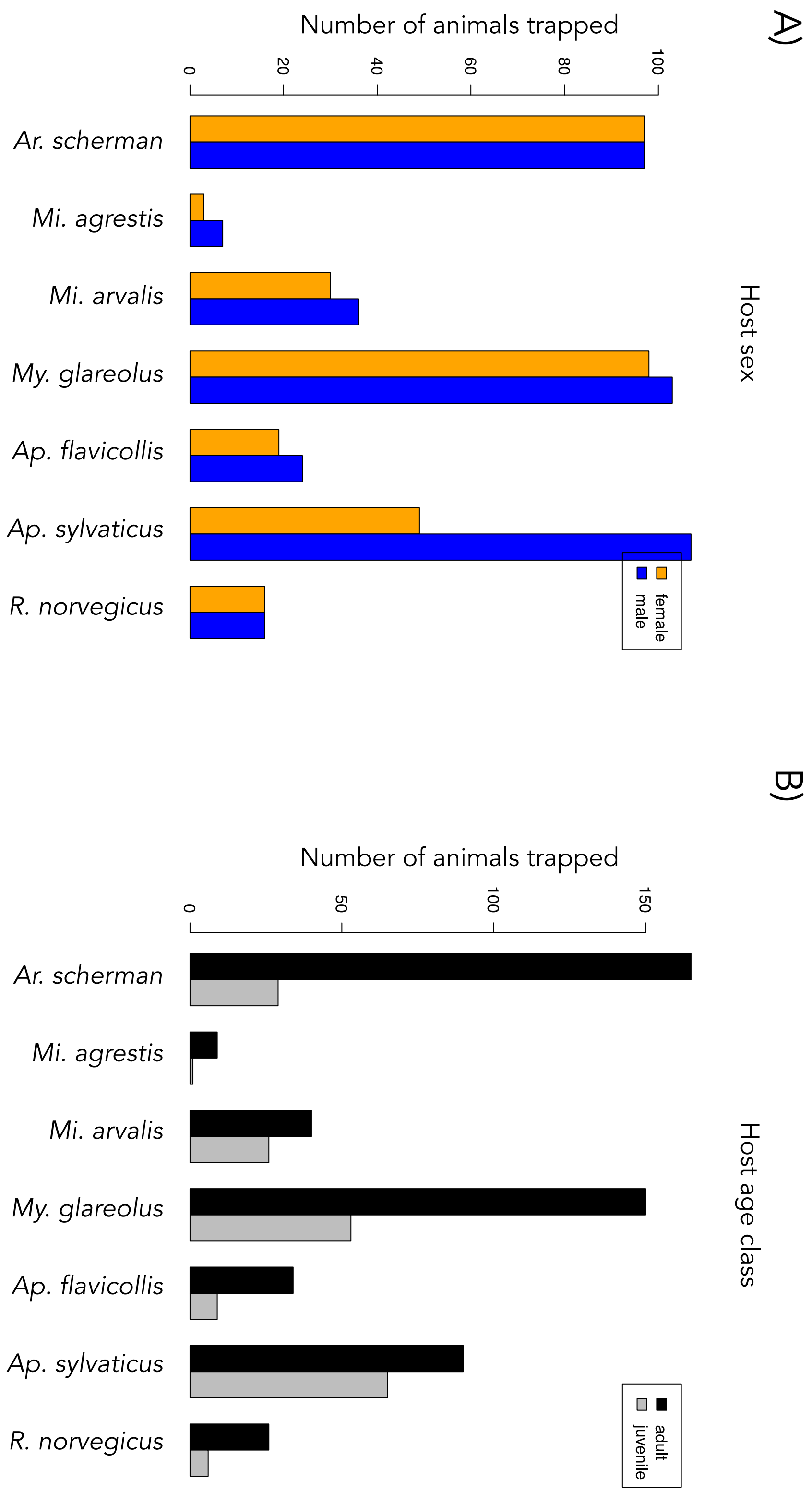

Figure S3. Data filtering process for 16S meta-barcoding identification of pathogenic bacteria.

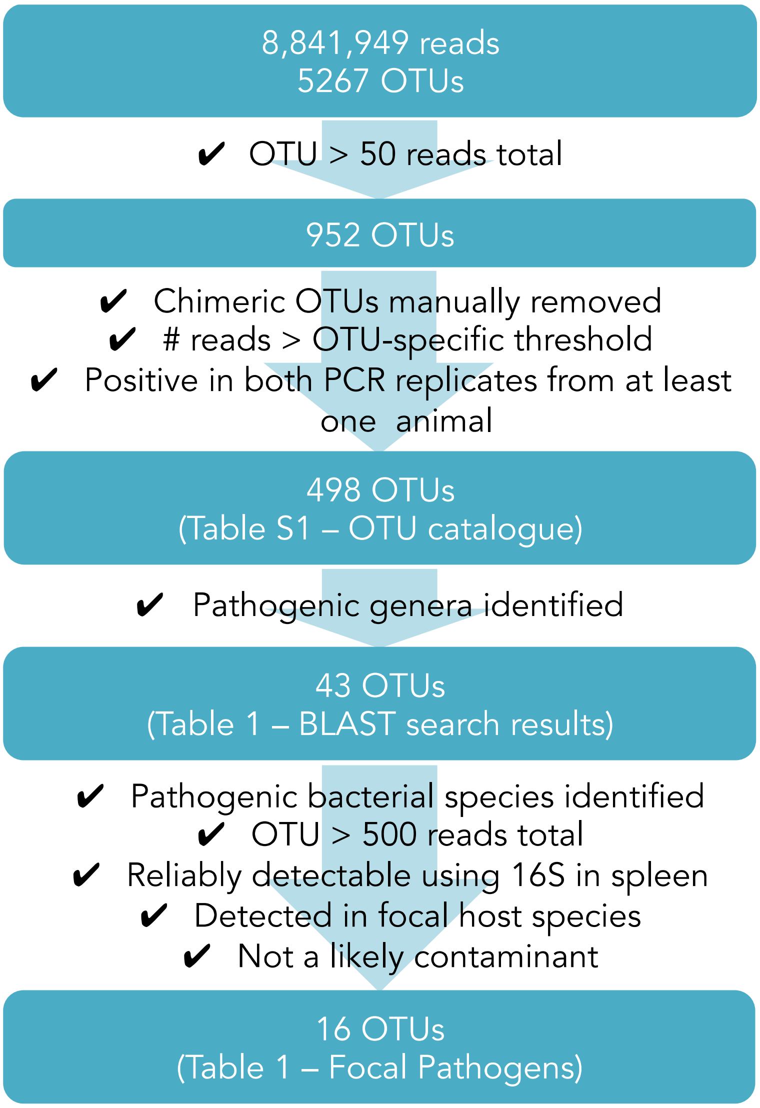

#### Figure S4. Phylogeny of Mycoplasma species and OTUs.

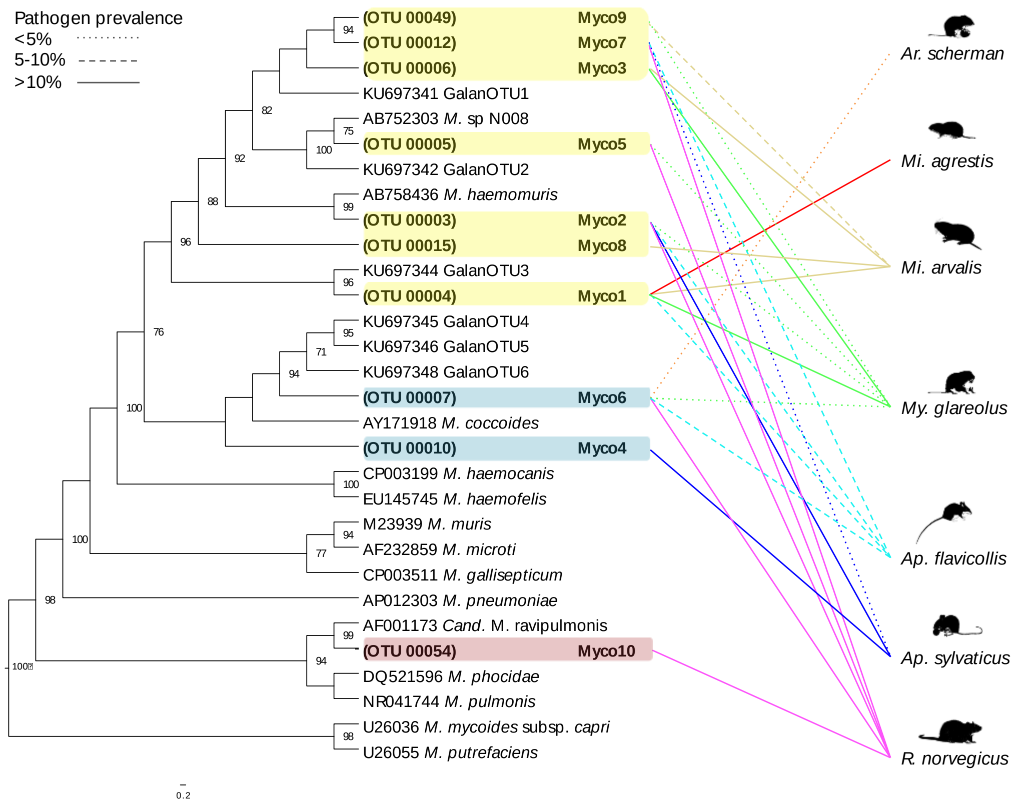

Figure S5. Results of Akaike Information Criterion (AIC)-based model selection analysis of the effect of extrinsic factors on pathogen diversity. Plotted here are the coefficients for each factor level remaining in the best model after model selection. The best model (including age, site, and host species) had an AIC weight of 23%.

| model | aic | weights |
|---|---|---|
| divGRP ~ 1 + AGE + HABITAT + HostSPP | 101.8469 | 0.22927215 |
| divGRP ~ 1 + YEAR + AGE + HABITAT + HostSPP | 102.4115 | 0.17288341 |
| divGRP ~ 1 + AGE + SEX + HABITAT + HostSPP | 102.5739 | 0.15939549 |
| divGRP ~ 1 + YEAR + AGE + SEX + HABITAT + HostSPP | 103.1328 | 0.12053366 |
| divGRP ~ 1 + SITE + AGE + HABITAT + HostSPP | 103.6340 | 0.09381694 |

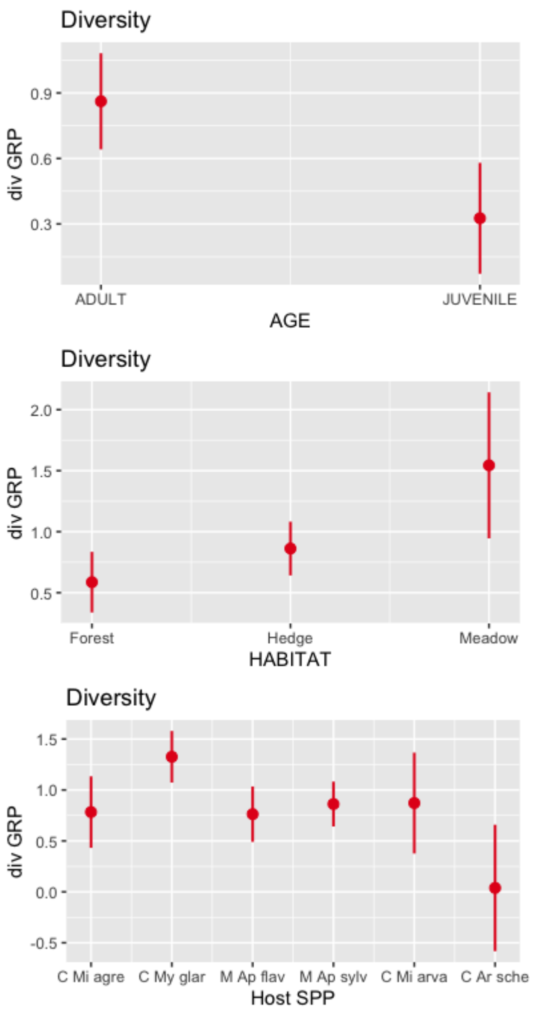

**Figure S6. Cook's distances for co-infection outliers.** Cook's distance was used to measure the impact of removing each datapoint from the regression of the relationship between within-host co-infection rates and the number of co-circulating bacterial (A) or both viral and bacterial (B) pathogens in each host species (given by its Shannon index; Figures 3C and 3D, respectively, in the main text). Higher Cook's distance indicate greater likelihood of outlier status, with a more liberal rule of thumb stating outliers are those with Cook's *D* of over 4/n (where n = the number of observations; here: 0.57) and more conservative stating a cutoff of 4 x mean Cook's *D* (horizontal red line).

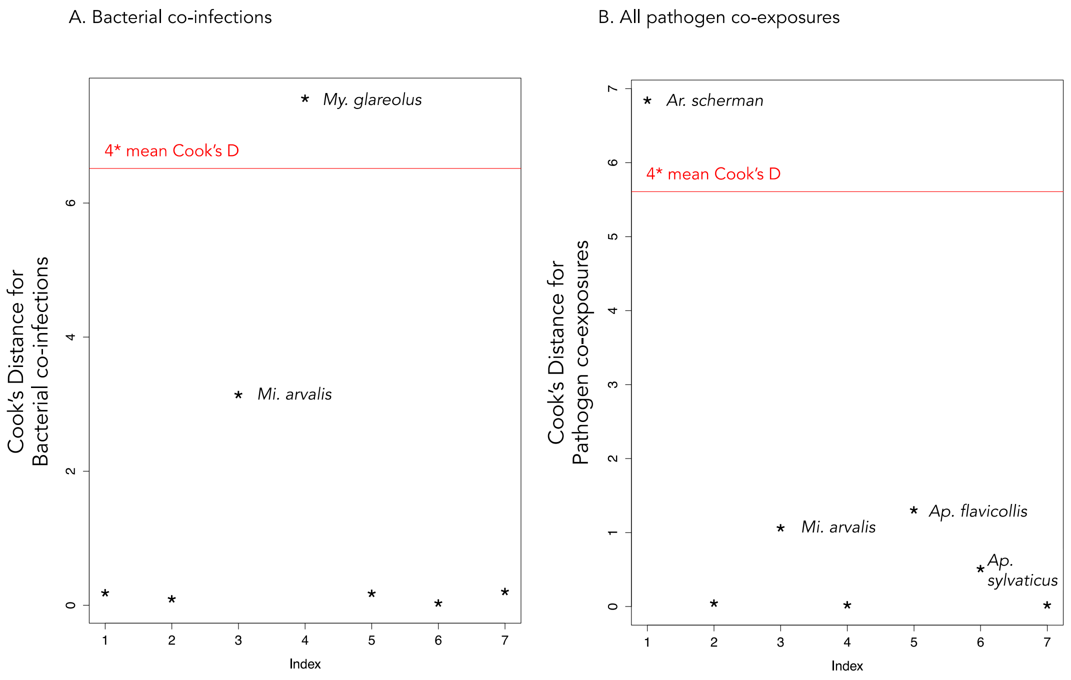

Figure S7. Results of multiple correspondence analysis (MCA) for pathogen community composition in rodents (including *R. norvegicus*) are described by (A) the contribution of each dimension to the overall variance in the data, (B) variable correlations with the first two dimensions of the MCA, and (C) variable contributions to each orthogonal MCA dimension. Horizontal line in (A) represents the per cent variance expected due to chance (100/14) = 7.14%.

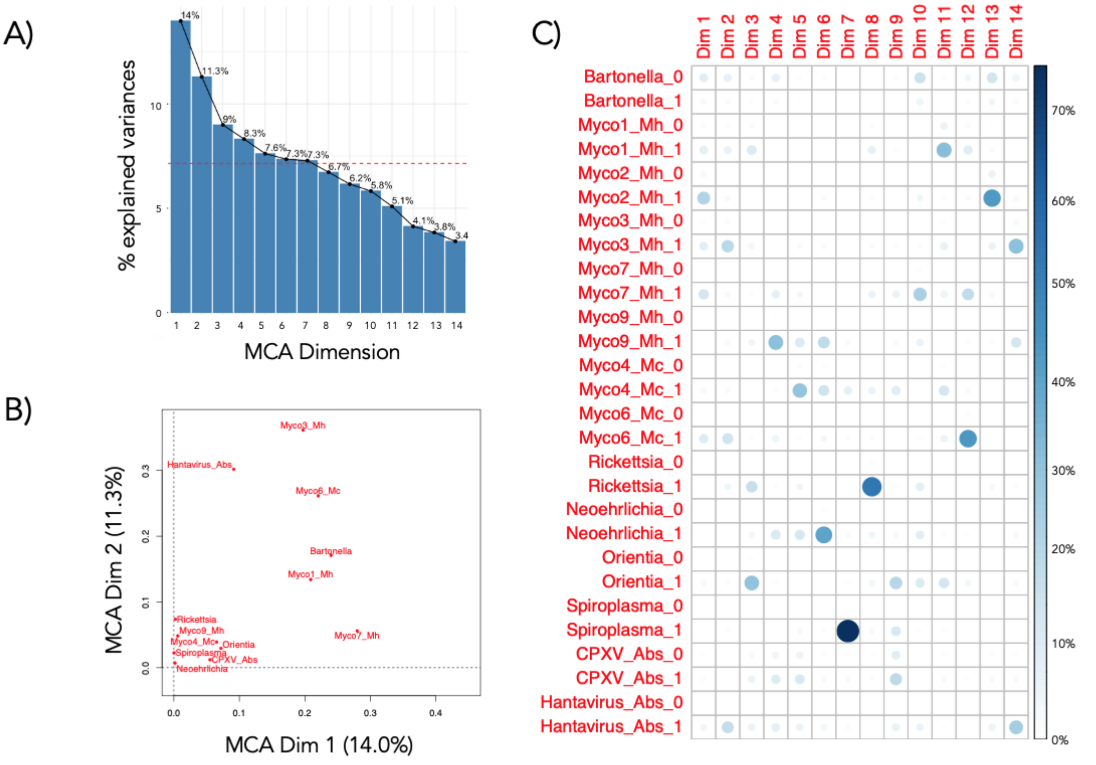

**Figure S8. Variance in pathogen community structure among (A) host species and (B) habitats including *R. norvegicus* from farms.** Pathogen community structure is represented by plotting the mean values for the first and second dimensions described by multiple correspondence analysis (MCA). MCA Dim1 and MCA Dim 2 collectively accounted for 25.3% of total variance in the data. Ellipses include 95% of individual values for each factor group.

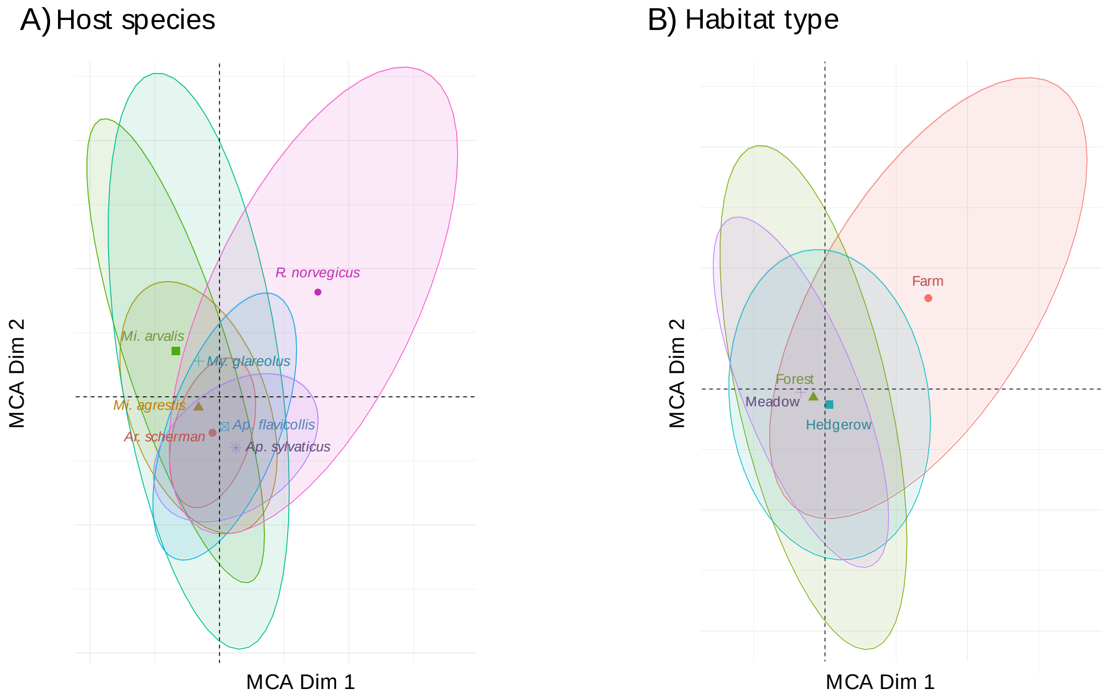

Figure S9. Data used for pathogen community composition multiple correspondence analysis (MCA).

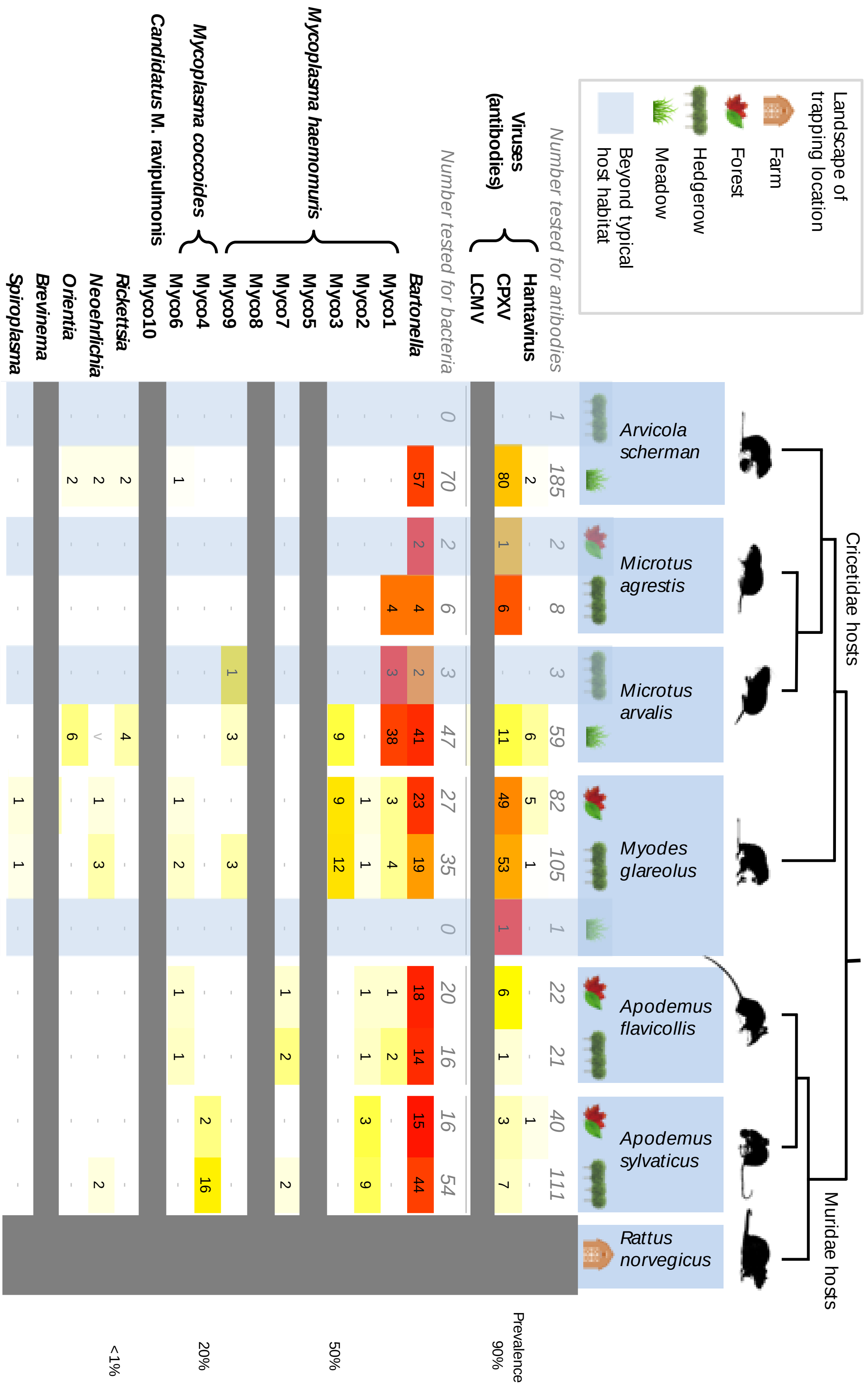

Figure S10: Pathogen community structure among extrinsic factors (A) host species (B) host age groups and (C) habitat types, represented by plotting the mean values for the first and second dimensions described by multiple correspondence analysis (MCA), excluding *Rattus norvegicus* hosts from farms. MCA Dim1 and MCA Dim 2 collectively accounted for 23% of total variance in the data. Ellipses include 95% of individual values for each factor group.

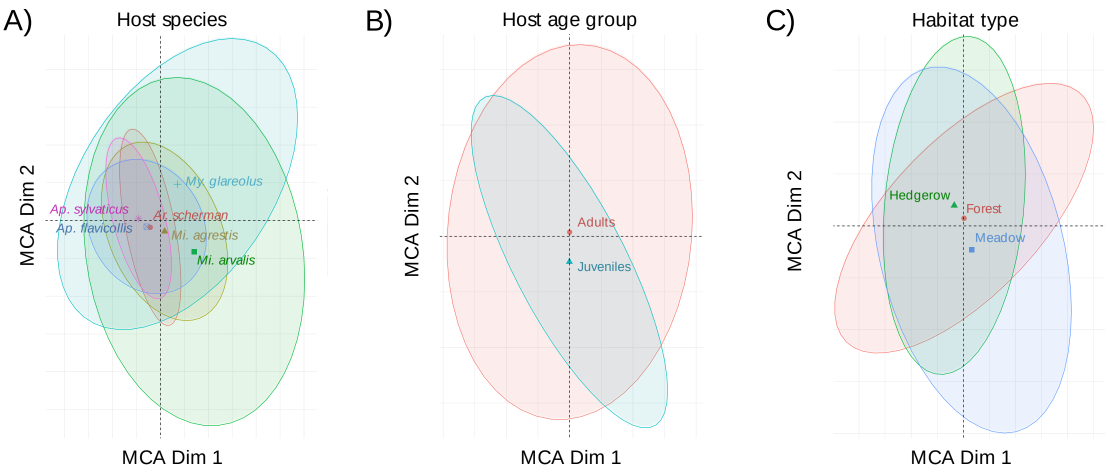

Figure S11. Association Screening (SCN) analysis results for Myco1, Myco3, and Hantavirus antibodies in *Mi. arvalis* and *My. glareolus* host species. The 95% confidence envelope is delimited by the lower limit (blue line) and the upper limit (red line). Observed frequencies of each co-occurrence status are considered statistically significant when they fall above (more frequent than expected) or below (more rare than expected) the confidence envelope.

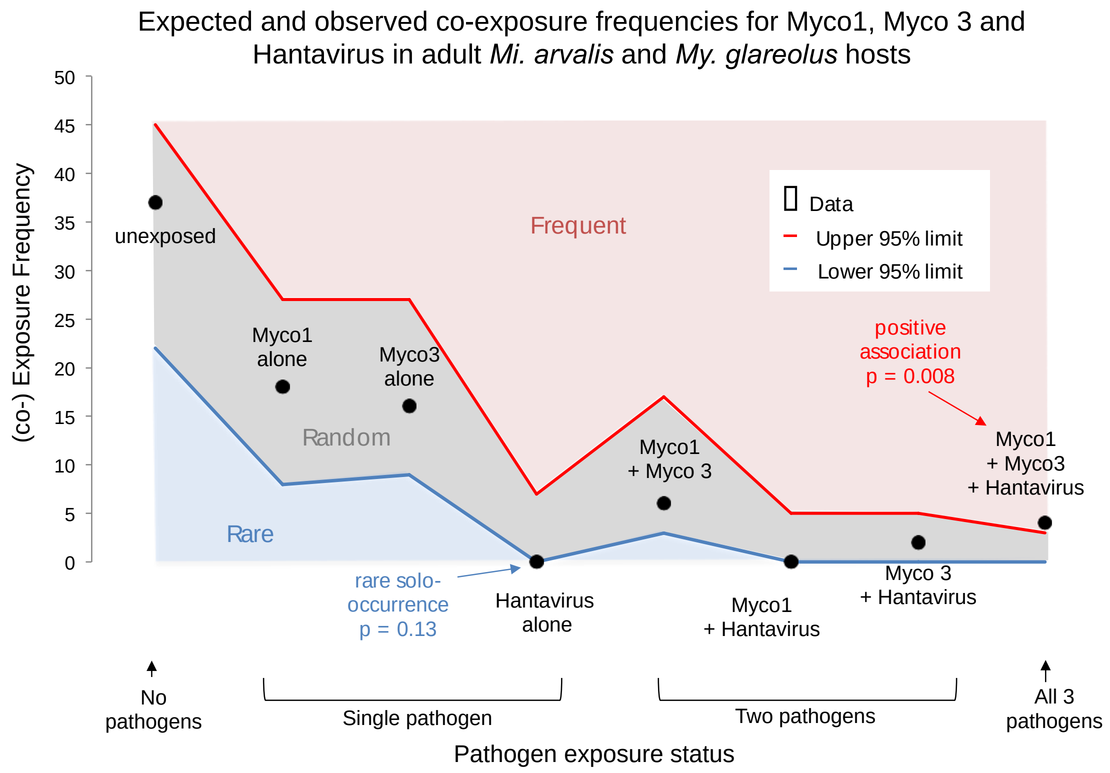

**Figure S12. Results of Akaike Information Criterion (AIC)-based model selection analysis of associations between occurrence of Myco1, Myco3 and hantavirus exposure.** Plotted here are the coefficients of marginal effects for each factor level remaining in the best model after model selection, as well as a table showing the AICc weights for the top-ranking models.

Myco 1 occurrence:

| model | aicc | weights |
|---|---|---|
| Myco1 ~ 1 + HostSPP + SITE + VirusHV | 63.75011 | 0.3290954 |
| Myco1 ~ 1 + HostSPP + SITE | 64.97544 | 0.1783383 |
| Myco1 ~ 1 + HostSPP + VirusHV | 64.99670 | 0.1764529 |
| Myco1 ~ 1 + HostSPP + SITE + VirusHV + Myco3 | 66.01408 | 0.1060980 |

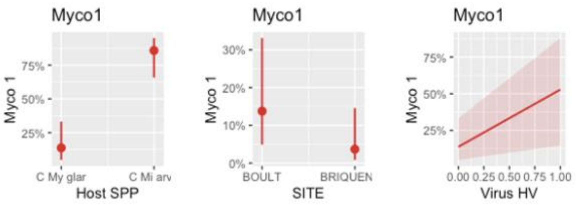

Myco 3 occurrence:

| model | aicc | weights |
|---|---|---|
| Myco3 ~ 1 + YEAR + VirusHV | 94.17075 | 0.13113324 |
| Myco3 ~ 1 + YEAR + SEX + VirusHV | 95.40717 | 0.07066885 |
| Myco3 ~ 1 + YEAR + HostSPP + VirusHV | 95.52461 | 0.06663879 |
| Myco3 ~ 1 + VirusHV | 95.60722 | 0.06394216 |
| Myco3 ~ 1 + YEAR + VirusHV + Myco1 | 96.01492 | 0.05215035 |

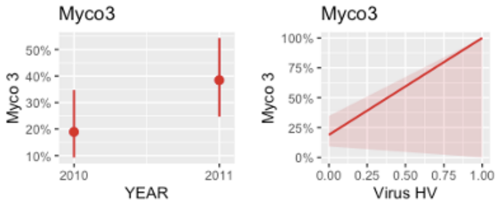

Anti-hantavirus antibody occurrence:

| model | aicc | weights |
|---|---|---|
| VirusHV ~ 1 + HABITAT + Myco1 | 38.98316 | 0.14414770 |
| VirusHV ~ 1 + YEAR + HABITAT + Myco1 | 39.36507 | 0.11909074 |
| VirusHV ~ 1 + SITE + HABITAT + Myco1 | 39.83984 | 0.09392533 |
| VirusHV ~ 1 + YEAR + SITE + HABITAT + Myco1 | 40.20726 | 0.07816227 |

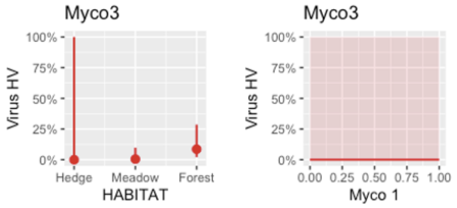

##### Figure S13. Results of Akaike Information Criterion (AIC)-based model selection analysis of associations between occurrence of Myco2 and Myco4.

Plotted here are the coefficients of marginal effects for each factor level remaining in the best model after model selection, as well as a table showing the AICc weights for the top-ranking models.

Myco 2 occurrence:

| model | aicc | weights |
|---|---|---|
| Myco2 ~ 1 + AGE + Myco4 | 59.09240 | 0.11542962 |
| Myco2 ~ 1 + AGE | 59.67962 | 0.08606051 |
| Myco2 ~ 1 + AGE + SITE + Myco4 | 60.94980 | 0.04560230 |
| Myco2 ~ 1 + Myco4 | 60.96060 | 0.04535680 |

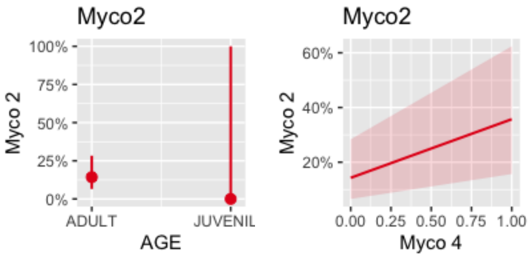

Myco 4 occurrence:

| model | aicc | weights |
|---|---|---|
| Myco4 ~ 1 + Myco2 | 76.56002 | 0.07727090 |
| Myco4 ~ 1 | 76.76929 | 0.06959445 |
| Myco4 ~ 1 + YEAR + Myco2 | 77.62886 | 0.04528143 |
| Myco4 ~ 1 + HABITAT + Myco2 | 77.72327 | 0.04319357 |
| Myco4 ~ 1 + YEAR | 77.79178 | 0.04173912 |
| Myco4 ~ 1 + HABITAT | 77.83586 | 0.04082918 |
| Myco4 ~ 1 + AGE + Myco2 | 78.04512 | 0.03677317 |
| Myco4 ~ 1 + SEX + Myco2 | 78.11051 | 0.03559025 |
| Myco4 ~ 1 + SEX | 78.25622 | 0.03308957 |

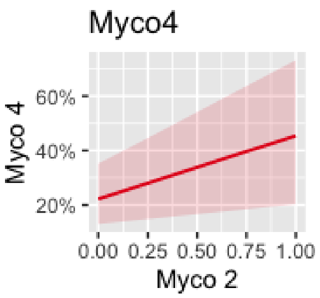

Figure S14. Results of Akaike Information Criterion (AIC)-based model selection analysis of associations between occurrence of Bartonella and CPXV exposure. Plotted here are the coefficients of marginal effects for each factor level remaining in the best model after model selection, as well as a table showing the AICc weights for the top-ranking models.

*Ar. scherman* hosts

| model | aicc | weights |
|---|---|---|
| Bartonella ~ 1 + YEAR + VirusCPXV | 56.62781 | 0.2894498 |
| Bartonella ~ 1 + YEAR + SEX + VirusCPXV | 58.56816 | 0.1097063 |

| model | aicc | weights |
|---|---|---|
| VirusCPXV ~ 1 + YEAR + Bartonella | 88.40767 | 0.2734596 |
| VirusCPXV ~ 1 + YEAR + AGE + Bartonella | 89.82668 | 0.1345116 |
| VirusCPXV ~ 1 + YEAR + SITE + Bartonella | 90.26341 | 0.1081247 |

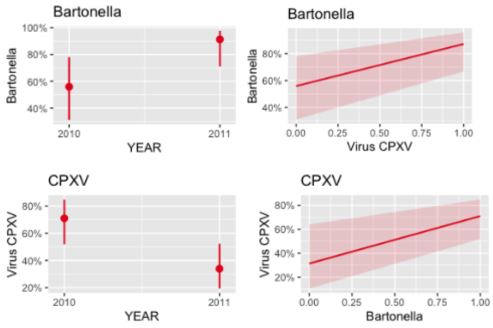

*Mi. agrestis* hosts

| model | aicc | weights |
|---|---|---|
| Bartonella ~ 1 | 11.66403 | 0.4936953 |
| Bartonella ~ 1 + AGE | 12.14163 | 0.3888209 |

| model | aicc | weights |
|---|---|---|
| VirusCPXV ~ 1 | 13.25168 | 0.3888426 |
| VirusCPXV ~ 1 + AGE | 14.77577 | 0.1814766 |
| VirusCPXV ~ 1 + SITE | 14.77577 | 0.1814766 |

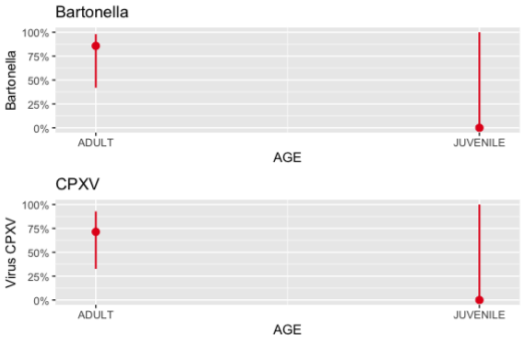

*Mi. arvalis* hosts

| model | aicc | weights |
|---|---|---|
| Bartonella ~ 1 + AGE | 40.81747 | 0.10910679 |
| Bartonella ~ 1 + AGE + SITE | 41.10473 | 0.09450909 |
| Bartonella ~ 1 + YEAR + SITE | 41.55053 | 0.07562595 |
| Bartonella ~ 1 | 41.64967 | 0.07196864 |
| Bartonella ~ 1 + YEAR + AGE + SITE | 41.68289 | 0.07078295 |
| Bartonella ~ 1 + SITE | 41.83808 | 0.06549806 |

| model | aicc | weights |
|---|---|---|
| VirusCPXV ~ 1 + SITE | 37.07233 | 0.17026596 |
| VirusCPXV ~ 1 | 37.98874 | 0.10767935 |
| VirusCPXV ~ 1 + SITE + SEX | 38.67292 | 0.07648314 |

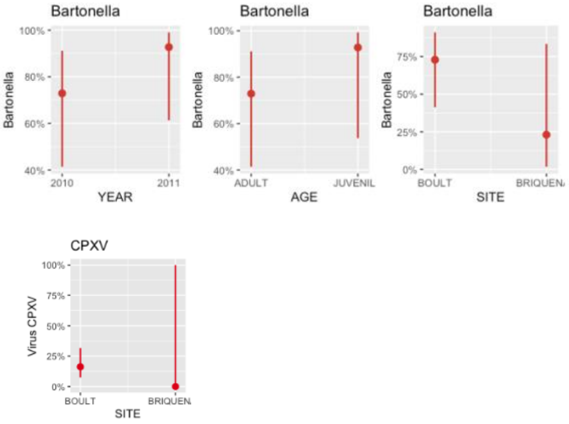

#### *My. glareolus* hosts

| model | aicc | weights |
|---|---|---|
| Bartonella ~ 1 + AGE + SITE + HABITAT | 52.11549 | 0.2865622 |
| Bartonella ~ 1 + YEAR + AGE + SITE + HABITAT | 53.08577 | 0.1764114 |
| Bartonella ~ 1 + AGE + SITE + SEX + HABITAT | 53.18898 | 0.1675382 |

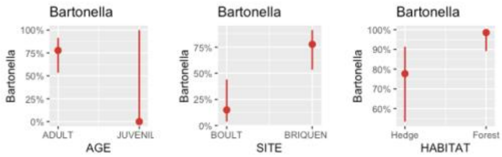

| model | aicc | weights |
|---|---|---|
| VirusCPXV ~ 1 + YEAR | 65.76170 | 0.16012716 |
| VirusCPXV ~ 1 + YEAR + SITE | 66.85951 | 0.09248655 |
| VirusCPXV ~ 1 + YEAR + SEX | 66.97705 | 0.08720769 |
| VirusCPXV ~ 1 + YEAR + Bartonella | 67.19235 | 0.07830731 |

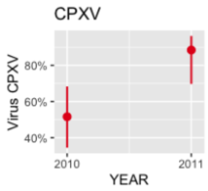

#### *Ap. flavicollis* hosts

| model | aicc | weights |
|---|---|---|
| Bartonella ~ 1 + YEAR | 27.01574 | 0.07494560 |
| Bartonella ~ 1 | 27.23356 | 0.06721206 |
| Bartonella ~ 1 + VirusCPXV | 27.29479 | 0.06518550 |
| Bartonella ~ 1 + SEX + VirusCPXV | 27.54878 | 0.05741128 |
| Bartonella ~ 1 + YEAR + VirusCPXV | 27.59684 | 0.05604811 |
| Bartonella ~ 1 + SEX | 28.75333 | 0.03143636 |
| Bartonella ~ 1 + AGE + VirusCPXV | 28.75937 | 0.03134154 |
| Bartonella ~ 1 + YEAR + SEX + VirusCPXV | 28.76480 | 0.03125666 |
| Bartonella ~ 1 + HABITAT + VirusCPXV | 28.95094 | 0.02847890 |

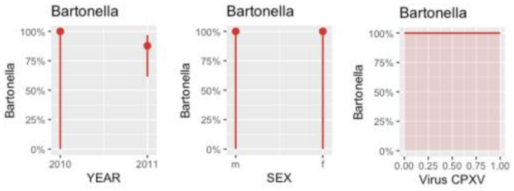

| model | aicc | weights |
|---|---|---|
| VirusCPXV ~ 1 + AGE + SITE | 34.69504 | 0.04799287 |
| VirusCPXV ~ 1 + SITE | 35.27708 | 0.03587477 |
| VirusCPXV ~ 1 + AGE + SITE + SEX + Bartonella | 35.33126 | 0.03491602 |
| VirusCPXV ~ 1 + AGE + SITE + Bartonella | 35.37004 | 0.03424543 |
| VirusCPXV ~ 1 + AGE + SITE + SEX | 35.37328 | 0.03419009 |
| VirusCPXV ~ 1 + SITE + SEX + Bartonella | 35.40481 | 0.03365532 |
| VirusCPXV ~ 1 + AGE + HABITAT + Bartonella | 35.56353 | 0.03108766 |
| VirusCPXV ~ 1 + AGE + SEX + Bartonella | 35.58134 | 0.03081196 |
| VirusCPXV ~ 1 + SITE + SEX | 35.70837 | 0.02891586 |
| VirusCPXV ~ 1 + HABITAT + Bartonella | 35.80688 | 0.02752611 |

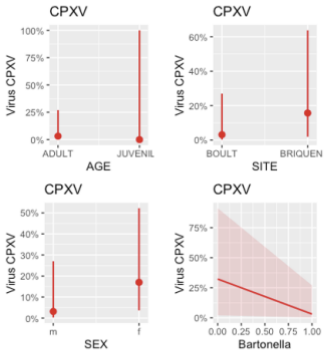

#### *Ap. sylvaticus* hosts

| model | aicc | weights |
|---|---|---|
| Bartonella ~ 1 + YEAR | 57.26551 | 0.06418336 |
| Bartonella ~ 1 + YEAR + AGE | 57.59187 | 0.05451983 |
| Bartonella ~ 1 + YEAR + SEX | 57.62474 | 0.05363114 |
| Bartonella ~ 1 + AGE | 57.84906 | 0.04794082 |
| Bartonella ~ 1 | 57.87548 | 0.04731161 |
| Bartonella ~ 1 + YEAR + HABITAT | 58.06706 | 0.04298992 |
| Bartonella ~ 1 + YEAR + SEX + HABITAT | 58.37090 | 0.03693088 |
| Bartonella ~ 1 + YEAR + AGE + HABITAT | 58.69315 | 0.03143507 |
| Bartonella ~ 1 + SEX | 58.87877 | 0.02864885 |
| Bartonella ~ 1 + YEAR + SITE | 58.99687 | 0.02700607 |
| Bartonella ~ 1 + YEAR + AGE + SEX | 59.06775 | 0.02606583 |
| Bartonella ~ 1 + HABITAT | 59.16057 | 0.02488375 |

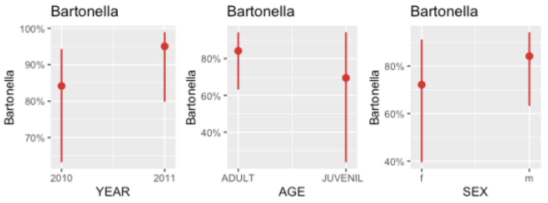

| model | aicc | weights |
|---|---|---|
| VirusCPXV ~ 1 | 50.55493 | 0.08931698 |
| VirusCPXV ~ 1 + AGE + SEX | 50.71568 | 0.08241898 |
| VirusCPXV ~ 1 + SEX | 51.18933 | 0.06503910 |
| VirusCPXV ~ 1 + AGE | 51.86424 | 0.04641107 |
| VirusCPXV ~ 1 + YEAR + AGE + SEX | 51.94742 | 0.04452035 |
| VirusCPXV ~ 1 + HABITAT | 52.33145 | 0.03674232 |
| VirusCPXV ~ 1 + YEAR | 52.40667 | 0.03538617 |

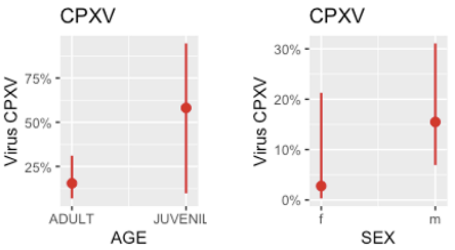

#### *R. norvegicus* hosts

| model | aicc | weights |
|---|---|---|
| Bartonella ~ 1 | 21.22183 | 0.14603691 |
| Bartonella ~ 1 + SITE | 21.38836 | 0.13436963 |
| Bartonella ~ 1 + VirusCPXV | 22.56497 | 0.07461133 |
| Bartonella ~ 1 + SITE + VirusCPXV | 22.84397 | 0.06489619 |
| Bartonella ~ 1 + SITE + SEX | 22.88143 | 0.06369227 |
| Bartonella ~ 1 + SEX | 22.99030 | 0.06031776 |

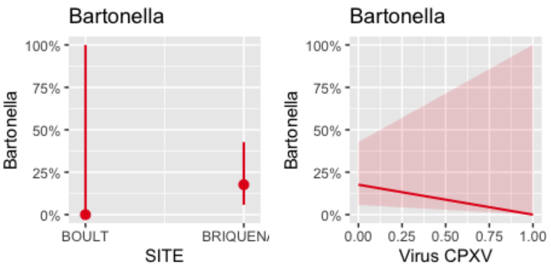

| model | aicc | weights |
|---|---|---|
| VirusCPXV ~ 1 | 25.12036 | 0.15829184 |
| VirusCPXV ~ 1 + AGE | 25.34213 | 0.14167780 |
| VirusCPXV ~ 1 + Bartonella | 26.46349 | 0.08087246 |
| VirusCPXV ~ 1 + YEAR | 26.80155 | 0.06829559 |
| VirusCPXV ~ 1 + AGE + Bartonella | 27.01610 | 0.06134857 |

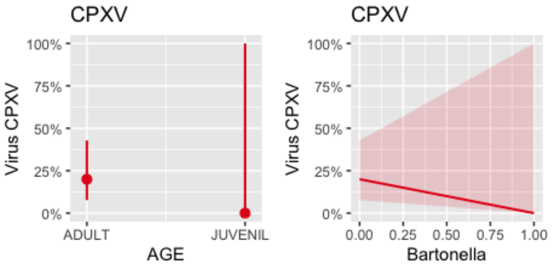

Figure 15: Results of Akaike Information Criterion (AIC)-based model selection analysis of associations between occurrence of *Mycoplasma haemomuris* (MH) and *Mycoplasma coccoides* (MC). Plotted here are the coefficients of marginal effects for each factor level remaining in the best model after model selection, as well as a table showing the AICc weights for the top-ranking models.

| model | aicc | weights |
|---|---|---|
| MH ~ 1 + YEAR + SITE + HostSPP + MC | 182.0843 | 0.19313870 |
| MH ~ 1 + YEAR + SITE + HABITAT + HostSPP + MC | 182.9863 | 0.12302929 |
| MH ~ 1 + SITE + HostSPP + MC | 183.8526 | 0.07977764 |
| MH ~ 1 + SITE + HABITAT + HostSPP + MC | 183.8819 | 0.07861901 |

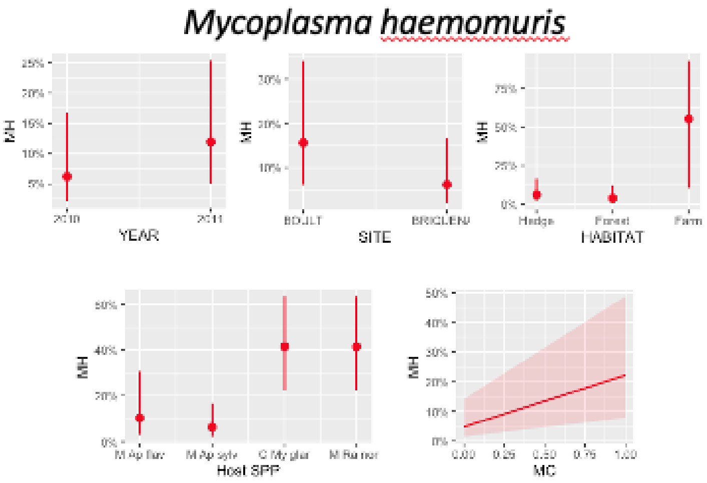

| model | aicc | weights |
|---|---|---|
| MC ~ 1 + SEX + HostSPP + MH | 142.1091 | 0.11000460 |
| MC ~ 1 + HostSPP + MH | 142.3187 | 0.09905957 |
| MC ~ 1 + YEAR + HostSPP + MH | 143.3310 | 0.05971368 |
| MC ~ 1 + SITE + SEX + HostSPP + MH | 143.4349 | 0.05668966 |
| MC ~ 1 + YEAR + SEX + HostSPP + MH | 143.5415 | 0.05374968 |
| MC ~ 1 + SITE + HostSPP + MH | 143.5836 | 0.05262800 |
| MC ~ 1 + AGE + HostSPP + MH | 143.9068 | 0.04477509 |
| MC ~ 1 + SEX + HABITAT + HostSPP + MH | 143.9531 | 0.04375063 |
| MC ~ 1 + SEX + AGE + HostSPP + MH | 144.0519 | 0.04164170 |
| MC ~ 1 + HABITAT + HostSPP + MH | 144.0631 | 0.04141020 |

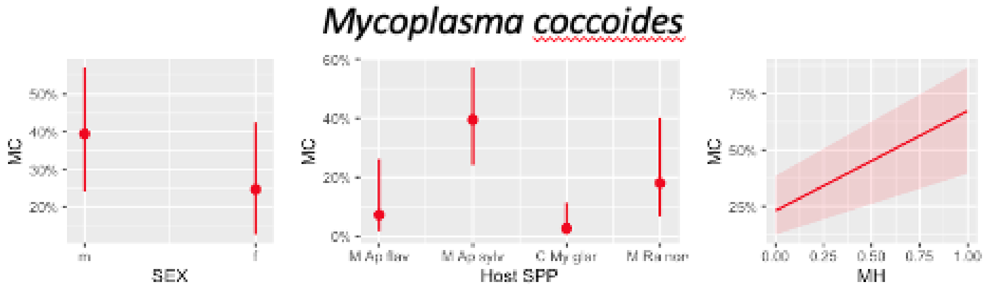

Figure 16: Results of Akaike Information Criterion (AIC)-based model selection analysis of associations between occurrence of *Bartonella* spp. and hemotropic *Mycoplasma* spp. infections across rodent species. Plotted here are the coefficients of marginal effects for each factor level remaining in the best model after model selection, as well as a table showing the AICc weights for the top-ranking models.

Global Model for *Bartonella* spp.

| model | aicc | weights |
|---|---|---|
| Bartonella ~ 1 + YEAR + HABITAT + HMyco | 280.9917 | 0.16167656 |
| Bartonella ~ 1 + YEAR + HABITAT | 282.4646 | 0.07741243 |
| Bartonella ~ 1 + YEAR + HABITAT + HostSPP + HMyco | 282.8027 | 0.06537150 |
| Bartonella ~ 1 + YEAR + HABITAT + SEX + HMyco | 282.9414 | 0.06099185 |

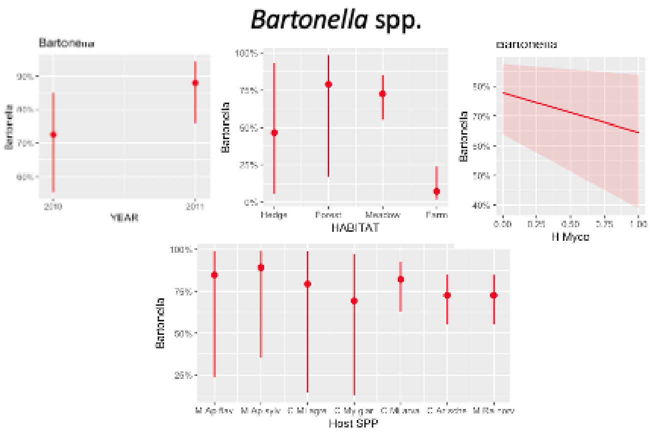

Global Model for hemotropic *Mycoplasma* spp

| model | aicc | weights |
|---|---|---|
| HMyco ~ 1 + YEAR + SITE + HostSPP + Bartonella | 271.2146 | 0.08379287 |
| HMyco ~ 1 + HABITAT + SITE + HostSPP | 271.4399 | 0.07486482 |
| HMyco ~ 1 + YEAR + HABITAT + SITE + HostSPP | 272.0079 | 0.05635645 |
| HMyco ~ 1 + YEAR + SITE + SEX + HostSPP + Bartonella | 272.1142 | 0.05343822 |
| HMyco ~ 1 + YEAR + HABITAT + SITE + HostSPP + Bartonella | 272.1852 | 0.05157477 |
| HMyco ~ 1 + HABITAT + SITE + SEX + HostSPP | 272.2139 | 0.05084189 |
| HMyco ~ 1 + HABITAT + SITE + HostSPP + Bartonella | 272.2634 | 0.04959914 |
| HMyco ~ 1 + YEAR + SITE + HostSPP | 272.3821 | 0.04674085 |
| HMyco ~ 1 + SITE + HostSPP + Bartonella | 272.4397 | 0.04541321 |
| HMyco ~ 1 + SITE + HostSPP | 272.7883 | 0.03814905 |

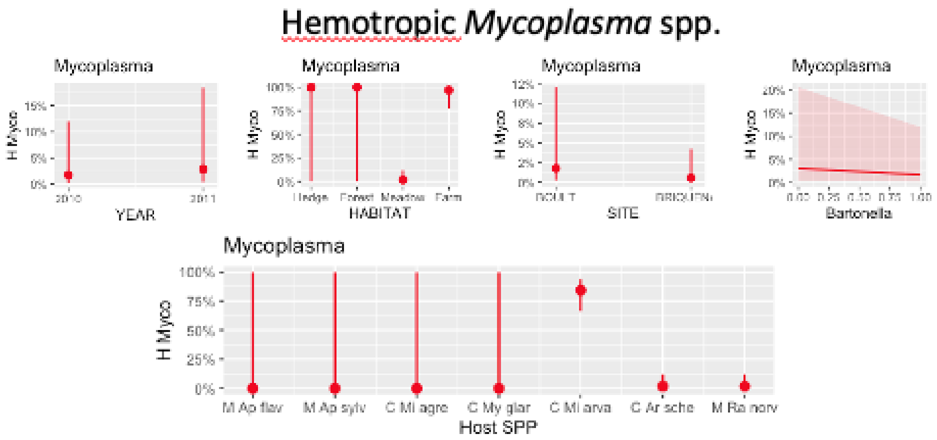

Models for *Mi. agrestis*

| model | aicc | weights |
|---|---|---|
| Bartonella ~ 1 | 11.66403 | 0.2518089 |
| Bartonella ~ 1 + AGE | 12.14163 | 0.1983178 |
| Bartonella ~ 1 + YEAR | 13.13012 | 0.1209802 |

| model | aicc | weights |
|---|---|---|
| HMyco ~ 1 | 13.75702 | 0.1962635 |
| HMyco ~ 1 + SEX | 14.03817 | 0.1705253 |
| HMyco ~ 1 + HABITAT | 14.03817 | 0.1705253 |

Models for *Mi. arvalis* (3 from hedgerows excluded b/c all infected with HM)

| model | aicc | weights |
|---|---|---|
| Bartonella ~ 1 + SITE | 35.52989 | 0.12017117 |
| Bartonella ~ 1 + SITE + AGE | 35.75816 | 0.10720871 |
| Bartonella ~ 1 + YEAR + SITE | 36.57428 | 0.07128755 |
| Bartonella ~ 1 | 37.14626 | 0.05355600 |
| Bartonella ~ 1 + AGE | 37.14810 | 0.05350681 |
| Bartonella ~ 1 + AGE + HMyco | 37.19535 | 0.05225752 |
| Bartonella ~ 1 + YEAR + SITE + AGE | 37.32500 | 0.04897732 |
| Bartonella ~ 1 + HMyco | 37.50453 | 0.04477252 |

| model | aicc | weights |
|---|---|---|
| HMyco ~ 1 + SITE | 30.83130 | 0.16865598 |
| HMyco ~ 1 + YEAR + SITE | 31.28056 | 0.13472432 |
| HMyco ~ 1 + SITE + SEX | 31.90781 | 0.09845569 |
| HMyco ~ 1 + YEAR + SITE + SEX | 32.55396 | 0.07127385 |
| HMyco ~ 1 + SITE + AGE | 32.73952 | 0.06495857 |

Models for *My. glareolus*

| model | aicc | weights |
|---|---|---|
| Bartonella ~ 1 + YEAR + HABITAT + SITE + AGE + HMyco | 51.44478 | 0.21037388 |
| Bartonella ~ 1 + HABITAT + SITE + AGE + HMyco | 51.73181 | 0.18224805 |
| Bartonella ~ 1 + HABITAT + SITE + AGE | 52.11549 | 0.15043445 |
| Bartonella ~ 1 + YEAR + HABITAT + SITE + AGE | 53.08577 | 0.09260939 |
| Bartonella ~ 1 + HABITAT + SITE + AGE + SEX | 53.18898 | 0.08795132 |
| Bartonella ~ 1 + HABITAT + SITE + AGE + SEX + HMyco | 53.31144 | 0.08272748 |

| model | aicc | weights |
|---|---|---|
| HMyco ~ 1 + YEAR + SITE + Bartonella | 72.70929 | 0.1843991 |
| HMyco ~ 1 + YEAR + Bartonella | 73.68877 | 0.1129969 |
| HMyco ~ 1 + YEAR + SITE + SEX + Bartonella | 74.67863 | 0.0688845 |

Models for *Ap. flavicollis*

| model | aicc | weights |
|---|---|---|
| Bartonella ~ 1 + YEAR | 27.01574 | 0.09595236 |
| Bartonella ~ 1 | 27.23356 | 0.08605116 |
| Bartonella ~ 1 + HMyco | 27.77084 | 0.06577894 |
| Bartonella ~ 1 + YEAR + HMyco | 28.12909 | 0.05499127 |
| Bartonella ~ 1 + SEX | 28.75333 | 0.04024777 |

| model | aicc | weights |
|---|---|---|
| HMyco ~ 1 | 40.25649 | 0.09298047 |
| HMyco ~ 1 + Bartonella | 40.79378 | 0.07107582 |
| HMyco ~ 1 + HABITAT | 41.14676 | 0.05957608 |
| HMyco ~ 1 + YEAR | 41.54577 | 0.04880097 |
| HMyco ~ 1 + HABITAT + Bartonella | 41.87876 | 0.04131620 |
| HMyco ~ 1 + SITE | 41.92840 | 0.04030338 |
| HMyco ~ 1 + AGE | 42.02595 | 0.03838475 |

Models for *Ap. sylvaticus*

| model | aicc | weights |
|---|---|---|
| Bartonella ~ 1 + YEAR | 57.26551 | 0.06303068 |
| Bartonella ~ 1 + YEAR + AGE | 57.59187 | 0.05354070 |
| Bartonella ~ 1 + YEAR + SEX | 57.62474 | 0.05266797 |
| Bartonella ~ 1 + AGE | 57.84906 | 0.04707984 |
| Bartonella ~ 1 | 57.87548 | 0.04646193 |
| Bartonella ~ 1 + YEAR + HABITAT | 58.06706 | 0.04221786 |
| Bartonella ~ 1 + YEAR + SEX + HABITAT | 58.37090 | 0.03626764 |
| Bartonella ~ 1 + YEAR + AGE + HABITAT | 58.69315 | 0.03087052 |
| Bartonella ~ 1 + SEX | 58.87877 | 0.02813434 |
| Bartonella ~ 1 + YEAR + SITE | 58.99687 | 0.02652106 |

| model | aicc | weights |
|---|---|---|
| HMyco ~ 1 | 86.53686 | 0.12549877 |
| HMyco ~ 1 + SEX | 87.84557 | 0.06523126 |
| HMyco ~ 1 + YEAR | 87.95622 | 0.06172024 |
| HMyco ~ 1 + SITE | 88.08997 | 0.05772771 |
| HMyco ~ 1 + HABITAT | 88.51300 | 0.04672237 |

**Figure S17. Benjamini-Hochberg correction for multiple tests.** Comparison of raw p-values from hypothesis tests (black bars) and p-values adjusted for false discovery using the Benjamini-Hochberg correction (red bars). P-values (N=77) were sorted lowest to highest, and only adjusted p-values below 0.1 are illustrated here.

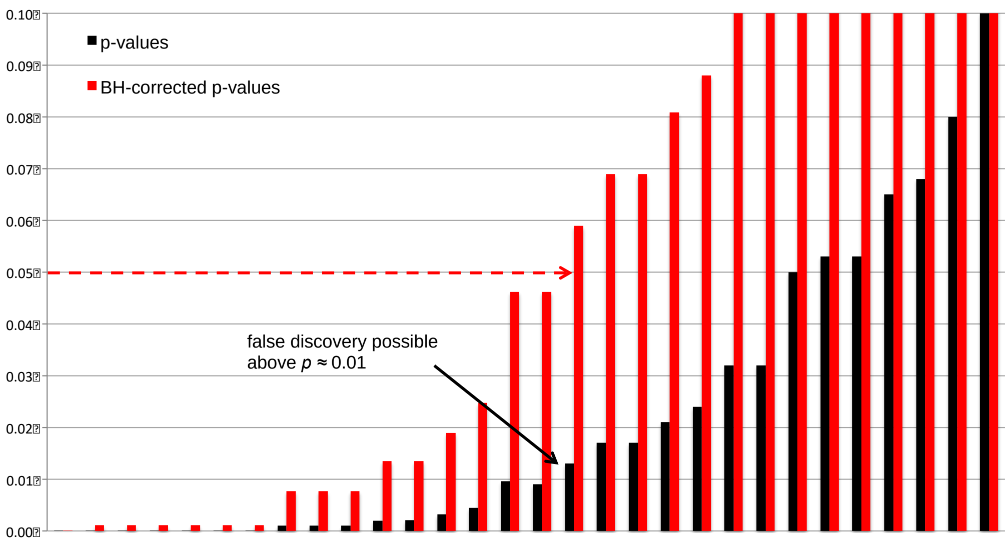

### Appendices

#### Appendix 1: Description of rare host species, their pathogens, and other under-detected pathogens.

##### Rare Host Species and their pathogens

Five *Microtus subterraneus* (European pine vole) and one each of three additional host species (*Ondatra zibethicus* (muskrat), *Myocastor coypus* (the coypu), and an unidentified *Apodemus* species) were also found in these communities, but excluded from analyses due to their rarity. There was no evidence for past exposure to hantavirus, CPXV, LCMV or TBE virus (with just one animal missing data). The six animals from which there were 16S data showed that each of these animals harbored one bacterial OTU (see table below). These included both common OTUs (*Bartonella* sp. and Myco2) and two very rare species: *Brevinema* sp., and a likely pulmonary *Mycoplasma* OTU 04125, discussed below.

| Host species (each individual) | Year | Site | Habitat | Age | Sex | Pathogen exposures |
|---|---|---|---|---|---|---|
| ***Mi. subterraneus*** **(Cricetidae)** | 2010 | Boult | Hedge | n/a | Male | No viruses; *Brevinema* sp., |
| ***Mi. subterraneus*** | 2010 | Boult | Hedge | Adult | Female | No viruses; 16S n/a |
| ***Mi. subterraneus*** | 2010 | Boult | Forest | Adult | Female | No viruses; *Bartonella* sp. |
| ***Mi. subterraneus*** | 2011 | Boult | Forest | n/a | Male | No viruses; *Bartonella* sp. |
| ***Mi. subterraneus*** | 2011 | Boult | Forest | n/a | Male | No viruses; *Bartonella* sp. |
| ***On. zibethicus*** **(Cricetidae)** | 2011 | Briquenay | Forest | 2010 | Male | No viruses; Myco4125, |
| ***Myo. coypus*** **(Echimyidae)** | 2011 | Boult | Forest | 2010 | Male | Virus n/a; Myco2, |
| **Unidentified *Apodemus* spp. (Muridae)** | 2011 | Boult | Hedge | 2010 | Female | No viruses; 16S n/a |

##### Under-detected Pathogens

Three *Mycoplasma* OTUs with fewer than 500 reads in the dataset nevertheless likely represented true infections worth noting. All 255 sequences of OTU 00771 were found in replicates of a single *Ra. norvegicus* individual, and displayed 100% sequence identity to *M. pulmonis*, another pneumotropic *Mycoplasma* species of both wild and laboratory rats that has been shown to asymptomatically infect humans in regular contact with infected animals (Piasecki et al., 2017). Another *Ra. norvegicus* individual harboured 51 out of 59 reads (86%) for OTU 00316 that displayed 4% sequence divergence from the hemotropic Myco2. Finally, all 164 sequences of OTU 04125 were found in the single *On. zibethica* (non-focal host species) individual, and its sequence matches most closely with that of Candidatus *M. ravipulmonis* (Accession AF001173; 91% identity). Curiously, the next closest GenBank hit for Myco10 was an uncultured intestinal *Mycoplasma* from small abalone *Haliotis diversicolor* in inland ponds (Accession HQ393436; 89% identity; (Huang et al.,

2010)) and *Myco. orale* from both squid (*Doryteuthis opalescens*; Accession JQ192010; 89% identity; (Bik et al., 2016)) and throat swabs of human subjects (from several unpublished direct submission datasets; 89% identity).

Four additional OTUs were recognized as pathogenic despite their failure to meet inclusion criteria (below 500 reads in the dataset), but are also worth noting. Three belong to the genus *Borrelia* (Borrelia1/OTU 00318 in one *My. glareolus* and one *Ap. sylvaticus*, Borrelia2/OTU 00514 in two *Ar. scherman* individuals, Borrelia3/OTU 00071 in a second *My. glareolus*), and the fourth belongs to the genus *Leptospira* (Leptospira1/OTU 01015 in one *Ap. sylvaticus* and one *Ra. norvegicus*), both of which have close sequence identity to pathogens, but which are not likely to be systematically detectable from splenic tissue of infected animals.

Remarkably, the 16S data also included three unclassified OTUs (00056 (Sarco1), 00191(Sarco2) and 00254(Sarco3)) meeting the inclusion criteria for consideration of prevalence whose closest GenBank sequence matches were not bacterial, but rather grouped with a diversity of plastid sequences from pathogenic coccidian species in the Sarcocystidae family. In addition to *T. gondii* (U87145, 95% identity to Sarco1, 94% identity to Sarco3, 90% identity to Sarco2), these included *Sarcocystis muris* (AF255924, 98% identity to OTU Sarco3, 95% identity to Sarco1, 89% identity to OTU Sarco2), *Neospora caninum* (AF204319, 96% identity to Sarco1, 95% identity to Sarco3, 92% identity to Sarco2), and *Hyaloklossia lieberkuehni* (AF297120, 97% identity to Sarco1, 95% identity to Sarco3, 90% identity to Sarco2). Sarco1 was found in 2 *Ar. scherman,* 1 *Mi. agrestis,* 1 *Mi. subterraneus* and 7 *My. glareolus* individuals; Sarco2 was found in 1 *My. glareolus* and Sarco3 was found in 3 *My. glareolus.* There was also one co-infection (Sarco1and Sarco3 in one *My. glareolus*). However, the reliability of detection by this method is unclear, so these data were excluded from the study.

Bik, E. M., Costello, E. K., Switzer, A. D., Callahan, B. J., Holmes, S. P., Wells, R. S., Carlin, K. P., Jensen, E. D., Venn-Watson, S., & Relman, D. A. (2016). Marine mammals harbor unique microbiotas shaped by and yet distinct from the sea. *Nature Communications*, *7*, 10516. https://doi.org/10.1038/ncomms10516

Huang, Z. Bin, Guo, F., Zhao, J., Li, W. D., & Ke, C. H. (2010). Molecular analysis of the intestinal bacterial flora in cage-cultured adult small abalone, Haliotis diversicolor. *Aquaculture Research*, *41*(11), e760–e769. https://doi.org/10.1111/j.1365-2109.2010.02577.x

Piasecki, T., Chrzastek, K., & Kasprzykowska, U. (2017). *Mycoplasma pulmonis* of Rodents as a Possible Human Pathogen. *Vector-Borne and Zoonotic Diseases*, *17*(7), 475–477. https://doi.org/10.1089/vbz.2016.2104

#### Appendix 2: Scripts and data for statistical analyses

Scripts and data for statistical analyses are provided in Statistical Analysis Scripts and Data File.zip in Zenodo (https://doi.org/10.5281/zenodo.7092812)

#### Appendix 3: References for Table 1

Literature Referenced in Table 1 of Main Text

Alabí, A. S., Monti, G., Otth, C., Sepulveda-García, P., Sánchez-Hidalgo, M., de Mello, V. V. C., Machado, R. Z., André, M. R., Bittencourt, P., & Müller, A. (2020). Molecular survey and genetic diversity of hemoplasmas in rodents from Chile. *Microorganisms*, *8*(10), 1–16. https://doi.org/10.3390/microorganisms8101493

Andersson, M., & Råberg, L. (2011). Wild rodents and novel human pathogen Candidatus *Neoehrlichia mikurensis*, southern Sweden. *Emerging Infectious Diseases*, *17*(9), 1716–1718. https://doi.org/10.3201/eid1709.101058

Bekele, A. Z., Koike, S., & Kobayashi, Y. (2011). Phylogenetic diversity and dietary association of rumen *Treponema* revealed using group-specific 16S rRNA gene-based analysis. In *FEMS Microbiology Letters* (Vol. 316, Issue 1, pp. 51–60). https://doi.org/10.1111/j.1574-6968.2010.02191.x

Bernard, K. (2012). The genus *Corynebacterium* and other medically relevant coryneform-like bacteria. In *Journal of Clinical Microbiology* (Vol. 50, Issue 10, pp. 3152–3158). https://doi.org/10.1128/JCM.00796-12

Bunikis, J., & Barbour, A. G. (2005). Third *Borrelia* Species in White-footed Mice. *Emerging Infectious Diseases*, *11*(7), 1150–1151. www.cdc.gov/eid

Calarco, L., & Ellis, J. (2020). Contribution of introns to the species diversity associated with the apicomplexan parasite, Neospora caninum. *Parasitology Research*, *119*(2), 431–445. https://doi.org/10.1007/s00436-019-06561-x

Cisak, E., Wójcik-Fatla, A., Zając, V., Sawczyn, A., Sroka, J., & Dutkiewicz, J. (2015). Spiroplasma - An emerging arthropod-borne pathogen? In *Annals of Agricultural and Environmental Medicine* (Vol. 22, Issue 4, pp. 589–593). Institute of Agricultural Medicine. https://doi.org/10.5604/12321966.1185758

Conrado, F. D. O., do Nascimento, N. C., dos Santos, A. P., Zimpel, C. K., Messick, J. B., & Biondo, A. W. (2015). Occurrence and identification of hemotropic mycoplasmas (Hemoplasmas) in free ranging and laboratory rats (Rattus norvegicus) from two Brazilian zoos. *BMC Veterinary Research*, *11*(1), 286. https://doi.org/10.1186/s12917-015-0601-8

Defosse, D. L., Johnson, R. C., Paster, B. J., Dewhirst, F. E., & Fraser, G. J. (1995). *Brevinema andersonii* gen. nov., sp. nov., an infectious spirochete isolated from the short-tailed shrew (*Blarina brevicauda*) and the white-footed mouse (*Peromyscus leucopus*). *International Journal of Systematic Bacteriology*, *45*(1), 78–84. https://doi.org/10.1099/00207713-45-1-78

Deng, H. K., Le Rhun, D., Buffet, J. P. R., Cotte, V., Read, A., Birtles, R. J., & Vayssier-Taussat, M. (2012). Strategies of exploitation of mammalian reservoirs by Bartonella species. *Veterinary Research*, *43*. https://doi.org/10.1186/1297-9716-43-15

Duron, O., Noël, V., McCoy, K. D., Bonazzi, M., Sidi-Boumedine, K., Morel, O., Vavre, F., Zenner, L., Jourdain, E., Durand, P., Arnathau, C., Renaud, F., Trape, J. F., Biguezoton, A. S., Cremaschi, J., Dietrich, M., Léger, E., Appelgren, A., Dupraz, M., … Chevillon, C. (2015). The Recent Evolution of a Maternally-Inherited Endosymbiont of Ticks Led to the Emergence of the Q Fever Pathogen, Coxiella burnetii. *PLoS Pathogens*, *11*(5). https://doi.org/10.1371/journal.ppat.1004892

Galan, M., Razzauti, M., Bard, E., Bernard, M., Brouat, C., Charbonnel, N., Dehne-Garcia, A., Loiseau, A., Tatard, C., Tamisier, L., Vayssier-Taussat, M., Vignes, H., & Cosson, J.-F. (2016). 16S rRNA Amplicon Sequencing for Epidemiological Surveys of Bacteria in Wildlife. *MSystems*, *1*(4), e00032-16. https://doi.org/10.1128/mSystems.00032-16

Gonçalves, L. R., Roque, A. L. R., Matos, C. A., Fernandes, S. de J., Olmos, I. D. F., Machado, R. Z., & André, M. R. (2015). Diversity and molecular characterization of novel hemoplasmas infecting wild rodents from different Brazilian biomes. *Comparative Immunology, Microbiology and Infectious Diseases*, *43*, 50–56. https://doi.org/10.1016/j.cimid.2015.10.006

Goto, K., Ohashi, H., Ebukuro, S., Itoh, K., Tohma, Y., Takakura, A., Wakana, S., Ito, M., & Itoh, T. (n.d.). *Isolation and Characterization of Helicobacter Species from the Stomach of the House Musk Shrew (*Suncus murinus*) with Chronic Gastritis*.

Kämpfer, P., Falsen, E., Frischmann, A., & Busse, H. J. (2012). *Dietzia aurantiaca* sp. nov., isolated from a human clinical specimen. *International Journal of Systematic and Evolutionary Microbiology*, *62*(PART 3), 484–488. https://doi.org/10.1099/ijs.0.032557-0

Kim, W., Song, M.-O., Song, W., Kim, K.-J., Chung, S.-I., Choi, S., & Park, Y.-H. (2003). Comparison of 16S rDNA analysis and rep-PCR genomic fingerprinting for molecular identification of *Yersinia pseudotuberculosis*. In *Antonie van Leeuwenhoek* (Vol. 83).

Krause, P. J., Fish, D., Narasimhan, S., & Barbour, A. G. (2015). *Borrelia miyamotoi* infection in nature and in humans. In *Clinical Microbiology and Infection* (Vol. 21, Issue 7, pp. 631–639). Elsevier B.V. https://doi.org/10.1016/j.cmi.2015.02.006

Melito, P. L., Munro, C., Chipman, P. R., Woodward, D. L., Booth, T. F., & Rodgers, F. G. (2001). *Helicobacter winghamensis* sp. nov., a novel *Helicobacter* sp. isolated from patients with gastroenteritis. *Journal of Clinical Microbiology*, *39*(7), 2412–2417. https://doi.org/10.1128/JCM.39.7.2412-2417.2001

Nicklas, W., Bisgaard, M., Aalbæk, B., Kuhnert, P., & Christensen, H. (2015). Reclassification of *Actinobacillus muris* as *Muribacter muris* gen. Nov., comb. nov. *International Journal of Systematic and Evolutionary Microbiology*, *65*(10), 3344–3351. https://doi.org/10.1099/ijsem.0.000417

Orosz, F. (2015). Two recently sequenced vertebrate genomes are contaminated with apicomplexan species of the Sarcocystidae family. *International Journal for Parasitology*, *45*(13), 871–878. https://doi.org/10.1016/j.ijpara.2015.07.002

Paris, D. H., Shelite, T. R., Day, N. P., & Walker, D. H. (2013). Review article: Unresolved problems related to scrub typhus: A seriously neglected life-threatening disease. In *American Journal of Tropical Medicine and Hygiene* (Vol. 89, Issue 2, pp. 301–307). https://doi.org/10.4269/ajtmh.13-0064

Patterson, M. M., Schrenzel, M. D., Feng, † Y, Xu, S., Dewhirst, F. E., Paster, B. J., Thibodeau, S. A., Versalovic, J., & Fox, J. G. (2000). Species Cultured from Gastrointestinal Tissues of Syrian Hamsters. In *JOURNAL OF CLINICAL MICROBIOLOGY* (Vol. 38, Issue 10).

Pettersson, B., Tully, J. G., Bolske, G., & Johansson, K.-E. (2000). Updated phylogenetic description of the *Mycoplasma hominis* cluster (Weisburg et al. 1989) based on 16S rDNA sequences. In *International Journal of Systematic and Evolutionary Microbiology* (Vol. 50).

Piasecki, T., Chrzastek, K., & Kasprzykowska, U. (2017). *Mycoplasma pulmonis* of Rodents as a Possible Human Pathogen. *Vector-Borne and Zoonotic Diseases*, *17*(7), 475–477. https://doi.org/10.1089/vbz.2016.2104

Schüler, W., Bunikis, I., Weber-Lehman, J., Comstedt, P., Kutschan-Bunikis, S., Stanek, G., Huber, J., Meinke, A., Bergstroäm, S., & Lundberg, U. (2015). Complete genome sequence of *Borrelia afzelii* K78 and comparative genome analysis. *PLoS ONE*, *10*(3). https://doi.org/10.1371/journal.pone.0120548

Vandamme, P., Vancanneyt, M., Pot, B., Mels, L., Hoste, B., Dewettinck, D., Vlaes, L., Van Den Borre, C., Higgins, R., Hommez, J., Kersters, K., Butzler, J.-P., & Goossens, H. (1992). Polyphasic Taxonomic Study of the Emended Genus *Arcobacter* with *Arcobacter butzleri* comb. nov. and *Arcobacter skirrowii* sp. nov., an Aerotolerant Bacterium Isolated from Veterinary Specimens. *International Journal of Systematic Bacteriology*, *42*(3), 344–356. https://doi.org/10.1099/00207713-42-3-344

Vincent, A. T., Schiettekatte, O., Goarant, C., Neela, V. K., Bernet, E., Thibeaux, R., Ismail, N., Khalid, M. K. N. M., Amran, F., Masuzawa, T., Nakao, R., Korba, A. A., Bourhy, P., Veyrier, F. J., & Picardeau, M. (2019). Revisiting the taxonomy and evolution of pathogenicity of the genus Leptospira through the prism of genomics. *PLoS Neglected Tropical Diseases*, *13*(5). https://doi.org/10.1371/journal.pntd.0007270

Yamamoto, S., Morita1, C., & Tsuchiya2, K. (1992). Isolation of spotted fever group Rickettsia from *Apodemus speciosus* in an endemic area in Japan. *Jpn. J. Med. Sci. Biol*, *45*, 81–86.
